# A global view of aging and Alzheimer’s pathogenesis-associated cell population dynamics and molecular signatures in the human and mouse brains

**DOI:** 10.1101/2022.09.28.509825

**Authors:** Andras Sziraki, Ziyu Lu, Jasper Lee, Gabor Banyai, Sonya Anderson, Abdulraouf Abdulraouf, Eli Metzner, Andrew Liao, Jason Banfelder, Alexander Epstein, Chloe Schaefer, Zihan Xu, Zehao Zhang, Li Gan, Peter T. Nelson, Wei Zhou, Junyue Cao

**Affiliations:** Laboratory of Single Cell Genomics and Population Dynamics, The Rockefeller University, New York, NY, USA; The David Rockefeller Graduate Program in Bioscience, The Rockefeller University, New York, NY, USA; Department of Pathology and Sanders-Brown Center on Aging, University of Kentucky, Lexington, KY, USA; The Tri-Institutional M.D-Ph.D Program, New York, NY, USA; The Tri-Institutional Ph.D. Program in Computational Biology & Medicine, New York, NY, USA; High Performance Computing Resource Center, The Rockefeller University, New York, NY, USA; Helen & Robert Appel Alzheimer’s Disease Research Institute, Weill Cornell Medicine, New York, NY, USA

**Author notes:** Correspondence (W.Z.), (J.C.). These authors contributed equally (Andras Sziraki, Ziyu Lu, Jasper Lee). Senior author (Wei Zhou, Junyue Cao).

## Abstract

Conventional single-cell genomics approaches are limited by throughput and thus may have failed to capture aspects of the molecular signatures and dynamics of rare cell types associated with aging and diseases. Here, we developed *EasySci*, an extensively improved single-cell combinatorial indexing strategy, for investigating the age-dependent dynamics of transcription and chromatin accessibility across diverse brain cell types. We profiled ∼1.5 million single-cell transcriptomes and ∼400,000 single-cell chromatin accessibility profiles across mouse brains spanning different ages, genotypes, and both sexes. With a novel computational framework designed for characterizing cellular subtypes based on the expression of both genes and exons, we identified > 300 cell subtypes and deciphered their underlying molecular programs and spatial locations especially for rare cell types (*e.g.,* pinealocytes, tanycytes). Leveraging these data, we generated a global readout of age-dependent changes at cell subtype resolution, providing insights into cell types that expand (*e.g.,* rare astrocytes and vascular leptomeningeal cells in the olfactory bulb, reactive microglia, and oligodendrocytes) or are depleted (*e.g.,* neuronal progenitors, neuroblasts, committed oligodendrocyte precursors) as age progresses. Furthermore, we explored cell-type-specific responses to genetic perturbations associated with Alzheimer’s disease (AD) and identified rare cell types depleted (*e.g., mt-Cytb*+*, mt-Rnr2*+ choroid plexus epithelial cells) or enriched (*e.g., Col25a1*+, *Ndrg1*+ interbrain and midbrain neurons) in both AD models. Key findings are consistent between males and females, validated across the transcriptome, chromatin accessibility, and spatial analyses. Finally, we profiled a total of 118,240 single-nuclei transcriptomes from twenty-four post-mortem human brain samples derived from control and AD patients, revealing highly cell-type-specific and region-specific gene expression changes associated with AD pathogenesis. Critical AD-associated gene signatures were validated in both human and mice. In summary, these data comprise a rich resource for exploring cell-type-specific dynamics and the underlying molecular mechanisms in normal and pathological mammalian aging.

## Introduction

The mammalian brain is a remarkably complex system made up of millions of highly heterogeneous cells, comprising a myriad of different cell types and subtypes^1,2^. Progressive changes in brain cell populations, which occur during the normal aging process, may contribute to the functional decline of the entire organ and increased risks for neurodegenerative diseases such as Alzheimer’s disease (AD)^3,4^. While the recent advances in single-cell genomics have created unprecedented opportunities to explore the cell-type-specific dynamics across the entire mammalian brain in aging and AD models^5–8^, most prior studies relied on a relatively shallow sampling of the brain cell populations, possibly resulting in poor sensitivity to investigate the dynamics of cell types during aging, particularly with respect to rare aging or AD-associated cell types. While providing proof of key concepts, the prior studies were also technically limited in several ways, including failing to recover isoform-level gene expression patterns and the associated chromatin landscape that regulates cell-type-specific alterations across aging stages, especially in rare cell types.

To address these challenges and achieve a systematic view of cell-type-specific dynamics in aging and AD, we extensively optimized single-cell RNA sequencing by combinatorial indexing, a methodological framework involving split-pool barcoding of cells or nuclei for single-cell transcriptome profiling^9^. While the original method has been widely used to study embryonic and fetal tissues^10,11^, it remains restricted to gene quantification proximal to the 3’ end (*i.e.,* full-length transcript isoform information is lost) and is limited in terms of efficiency and cell recovery (up to 95% cell loss rate)^11^, which pose a challenge when dealing with aged tissues. To overcome these constraints, we have undertaken a comprehensive series of optimizations focusing on critical aspects such as cell lysis, fixation, preservation, enzyme efficiency, oligonucleotide design, and purification methodologies (representative examples are provided in **Figure S1 and S2; Methods)**. Several test conditions were inspired by optimizations described in recently developed or optimized single-cell techniques^12,13^.

The major improvements of the resulting method, *EasySci-RNA* (**Figure 1A**), include (i) *one million* single-cell transcriptomes prepared at a library preparation cost of around $700, less than 1/100 the cost of the commercial platforms ^14^ (**Figure 1B and 1C)**. Of note, this cost mainly includes the reagents cost for scRNA-seq library preparation and does not include the cost of personnel or sequencing; (ii) nuclei are deposited to different wells for reverse transcription with indexed oligo-dT and random hexamer primers (*i.e.,* different molecular barcodes to separate reads primed by two types of primers and across different wells), thus recovering cell-type-specific gene expression with full gene body coverage (**Figure 1D**); (iii) chemically modified ligation adapter oligos were included in the ligation reaction to prevent the formation of primer-dimers and increase the detection efficiency (**Figure S2**); (iv) cell recovery rate, as well as the number of transcripts detected per cell, were significantly improved through optimized nuclei storage and enzymatic reactions (**Figure S2**). The optimized technique yields significantly higher signals per nucleus compared with the original sci-RNA-seq3 protocols as well as the popular commercial platform (*i.e.,* 10x Genomics, SPLiT-seq) (**Figure 1E; Figure S2N-O**); (v) An extensively improved single-cell data processing pipeline was developed for both gene counting and exonic counting utilizing paired-end single-cell RNA sequencing data (**Methods**).

**Figure 1.**
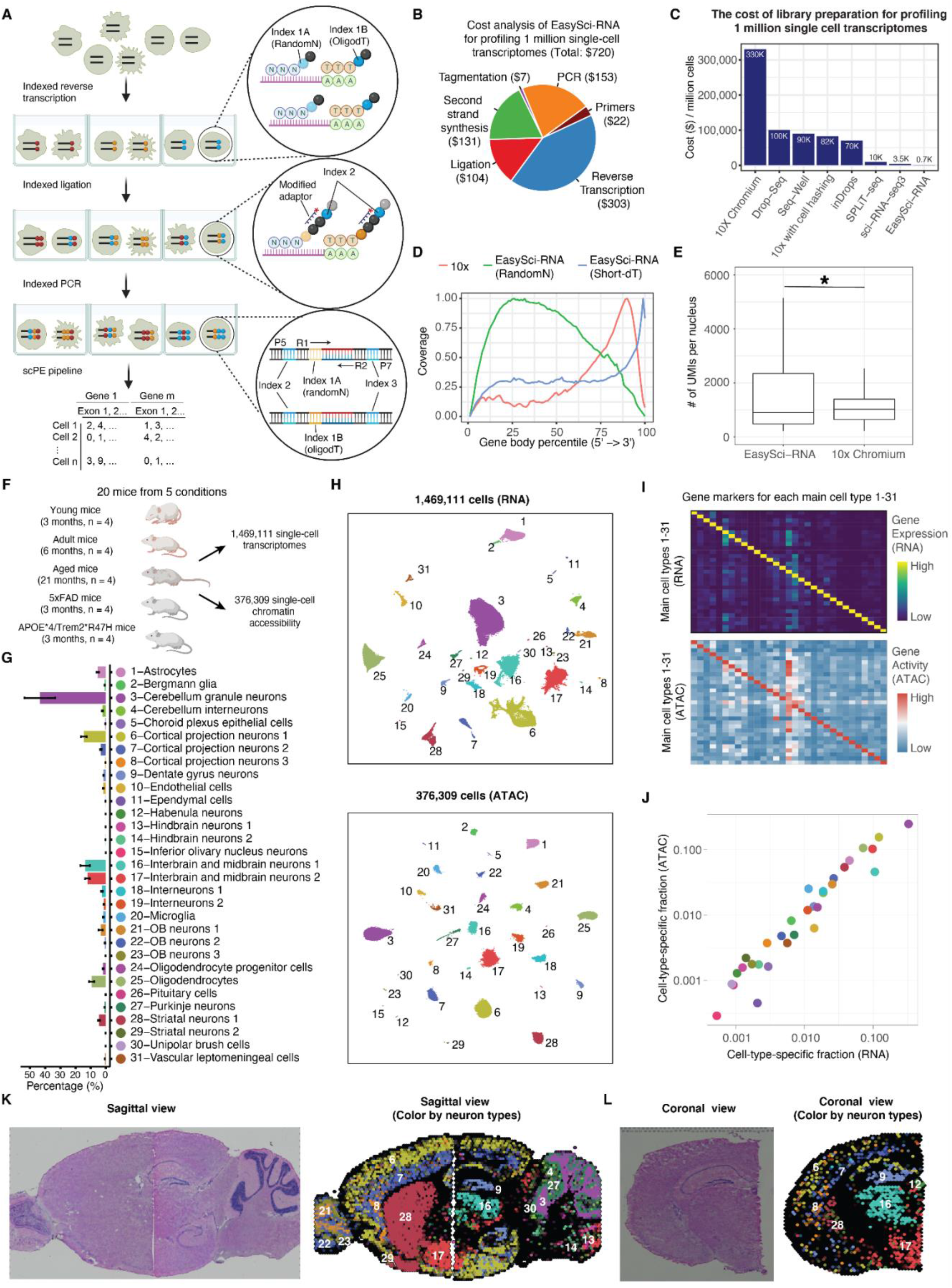
*EasySci* enables high-throughput and low-cost single-cell transcriptome and chromatin accessibility profiling across the entire mammalian brain. (A) *EasySci-RNA* workflow. Key steps are outlined in the texts. (B) Pie chart showing the estimated cost compositions of library preparation for profiling 1 million single-nucleus transcriptomes using *EasySci-RNA*. (C) Bar plot comparing different single-cell RNA-seq methods in terms of their cost of the library preparation for 1 million single-nucleus transcriptomes. The cost of the optimized sci-RNA-seq3 and SPLiT-seq were calculated using data from previous publications^13,31^. The cost of 10x with cell hashing was calculated using data from a previous report^32^. The cost of other techniques was calculated using data from a previous publication^14^. (D) Density plot showing the gene body coverage comparing single-cell transcriptome profiling using *10× genomics* and *EasySci-RNA*. Reads from indexed oligo-dT priming and random hexamers priming are plotted separately for *EasySci-RNA*. (E) Box plot showing the number of unique transcripts detected per mouse brain nucleus comparing *10× genomics* and an *EasySci-RNA* library at similar sequencing depth (∼3,800 raw reads/cell, Methods). For the box plot: middle lines, medians; upper and lower box edges, first and third quartiles, respectively; whiskers, 1.5 times the interquartile range. The star indicates p-value < 0.05 using a Wilcoxon rank-sum test. The UMIs per nucleus are calculated by combining the number of unique transcripts primed by both shortdT and random hexamer primers. (F) Experiment scheme to reconstruct a brain cell atlas of both gene expression and chromatin accessibility across different ages, sexes, and genotypes.(G) Bar plot showing the mean and standard error of the cell-type-specific proportions of the brain cell population across samples profiled by *EasySci-RNA*. (H) UMAP visualization of mouse brain cells by single-cell transcriptome (top) and chromatin accessibility (bottom), colored by main cell types in (G). (I) Heatmap showing the aggregated gene expression (top) and gene body accessibility (bottom) of the top ten marker genes (columns) in each main cell type (rows). For both RNA-seq and ATAC-seq, unique reads overlapping with the gene bodies of cell-type-specific markers were aggregated, normalized first by library size, and then scaled by the maximum expression or accessibility across all cell types. (J) Scatter plot showing the fraction of each cell type in the global brain population by single -cell transcriptome (x-axis) or chromatin accessibility analysis (y-axis). (K-L) Mouse brain sagittal (K) and coronal (L) sections showing the H&E staining (left) and the localizations of main neuron types through NNLS-based integration (right), colored by main cell types in (G). The numbers correspond to cell-type-specific cluster-ID in (G).

Leveraging the technical innovations during the development of *EasySci-RNA*, we further optimized the recently published single-cell chromatin accessibility profiling method by combinatorial indexing (sci-ATAC-seq3) ^15,16^. Critical additional improvements include (i) a tagmentation reaction with indexed Tn5 that are fully compatible with indexed ligation primers of *EasySci-RNA*; (ii) a modified nuclei extraction and cryostorage procedure to further increase the reaction efficiency and signal specificity (A comprehensive quality comparison with other scATAC protocols is shown in **Figure S3**). The detailed protocols for the *EasySci* method (RNA and ATAC), as well as the data processing pipeline, are both included as supplementary files (**Supplementary file 1-6**) to facilitate the uptake of the techniques to further enable individual laboratories to cost-efficiently generate gene expression and chromatin accessibility profiles from millions of single cells.

### A comprehensive single-cell catalog of the mouse brain in Aging and AD

We first applied the *EasySci* method to characterize cell-type-specific gene expression, and chromatin accessibility profiles across the entire mouse brain sampling at different ages, sexes, and genotypes (**Figure 1F**). We collected C57BL/6 wild-type mouse brains at three months (n=4), six months (n=4), and twenty-one months (n=4). To gain insight into the early molecular changes associated with the pathophysiology of AD, two mutants from the same C57BL/6 background at three months were included: an early-onset AD (EOAD) model (5xFAD) that overexpresses mutant human amyloid-beta precursor protein (APP) and human presenilin 1 (PS1) harboring multiple AD-associated mutations^17^; and a late-onset AD (LOAD) model (APOE*4/Trem2*R47H) that carries two of the highest risk factor mutations of LOAD, including a humanized ApoE knock-in allele and missense mutations in the mouse Trem2 gene^18,19^.

Nuclei were first extracted from the whole brain, then deposited to different wells for indexed reverse transcription (*EasySci-RNA*) or transposition (*EasySci-ATAC*), such that the first index indicated the originating sample and assay type of any given well. The resulting *EasySci* libraries were sequenced in two Illumina NovaSeq runs, yielding a total of 20 billion reads (around 10 billion for each library). After filtering out low-quality cells and potential doublets, we recovered gene expression profiles in 1,469,111 single nuclei (a median of 70,589 nuclei per brain sample, **Figure S4A; Methods**) and chromatin accessibility profiles in 376,309 single nuclei (a median of 18,112 nuclei per brain sample, **Figure S4B; Methods**) across conditions. Despite shallow sequencing depth (∼ 4340 and ∼ 16,000 raw reads per cell for RNA and ATAC, respectively), we recovered an average of 1,788 UMIs (RNA, median of 935 UMIs, an average of 828 (median 557) genes detected per nucleus, 12.8% duplication rate) and 5,515 unique fragments (ATAC, median of 3,918, 9.3% duplication rate) per nucleus (**Figure S4C-E)**, comparable to the published datasets^10,11,15^. A median of 19% of ATAC-seq reads was mapped to locations near the transcription start site (±1 kb) (**Figure S4F)**, comparable to the published sci-ATAC-seq3 approach^15^.

With UMAP visualization^20^ and Louvain clustering^21^, we identified 31 main cell types by gene expression clusters (a median of 16,370 cells per cell type; **Figure 1H; Methods**), annotated based on cell-type-specific gene markers^2^. Each cell type was observed in almost every individual, except the rare pituitary cells (0.09% of the cell population) that were missing in three out of twenty individuals (**Figure S5**). The cell-type-specific fractions in the global cell population ranged from 0.05% (Inferior olivary nucleus neurons) to 32.5% (Cerebellum granule neurons) (**Figure 1G**). An average of 74 marker genes were identified for each main cell type (defined as differentially expressed genes with at least a 2-fold difference between first and second-ranked cell types with respect to expression; FDR of 5%; and TPM > 50 in the target cell type; **Table S1**). In addition to the established marker genes, we identified many novel markers that were not previously associated with the respective cell types, such as markers for microglia (*e,g., Arhgap45* and *Wdfy4*), astrocytes (*e,g., Celrr* and *Adamts9*) and oligodendrocytes (*e,g., Sec14l5* and *Galnt5*) **(Figure S5B**).

We next sought to quantify isoform expression through a published computation pipeline^22^. We first merged random hexamer reads from each cell type in every mouse brain, yielding 613 pseudocells. We then pseudo-aligned the merged reads to the mouse transcriptome using Kallisto^23^ and produced a raw isoform count matrix. After filtering and normalization, we recovered abundance estimates for 33,361 isoforms corresponding to 12,636 genes (**Methods**). As expected, the previously identified main clusters can be readily resolved through isoform expression (**Figure S6A**). Compared with single-cell RNA-seq libraries^22^, a relatively lower fraction (∼40%) of *EasySci-RNA* reads were mapped to transcriptome with the above pipeline, potentially due to the high fraction of intronic reads in single nucleus RNA sequencing. Nevertheless, we identified certain isoforms strongly expressed in a given cell type even though their corresponding genes are not cell-type-specific (**Table S2**). For instance, *App-202*, an isoform of the amyloid precursor protein gene, exhibits differential expression in choroid plexus epithelial cells, while the expression of its parent gene (characterized by UMIs overlapping with the entirety of the *App* gene) does not display similar cell type specificity (**Figure S6B**). Similarly, *Aplp2-209*, an isoform of the amyloid beta precursor-like protein 2 gene, is differentially expressed in oligodendrocytes, while its cell-type-specificity is not detected at the gene level (**Figure S6C**). The differential expression of *Aplp2-209* in oligodendrocytes is further validated using the Tabula Muris Senis mouse aging atlas dataset^7^ (**Figure S6D**).

To reconstruct a brain cell atlas of both gene expression and chromatin accessibility, we applied a deep learning-based strategy^24^ to integrate the 376,309 single-cell chromatin accessibility profiles with gene expression data (**Figure 1H; Methods**), yielding all 31 main cell types defined by chromatin accessibility. The gene body accessibility and expression of marker genes across cell types were highly correlated (**Figure 1I**). The fraction of each cell type was highly consistent between the two molecular measurements as well (Pearson correlation r = 0.95, p-value = 6.68 × 10^−1^^6^) (**Figure 1J**). To gain more insight into the epigenetic controls of the diverse cell types in the brain, we next identified peaks of accessibility within each cell type, yielding a master set of 339,951 peaks. There was a median of 34% of reads in peaks per nucleus. UMAP dimension reduction using the resulting peak count matrix readily separates main cell types, further validating the integration-based annotations **(Figure S7A**). Through differential accessibility (DA) analysis, we identified a median of 474 differential accessible peaks per cell type (FDR of 5%, TPM > 20 in the target cell type, **Figure S7B and S7C; Table S3**). Key cell-type-specific TF regulators were discovered by correlation analysis between motif accessibility and expression patterns across diverse cell types **(Figure S7D**), such as *Spi1* in microglia^25^, *Nr4a2* in cortical projection neurons 3 ^26^, and *Pou4f1* in inferior olivary nucleus neurons ^27^.

As a step toward a spatially resolved brain atlas, we integrated our dataset with a *10× Visium* spatial transcriptomics dataset^28–30^ through a modified non-negative least squares (NNLS) approach (**Methods**). Aggregated cell-type-specific gene expression data were used as input to decompose mRNA counts at individual spatial locations of both sagittal and coronal sections of the entire mouse brain, thereby estimating the cell-type-specific abundance across locations. As expected, specific brain cell types were mapped to distinct anatomical locations (**Figure 1K and 1L**), especially for region-specific cell types such as cortical projection neurons (Clusters 6,7,8), cerebellum granule neurons (Cluster 3), and hippocampal dentate gyrus neurons (Cluster 9). The integration analysis further confirmed the annotations and spatial locations of main cell types in our single-cell datasets.

### A computational framework tailored to characterize cellular subtypes in the mammalian brain

To investigate the molecular signatures and spatial distributions of cellular subtypes in the brain, we developed a computational framework tailored to sub-cluster level analysis. Key steps include: (i) sub-clustering analysis by expression of both genes and exons to increase the clustering resolution; (ii) Integration analysis with the scATAC-seq, published single cell RNA-seq and spatial datasets to validate and map the distribution of cellular subtypes (**Figure 2**).

**Figure 2.**
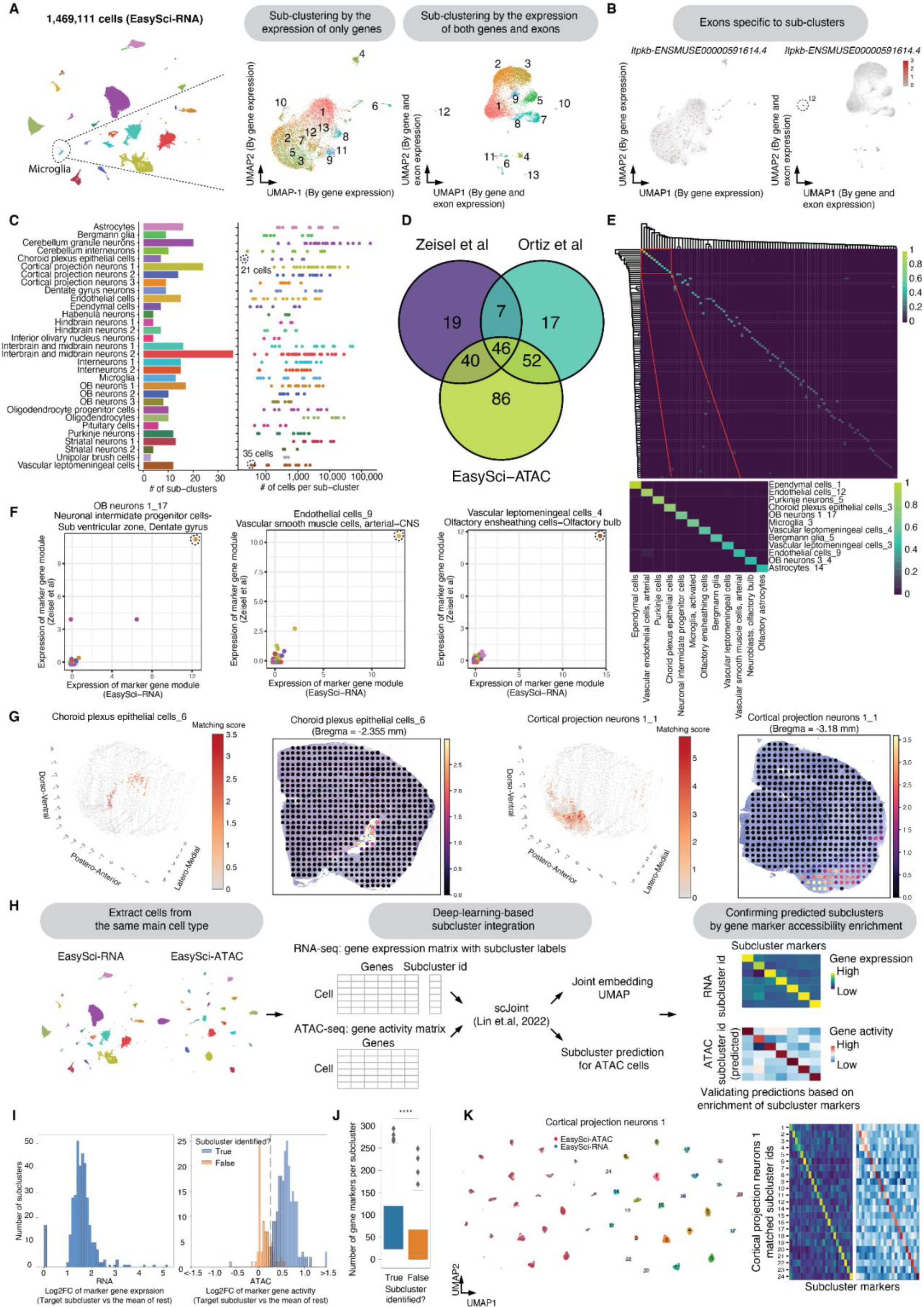
Identification of cellular subtypes in the mouse brain. (A) Schematic plot showing the computational framework for identifying cell sub-clusters. We subjected each main cell type to sub-clustering analysis based on both gene and exon expression. As an example, we performed UMAP analysis of microglia cells based on gene expression alone (left), or both gene and exon level expression (right). Cells are colored by sub-cluster ID from Louvain clustering analysis with combined gene and exon level information. Several sub-clusters cannot be separated from each other in the UMAP space by gene expression alone. (B) UMAP plots are the same as (A), showing the expression of an exonic marker *Itpkb-ENSMUSE00000591614.4* of microglia sub-cluster 12. Microglia-12 can be better separated when combining both gene and exon-level information. UMI counts for exon markers are scaled for library size, log-transformed, and then mapped to Z-scores. The values were capped at maximum 3 and minimum -3. (C) By sub-clustering analysis, we identified a total of 359 sub-clusters across 31 main cell types. The barplot (left) shows the number of sub-clusters for each main cell type. The dot plot (right) shows the number of cells from each sub-cluster. Two rare sub-clusters (choroid plexus epithelial cells-7 and vascular leptomeningeal cells-2) are circled out. (D) Venn diagram showing the number of validated subclusters using integration analysis with either Zeisel et al.^2^, Ortiz et al.^36^, or the EasySci-ATAC dataset. (E) This heatmap illustrates the similarity score, beta, between cell types derived from the EasySci dataset (rows) and the single-cell dataset from Zeisel et al.^2^ (columns). The colors represent the min-max normalized beta values obtained from cell-type correlation analysis (see Methods). (F) This display illustrates the expression of gene modules unique to subclusters across paired cell types between the EasySci and Zeisel et al. datasets^2^. Marker genes specific to EasySci subclusters were selected, then scaled to adjust for library size, log-transformed, translated into Z-scores, and finally grouped within both EasySci subclusters and corresponding cell types from Zeisel et al.^2^. (G) Using the cell2location approach^35^, we have located the spatial positions of EasySci subclusters (CPEC-6 and CPN 1-1) in a comprehensive brain spatial transcriptomics atlas^36^. The matching scores of a subcluster on the coronal sections are visualized in both 3D (left) and 2D (right). In the 3D figures, pixels with a matching score greater than 1 were retained, along with pixels at the brain’s edge to delineate the brain’s contour. (H) This illustration outlines the procedure for integrating scRNA and scATAC analyses. The single-cell gene count matrix (derived from scRNA-seq) and the single-cell gene activity matrix quantified by counting the reads overlapping with the gene body (derived from scATAC-seq) are incorporated as input into scJoint^24^ for integration analysis. Subcluster labels for each ATAC-seq cell are designated based on transfer learning, and predictions are assessed and verified by evaluating the accessibility of marker genes identified from the RNA-seq data. (I) The left histogram presents the enrichment folds (Log2FC) of marker gene expression per subcluster in the scRNA-seq dataset. The right histogram displays the enrichment folds (Log2FC) of marker gene activity per subcluster in the scATAC-seq dataset. Predicted sub-clusters in the ATAC-seq data that pass the enrichment cutoff (log2FC of gene activity > 0.25 between the target subcluster and the rest of cells) and contain more than 10 cells are considered matched. (J) Boxplots illustrate the number of markers per subcluster, comparing the subclusters that could be validated in ATAC-seq data to those that could not. (K) These figures display the subcluster integration results of cortical projection neurons 1. The UMAP plots (left) demonstrate the overlap of two molecular layers, color-coded by subcluster id; heatmaps reveal the enrichment of gene activity and gene accessibility of matching subclusters.

Rather than performing sub-clustering analysis with the gene expression alone, we exploited the unique feature of *EasySci-RNA* (*i.e.,* full gene body coverage), by incorporating both gene counts and exonic counts for principal component analysis followed by unsupervised clustering (**Methods**). The combined information significantly increased the resolution of the sub-clustering analysis. For example, we recovered several microglia subtypes that were not easily separated in clusters defined by gene expression alone (*e.g.,* sub-cluster 12 in microglia marked by an exonic marker *Itpkb-ENSMUSE00000591614.4*, **Figure 2A and 2B**). Leveraging this sub-clustering strategy, we identified a total of 359 sub-clusters, with a median of 1,038 cells in each group (**Figure 2C**). All sub-clusters were contributed by at least two individuals (median of twenty), with a median of nine exonic markers enriched in each sub-cluster (At least a 2-fold difference between first and second-ranked cell types with respect to expression; FDR of 5%; and TPM > 50 in the target sub-cluster, **Figure S9**; **Table S4**). Some subtype-specific exonic markers were not detected by conventional differential gene analysis (*e.g., Map2-ENSMUSE00000443205.3* in microglia-8, **Figure S8A)**. Notably, our strategy favors detecting extremely low-abundance cell types. For example, the smallest sub-cluster (choroid plexus epithelial cells-7) contained only 21 cells (0.001% of the brain population), representing rare pinealocytes in the brain based on gene markers such as *Tph1* and *Ddc*^33^ (**Figure S8BC**). Another example of the rare sub-clusters (vascular leptomeningeal cells-2, 35 cells) represents the tanycytes, validated by published gene markers (*e.g., Fndc3c1*, *Scn7a*^34^, **Figure S8DE**).

We next conducted rigorous validation of the identified subclusters using our scATAC-seq dataset, along with several independent datasets, which allowed us to validate the majority (75%) of the 359 cell subclusters and map their 3D spatial distributions across the mammalian brain **(Figure 2D, Methods)**. As an initial step, we incorporated cell type correlation analysis, which integrated our data with a substantial single-cell dataset featuring highly detailed cell type annotations^2^. This enabled the recognition of 112 subclusters, each of which matched with cell types documented in the previous study **(Figure 2E, Table S5)**. These corresponding cell types were further substantiated by the significant enrichment of cell-type-specific markers, exemplified by neuronal intermediate progenitor cells, vascular smooth muscle cells, and olfactory ensheathing cells **(Figure 2F)**.

As a secondary approach to validate the subclusters and infer their spatial locations in the brain, we integrated the *EasySci-RNA* dataset with the *10× Visium* spatial transcriptomics datasets^28–30^ using cell2location^35^, a Bayesian model designed to map fine-grained cell types. This integration allowed us to determine the region-specificity of the recovered cell types or sub-types. For instance, we identified cortical projection neurons subclusters mapping to different layers and regions of the cortex **(Figure S8F)** or astrocyte subtypes that specifically mapped to the olfactory bulb (sub-cluster 7), cortex (sub-cluster 3, 12), hippocampus (sub-cluster 6), corpus striatum (sub-cluster 11), midbrain (sub-cluster 4), hindbrain (sub-cluster 10) (**Figure S8G**), consistent with the known region-specificity of astrocytes^35^. Inspired by the success of the integration approach, we expanded the analysis to include an extensive spatial transcriptomics dataset encompassing 75 coronal sections of the entire mouse brain^36,35^. This permitted the spatial mapping of our identified subclusters, leading to the discovery of 122 subclusters with high spatial mapping scores (**Table S5**). The result further confirmed the notion of specific spatial distributions of subclusters. For instance, our analysis revealed that choroid plexus epithelial cells-6 were primarily situated in the lateral ventricle, whereas cortical projection neurons 1-1 were predominantly found in the amygdala (**Figure 2G**).

Additionally, we compared our detected sub-clusters with the EasySci-ATAC dataset by utilizing a deep learning-based method^24^ to integrate the snRNA-seq and snATAC-seq data from each primary cell type **(Figure 2H)**. This approach enabled us to discern 224 “corresponding subclusters” between the two molecular layers, characterized by concordant enrichment of gene activities in the target subcluster as opposed to other cells (**Figure 2I)**. As expected, the subclusters corroborated in the ATAC-seq data generally exhibit more marker genes than those which could not be validated (**Figure 2J**). As an example, the chromatin landscape for all 24 subclusters from cortical projection neurons 1 cells was recognized and corroborated by the significant enrichment of marker gene expression and activity in the target subcluster (**Figure 2K**). To investigate the cis-regulatory elements at the cell subtype resolution, we next performed a correlation-based linkage analysis (**Methods, Figure S10A**), and identified 112,665 positive links between genes and distal sites (at an empirically defined significance threshold of FDR = 0.01) (**Figure S10B-C**), unveiling a global network of putative enhancer-gene pairs shaping the molecular diversity of numerous brain cell types. Examples of previously uncharacterized enhancer-gene pairs, such as the regulatory relationship of gene *Arhgap25* and a site 39 kb away in microglia-9 (disease-associated microglia), and gene *Htr5a* and a site 32 kb away in several unipolar brush cell subtypes were presented (**Figure S10D**).

We next examined the key molecular programs underlying diverse cellular subtypes by gene module analysis. We clustered genes based on their expression variance across all 359 cell sub-clusters, revealing a total of 21 gene modules (GM) (**Figure S11-12; Table S6**). The largest gene module (GM1) corresponds to a group of housekeeping genes (*e,g.,* ribosomal synthesis) universally expressed across all sub-clusters. Several gene modules were enriched in specific cell types, such as the ependymal cell -specific gene module (GM11, enriched biological process: cilium movement, adjusted p-value = 1.2e-26) ^37^ (**Figure S11B**). Meanwhile, we detected gene modules that marked rare subtypes. For example, GM9, including genes in neuropeptide signaling (*e,g., Tbx19, Pomc*^38^), was highly enriched in a subtype of pituitary cells (PC-6) corresponding to corticotropic cells (**Figure S11B**). A similar analysis enabled us to characterize the molecular programs of other rare cell subtypes, including myeloid cells (microglia-13, 0.005% of the cell population, marked by GM19), pituitary stem cells (vascular leptomeningeal cells-12, 0.003% of the cell population, marked by GM20), as well as aforementioned pinealocytes (choroid plexus epithelial cells-7, 0.001% of the cell population, marked by GM2) (**Figure S12**). Remarkably, rare proliferating cells were identified through a cell-cycle-related gene module (GM6, enriched biological process: microtubule cytoskeleton organization involved in mitosis, adjusted p-value = 1.2e-44)^37^, including proliferating cells of neurons (OB neurons 1-17, 0.03% of the cell population), oligodendrocytes progenitor cells (oligodendrocytes progenitor cells-4, 0.04% of the cell population) and microglia (microglia-10, 82 cells, 0.006% of the cell population) (**Figure S11B**). These sub-clusters were marked by conventional proliferating markers such as *Mki67*, as well as a group of lncRNAs (*e.g., Gm29260, Gm37065*), most of which were not well-characterized in previous studies (**Figure S11C**).

### A global view of brain cell population dynamics across the adult lifespan at subtype resolution

To obtain a global view of brain cell population dynamics across the adult lifespan, we first quantified the cell-type-specific fractions recovered from cell populations in each individual mouse. We next performed differential abundance analysis across all 359 sub-clusters (**Methods**), yielding 45 significantly changed sub-clusters during the early growth stage (between 3 and 6 months) and 29 significantly changed sub-clusters upon aging (between 6 and 21 months; FDR of 0.05, at least two-fold change of cellular fractions, **Figure 3A; Table S7 and S8**). Significantly changed cell subtypes were highly consistent between male and female mice (**Figure 3B**).

**Figure 3.**
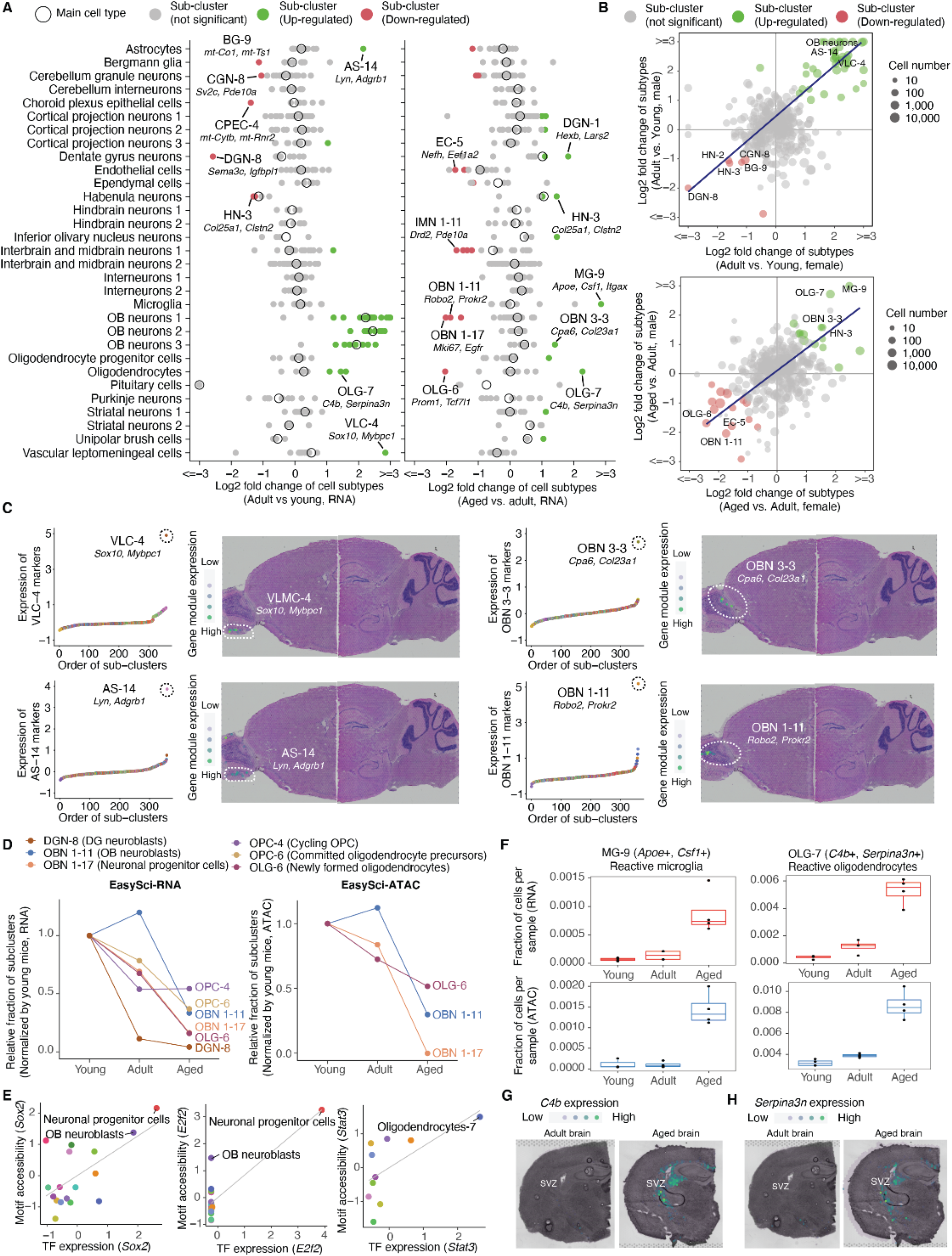
Identifying brain cell population changes across the adult lifespan at subtype resolution. (A) Dot plots showing the cell-type-specific fraction changes (*i.e.,* log-transformed fold change) of main cell types (circles) and sub-clusters (dots) in the early growth stage (adult vs. young, left plot) and the aging process (aged vs. adult, right plot) from *EasySci*-RNA data. Differential abundant sub-clusters were colored by the direction of changes. Representative sub-clusters were labeled along with top gene markers. AS, astrocytes; BG, Bergmann glia; CGN, cerebellum granule neurons; CPEC, choroid plexus epithelial cells; DGN, dentate gyrus neurons; EC, endothelial cells; HN; habenula neurons; IMN 1, interbrain and midbrain neurons 1; MG, microglia; OBN 1, OB neurons 1; OBN 3; OB neurons 3; OLG, oligodendrocytes; VLC; vascular leptomeningeal cells. (B) Scatter plots showing the correlation of the sub-cluster specific fraction changes between males and females in the early growth stage (top) and the aging stage (bottom), with a linear regression line. The most significantly changed sub-clusters are annotated on the plots. (C) Examples of development- or aging-associated subclusters are highlighted in (a) and their spatial positions. Left: scatter plots showing the aggregated expression of sub-cluster-specific marker genes across all sub-clusters. Right: plots showing the aggregated expression of sub-cluster-specific marker genes across a brain sagittal section in 10x Visium spatial transcriptomics data^28–30^. UMI counts for gene markers are scaled for library size, log-transformed, mapped to Z-scores and then aggregated. (D) Line plots showing the relative fractions of depleted subclusters across three age groups identified from *EasySci-RNA* (left) and *EasySci-ATAC* (right). (E) Scatter plots showing the correlated gene expression and motif accessibility of transcription factors enriched in OB neurons 1-17 (*Sox2* and *E2f2*, left and middle) and oligodendrocytes-7 (*Stat3*, right), together with a linear regression line. TF gene expressions are calculated by aggregating scRNA-seq gene counts for each main cluster, normalized by the library size, and then mapped to Z-scores. TF motif accessibilities are quantified by chromVar ^59^, then aggregated per main cell type and mapped to Z-scores (**Methods**). (F) Box plots showing the fractions of the reactive microglia (left) and reactive oligodendrocytes (right) across three age groups profiled by *EasySci-RNA* (top) and *EasySci-ATAC* (bottom). For all box plots: middle lines, medians; upper and lower box edges, first and third quartiles, respectively; whiskers, 1.5 times the interquartile range; and all individual data points are shown. (G-H) Mouse brain coronal sections showing the expression level of *C4b* (g) and *Serpina3n* (h) in the adult (left) and aged (right) brains from spatial transcriptomics analysis. UMI counts for gene markers are scaled for library size, log-transformed, and then mapped to Z-scores.

Almost all subtypes of olfactory bulb (OB) neurons showed a remarkable population expansion from young to adult mice (**Figure 3A, left**), consistent with the expansion of the OB region during the early growth stage^39^. Meanwhile, a rare astrocytes subtype (AS-14, *Lyn+ Adgrb1+*; 0.05% of the global population) and a vascular leptomeningeal cell subtype (VLC-4, *Sox10+ Mybpc1+*; 0.06% of the global population) showed substantial expansion (over 4-fold) in the same period. The observed cell population dynamics can be further cross-validated by two molecular layers (*i.e.,* RNA and ATAC) (**Figure S13D**). In fact, AS-14 and VLC-4 were spatially mapped to the OB region based on the expression of cell-type-specific gene markers in *10× Visium* spatial transcriptomic data^28–30^ (**Figure 3C, left**), suggesting their potential roles in OB expansion during the early growth stage. As a further illustration of this point, the astrocytes subtype AS-14 is featured with the high expression of *BAI1*, a gene marker involved in the clean-up of apoptotic neuronal debris produced during fast growth of the brain^40^. The vascular leptomeningeal cell subtype VLC-4 highly expresses gene markers of olfactory ensheathing cells (*e.g., Sox10* and *Mybpc1*^31,41^), a key cell type that supports the growth and regeneration of axons in the central nervous system^42^.

The aging-associated cell population changes between 6 and 21 months differed remarkably from the early growth stage. For example, all main cell types of OB neurons remain relatively stable during aging. Instead, we found aging-associated changes mostly in specific neuron subtypes. Key examples include the expansion of an OB neurons 3 subtype (OBN 3-3, marked by *Cpa6* and *Col23a1*) corresponding to a group of less-characterized excitatory neurons in the mitral cell layer of the OB region^43^, and the depletion of an OB neurons 1 subtype (OBN 1-11, marked by *Robo2* and *Prokr2*) corresponding to the OB neuroblasts^2,44^. These subtypes were spatially mapped to different areas of the olfactory bulb (**Figure 3C, right**), which contrasts with the early growth stage where almost all subtypes of OB neurons dramatically expanded across all regions.

We identified a total of 21 subtypes showing a marked reduction across the adult lifespan of the mouse brain. For example, the most depleted populations in the aged brain include OB neuroblasts (OBN 1 -11, marked by *Prokr2* and *Robo2*^2,44^), OB neuronal progenitor cells (OBN 1-17, marked by *Mki67* and *Egfr* ^45^), and dentate gyrus (DG) neuroblasts (DGN-8, marked by *Sema3c* and *Igfbpl1*^46^) (**Figure 3D, left**). Interestingly, the population of DG neuroblasts showed a substantial decrease even in the early growth stage, suggesting an earlier decline of DG neurogenesis compared to OB neurogenesis. In contrast to the depleted pool of neurogenesis progenitors, the proliferating oligodendrocyte progenitor cells (cycling OPCs, OPC-4, marked by *Pdgfra* and *Mki67*) remain relatively stable during aging. Instead, we detected the aging-associated depletion of the newly formed oligodendrocytes (oligodendrocytes-6 (OLG-6), marked by *Prom1* and *Tcf7l1*^45,47^) and committed oligodendrocyte precursors (OPC-6, marked by *Bmp4* and *Enpp6*^45,47,48^, indicating that the oligodendrocyte differentiation is impaired upon aging. Notably, the dynamics of multiple progenitor cell types, such as olfactory bulb neuroblasts (OBN 1-11), olfactory bulb neuronal progenitors (OBN 1-17), and newly formed oligodendrocytes (OLG-6), were effectively validated using the scATAC-seq dataset (**Figure 3D**, **Figure S14A and S14B**) and our companion study in which we tracked the dynamics of proliferating progenitor cells via metabolic labeling ^49^. Furthermore, we identified and validated certain subtype-specific transcription factor regulators using both gene expression and transcription factor motif accessibility. This includes recognized regulators of neurogenesis (for instance, *Sox2* and *E2f2*^50,51^), demonstrating the potential of our datasets to unveil key epigenetic signatures of aging-associated cell subtypes (**Figure 3E**).

In addition, we identified a total of 14 cell subtypes that exhibited a remarkable expansion in the aged brain, such as a microglia subtype (MG-9, *Apoe*+, *Csf1*+) corresponding to a previously reported disease-associated microglia^52^, and a reactive oligodendrocyte subtype (OLG-7, *C4b+*, *Serpina3n+* ^53,54^). With the chromatin accessibility dataset, we further confirmed its expansion (**Figure 3F; Figure S14B-D**), and identified its associated transcription factors. For example, the OLG-7 associated TF, *Stat3* (**Figure 3E**), plays a critical role in regulating inflammation and immunity in the brain^55^. To further validate the existence of the reactive oligodendrocyte subtype and characterize its spatial distribution in the brain, we performed a spatial transcriptomics experiment using both adult and aged mouse brains. Strikingly, we detected a significant enrichment of the reactive oligodendrocyte-specific markers (*e.g., C4b*, *Serpina3n)* around the subventricular zone (SVZ), a region critical for the continual production of new neurons in adulthood (**Figure 3G and 3H**), indicating an age-related activation of inflammation signaling around the adult neurogenesis niche.

We next explored the subtype-specific manifestation of key aging-related molecular signatures. Through differentially expressed gene analysis, we identified 7,135 aging-associated signatures across 359 sub-clusters (FDR of 5%, with at least a 2-fold change between aged and adult brains, **Table S9; Figure S15A**). Out of the 580 genes significantly altered in multiple (>= 3) subtypes, we detected 241 genes that were changed in concordant directions across subtypes (**Figure S15B**). For example, *Nr4a3*, a component of DNA repair machinery and a potential anti-aging target^56^, was significantly decreased in aged neurons, including striatal neurons, OB neurons, and interneurons. *Hdac4,* encoding a histone deacetylase and a recognized regulator of cellular senescence^57^, was significantly reduced in aged astrocytes and ependymal cells. Meanwhile, the Insulin-degrading enzyme (IDE), a key factor involved in amyloid-beta clearance^58^, showed increased expression mostly in subtypes of neurons, including interneurons, OB neurons, interbrain, and midbrain neurons. While many of these genes have been previously reported to be associated with aging, our analysis represents the first global view of their manifestations over 300 cell subtypes. In addition, we identified several non-coding RNAs that underwent age-associated changes in multiple cell subtypes, most of which showed high cell-type-specificity (e.g., *B230209E15Rik* in cortical projection neurons subtypes) but were not well-characterized previously (**Figure S15B**).

### A global view of AD pathogenesis-associated signatures and subtypes

Toward a global view of AD-associated cell population dynamics, we quantified the relative fraction of sub-clusters in the two AD models for comparison with their age-matched wild-type controls (3-month-old). We detected 16 and 14 significantly changed sub-clusters (FDR of 5%, at least two-fold change) in the EOAD (5xFAD) model and LOAD (APOE*4/Trem2*R47H) model, respectively (**Figure 4A, Table S10 and S11**). Most significantly altered subtypes showed consistent proportion changes in male and female mice (**Figure 4B**).

**Figure 4.**
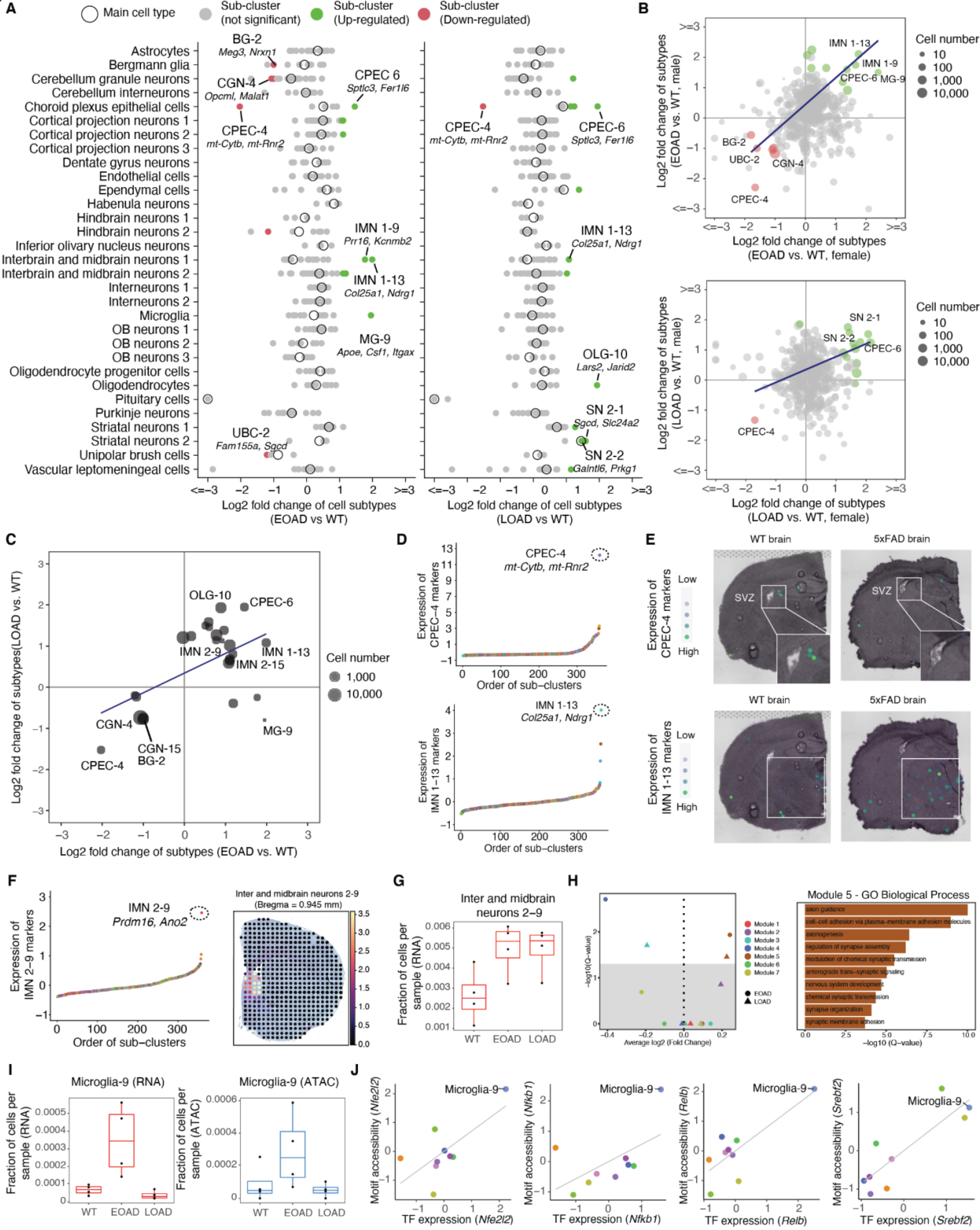
Identifying AD pathogenesis-associated cell subtypes. (A) Dot plots showing the log-transformed fold changes of main cell types (circles) and sub-clusters (dots) comparing EOAD vs. WT (left) and LOAD vs. WT (right). Differential abundant sub-clusters were colored by the direction of changes. Representative sub-clusters were labeled along with top gene markers. BG, Bergmann glia; CPEC, choroid plexus epithelial cells; IMN 1, interbrain and midbrain neurons 1; MG, microglia; OLG, oligodendrocytes; SN 2, striatal neurons 2. (B) Scatter plots showing the correlation of the log-transformed fold changes of sub-clusters (top: EOAD vs. WT, bottom: LOAD vs. WT) between males and females. (C) Scatter plot showing the correlation of the log-transformed fold changes of sub-clusters in two AD models (both compared with the wild-type). Only sub-clusters showing significant changes in at least one AD model are included. (D) Scatter plots showing the aggregated expression of gene markers of two cell subtypes (top: choroid plexus epithelial cells-4; bottom: the interbrain and midbrain neurons 1-13) across all sub-clusters from *EasySci-RNA* data. UMI counts for gene markers are scaled for library size, log-transformed, mapped to Z-scores, and then aggregated. (E) Brain coronal sections showing the spatial expression of subtype-specific gene markers of two subtypes (top: choroid plexus epithelial cells-4; bottom: the interbrain and midbrain neurons 1-13) in the WT and EOAD (5xFAD) brains in 10x Visium spatial transcriptomics data. (F) Scatter plot showing the aggregated expression of gene markers of Interbrain and midbrain neurons 2-9 across all sub-clusters from *EasySci-RNA* data (left) and the mapping score per pixels of the IMN 2-9 subcluster cells on the Bregma 0.945 mm coronal section highlighting the lateral septal nucleus region^36^ (right). UMI counts for gene markers are scaled for library size, log-transformed, mapped to Z-scores, and then aggregated. (G) Box plots showing the fraction of Interbrain and midbrain neurons 2-9 cells across different conditions profiled by *EasySci-RNA*. For all box plots in this figure: middle lines, medians; upper and lower box edges, first and third quartiles, respectively; whiskers, 1.5 times the interquartile range; and all individual data points are shown. (H) Gene module expression differences in both AD models (left) and the top enriched GO biological processes pathways of module 5 genes (right). The gray box indicated the modules without significant differences. (I) Box plots showing the fraction of microglia-9 cells across different conditions profiled by *EasySci-RNA* (left) or *EasySci-ATAC* (right). For all box plots in this figure: middle lines, medians; upper and lower box edges, first and third quartiles, respectively; whiskers, 1.5 times the interquartile range; and all individu al data points are shown. (J) Scatter plot showing the correlated gene expression and motif accessibility of four transcription factors (*Nfe2l2*, *Nfkb1*, *Relb,* and *Srebf2*) enriched in microglia-9, together with a linear regression line. TF gene expressions are calculated by aggregating scRNA-seq gene counts for each main cluster, normalized by the library size, and then mapped to Z-scores. TF motif accessibilities are quantified by chromVar ^59^, then aggregated per main cell type and mapped to Z-scores (**Methods**).

While these two AD mutants involved different genetic perturbations, the significantly altered cell subtypes were highly concordant (**Figure 4C**). For example, a rare choroid plexus epithelial cell subtype (CPEC-4, 0.02% of the total brain cell population) was strongly depleted (> two-fold decrease) in both AD models. This cell type is marked by significant enrichment of multiple mitochondrial genes, including *mt-Rnr1*, *mt-Rnr2*, *mt-Co1*, *mt-Cytb*, *mt-Nd1*, *mt-Nd2*, *mt-Nd5,* and *mt-Nd6*. Out of these gene markers, *mt-Rnr2* is involved in synthesizing neuroprotective factors against neurodegeneration by suppressing apoptotic cell death^60^. Other markers (*e.g., mt-Rnr1* and *mt-Nd5*) are associated with the phosphorylated Tau protein levels in cerebrospinal fluid^61^. While this subtype was rarely detected in our single-cell ATAC data, we could map the cell subtype to the area around the subventricular zone by the expression of its cell-type-specific markers in the spatial transcriptomics data (**Figure 4D and 4E, top**). Furthermore, our spatial transcriptomics experiment validated the depletion of this cell type in the EOAD (5xFAD) model (**Figure 4E**), suggesting a potential role of choroid plexus epithelial cell-specific mitochondrial dysfunction in neurodegenerative diseases.

By contrast, another choroid plexus epithelial cell subtype (CPEC-6, 0.045% of the total brain cell population; marked by *Sptlc3*+, *Fer1l6*+) expanded in both AD models (over two-fold increase) (**Figure 4B**). It is marked by the gene *Sptlc3,* which encodes a complex subunit that catalyzes the synthesis of sphingolipids, a group of bioactive molecules contributing to amyloid-beta production and Alzheimer’s pathogenesis^62^. Furthermore, we identified another rare interbrain and midbrain neuron subtype (IMN 1-13, 0.61% of the total brain population, marked by *Col25a1+, Ndrg1+*) that expanded considerably in both AD models (**Figure 4C**). This subtype is characterized by the expression of *Col25a1*, a membrane-associated collagen reported to promote intracellular amyloid plaque formation in mouse models^63^. Indeed, our spatial transcriptomic experiment detected an up-regulation of IMN 1-13 specific gene markers in the thalamus region of the 5xFAD mouse brain (**Figure 4D and 4E, bottom**), further validating the AD-associated neuron subtype change detected from single-cell transcriptome study. Additionally, we detected the AD-associated augmentation of a septal nuclei neuron subtype (**Figure 4F-H**) exhibiting heightened expression of an identified gene module enriched in genes related to axonogenesis (*e.g., Nrp1, Slit2*) and synaptogenesis (*e.g., Ptprd, Nrxn1*), and consistent in both the EOAD and LOAD models. This finding aligns with the observed enlargement of the septal nuclei region in human patients several years before the onset of memory decline^64^, although it has not been extensively characterized in murine brains. This underscores the utility of scalable single-cell approaches for identifying disease-associated rare cell types from under-explored regions.

Differences between the two AD models were identified. For instance, we observed a substantial expansion of the microglia subtype 9 (MG-9, 0.026% of the total brain cell population, marked by *ApoE+, Csf1+*) in the early-onset 5xFAD mice, aligning with previous reports^52^. This reactive microglia subtype was similarly expanded in the aged group but was not apparent in the late-onset APOE*4/Trem2*R47H model at three months of age (**Figure 4I, left**), potentially indicating a correlation with the onset of the disease. Consistent proportion changes were noted within the chromatin accessibility dataset (**Figure 4I, right**). To further elucidate the molecular mechanisms underpinning this reactive microglia subtype, we integrated both transcriptome and chromatin accessibility profiles and identified 199 genes differentially expressed in the reactive microglia subtype, many of which (44%) can be validated by the promoter accessibility (**Figure S14D**). The analyses also revealed key transcription factors exhibiting a consistent pattern of cell-type-specific gene expression and motif accessibility (**Figure 4J**), including transcription factors linked to the NF-kappa B signaling pathway (e.g., *Nfkb1* and *Relb*^65^) and those involved in oxidative stress protection (*e.g., Nfe2l2*^66^), and cholesterol homeostasis (*e.g., Srebf2*^67^). These molecular pathways may have critical regulatory roles in microglia specification and expansion during aging and AD. Furthermore, we performed linkage disequilibrium score regression analysis ^68^ to quantify the enrichment of genetic variants linked to human traits within cell-type-specific DEpeaks. Using DEpeaks across main cell types and across different microglia subtypes, we observed significant enrichment of AD heritability in microglia cells both at the main cell type level and particularly in the microglia-9 subcluster (**Figure S14E and S14F**), highlighting the role of DAM in AD pathogenesis.

Subsequently, we explored subtype-specific manifestations of key AD-related molecular signatures. Through differentially expressed gene analysis (**Methods**), we identified 6,792 and 7,192 sub-cluster-specific DE genes in the 5xFAD (EOAD) model and the APOE*4/Trem2*R47H (LOAD) model, respectively (**Figure S15C and S15E, Table S12 and S13**). Notably, we observed a global down-regulation of the mouse *Apoe* gene across a wide array of subclusters in the APOE*4/Trem2*R47H mice (**Figure S15F)**. This phenomenon could be attributable to the potential replacement of the *Apoe* gene with the human sequence, which does not align with the mouse genome. Concurrently, we detected a global alteration of *Thy1* across numerous neuron types in the 5xFAD mice, which aligns with the overexpression of all transgenes introduced in the 5xFAD model under the *Thy1* promoter (**Figure S15D)**.

Many AD-associated gene signatures exhibited concordant changes across cellular subtypes (**Figure S15D and S15F**). For example, markers involved in unfolded protein stress (*e.g., Hsp90aa1*) and oxidative stress (*e.g., Txnrd1*) were significantly upregulated in an overlapping set of neuron subtypes in the early-onset 5xFAD mice (**Figure S15D**), indicating increased stress levels and cellular damages in neurons even in the early stage of the EOAD brain. Meanwhile, the expression of *Reln*, a gene encoding a large secreted extracellular matrix protease involved in the ApoE biochemical pathway^69^, significantly decreased in various cell types (*e.g.,* OB neurons, interbrain and midbrain neurons, vascular cells, oligodendrocytes) in both early- and late-onset models (**Figure S15D and S15F**). This aligns with earlier studies that reported detectable depletion of *Reln* even before the onset of amyloid-beta pathology in the human frontal cortex^70^. Other intriguing observations included the overall upregulation of *Ide*, a gene implicated in amyloid-beta degradation, in the late-onset model, a pattern similar to the aged brain (**Figure S15B and S15F**). We also identified less-characterized genes. For example, *Tlcd4*, a gene involved in lipid trafficking and metabolism^71^, was significantly downregulated in thirty-five subclusters across a wide range of cell types (*e.g.,* OB neurons, vascular cells, oligodendrocytes) in the EOAD mice (**Figure S15D**), indicating a potential interplay between the lipid homeostasis and cellular changes in the early stage of AD.

Despite the distinct genetic perturbations or disease onsets in the two AD mouse models, the alterations in cell-type-specific molecular profiles were remarkably consistent. Illustrative of this, we detected 559 sub-cluster-specific DE genes shared between two AD mutants, such as genes involved in epilepsy (Adjusted p-value = 0.02, *e.g., Gria1, Med1, Plp1*)^37^ and oxidative stress protection pathway (Adjusted p-value = 0.05, *e.g., Arnt, Nfe2l2*)^37^. Intriguingly, an overwhelming majority (99%, or 555 of the 559) of these shared DE genes exhibited concurrent alterations in both AD mutants (Pearson correlation coefficient r = 0.96, p-value < 2.2e-16, **Figure S15G**). These findings suggest the presence of common molecular mechanisms between early- and late-onset AD models.

To systematically investigate the connection between aging and AD associated changes, we employed a ridge regression model to build transcriptomic aging clocks for each primary cell type using 80% of the wild-type mouse cells at 3, 6, and 21 months old, following the procedure detailed in^72^. We first merged cells with similar transcriptomes to minimize single-cell dropout effects, and then applied this model to predict the natural logarithm of the biological age of cells from both AD mouse models as well as the holdout wild-type data. Our findings revealed significantly accelerated biological age in both AD models (Mann-Whitney U test, p-value <= 1*10^-142^ for both EOAD and LOAD, **Figure S15H**). While most cell types demonstrated accelerated aging-related molecular changes, specific cell types only exhibited these signs in the late-onset model (**Figure S15I**). This is further validated by the changes of aging-associated gene signatures, such as the up-regulation of *Neat1* expression in astrocytes and the increased expression of *Zfp423* in Dentate gyrus neurons (**Figure S15K**), consistent across both aged and AD models.

### Identification of dysregulated gene signatures in human AD brains

To further elucidate molecular signatures associated with AD pathogenesis, we sequenced a total of 118,240 single-nuclei transcriptomes (a median of 5,585 nuclei per sample, with the sequencing depth of 13,850 raw reads and a median of 1,109 UMIs per nucleus, **Figure S16A and S16B**) from twenty-four human brain samples across two brain regions (hippocampus, superior and middle temporal lobe (SMTG)), derived from six Alzheimer’s disease patients and six age- and sex-matched controls (**Table S14**). Thirteen main cell types were identified through integration analysis with the mouse dataset and validated by the cluster-specific expression of known markers (**Figure 5A** and **Figure S16C-E**).

**Figure 5.**
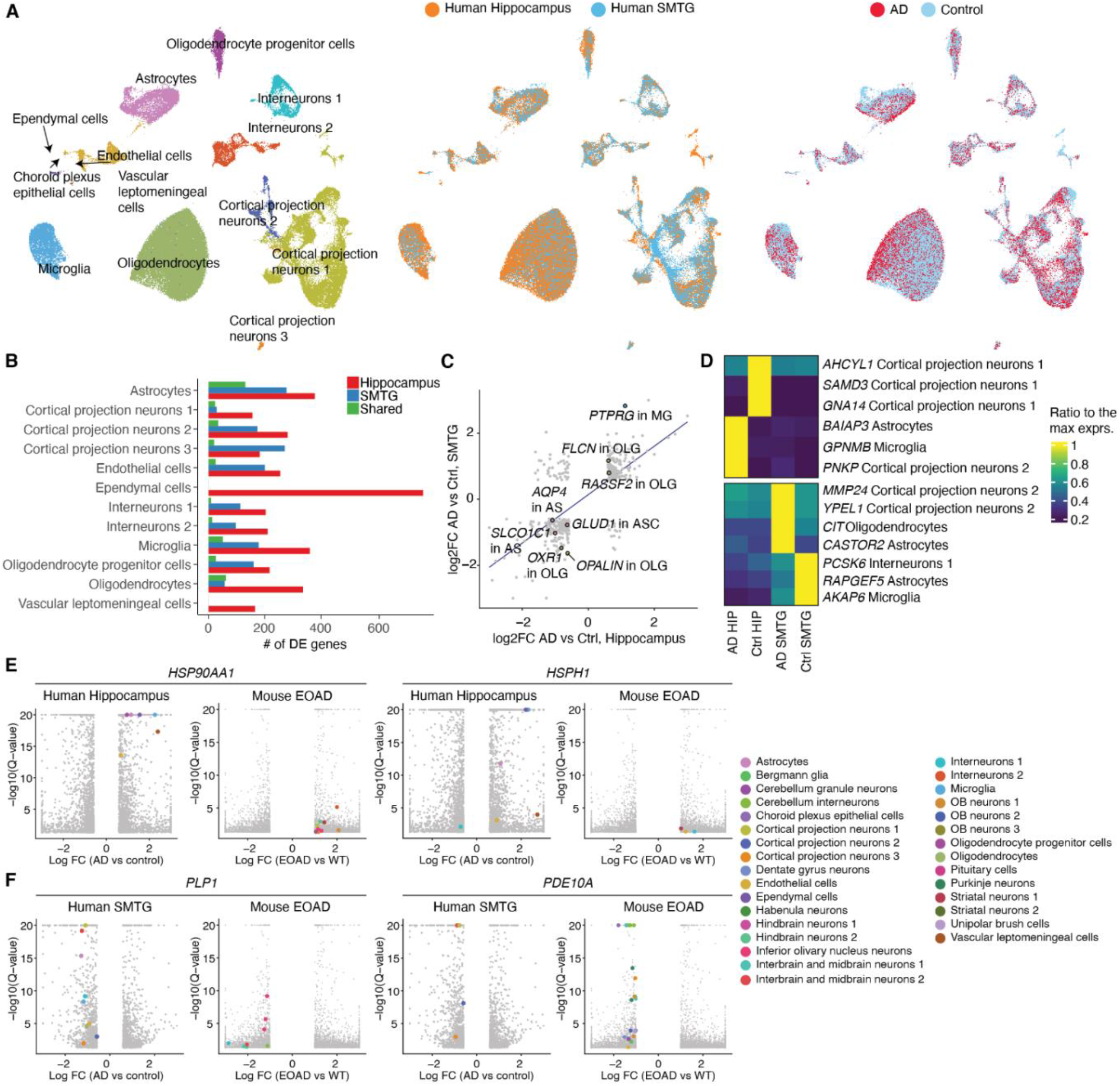
Identifying AD pathogenesis-associated gene expression signatures across regions and cell types in human brains. (A) UMAP visualization of single-cell transcriptomes of all human brain cells, colored by main cell types (left), region (middle) and conditions (right). (B) Bar plot showing the number of differentially expressed genes between AD and control samples in each cell type. DE genes are colored by whether they are unique to each region or shared between two regions. Of note, choroid plexus epithelial cells and vascular leptomeningeal cells were not included into the differential gene expression analysis in SMTG due to their low cell numbers. (C) We detected 394 DE genes significantly changed within the same main cell type in both regions. The scatterplot shows the correlation of the log2-transformed fold changes of these 394 shared DE genes in Hippocampus (x-axis) and in SMTG (y-axis). Key genes are annotated and colored by their corresponding main cell types. AS, astrocytes; MG, microglia; OLG, oligodendrocytes. (D) Heatmaps showing examples of region-specific DE genes for the hippocampus (left) and SMTG (bottom). Gene expressions were quantified as transcripts per million in the corresponding cell types in each group, and normalized to the maximum expression across groups. (E-F) Volcano plots showing the examples of top differentially expressed (DE) genes between the AD and control samples across main cell types in human brains or between EOAD and WT samples across cell subclusters in mouse brains. Highlighted genes are colored by the main cell type identity.

We next sought to discern the region- and cell-type-specific molecular signatures associated with human AD pathogenesis. By differential gene expression analysis, we identified a total of 4,171 and 2,149 cell-type-specific DE genes in the hippocampus and SMTG, respectively (**Figure 5B, Table S15**). 349 genes were significantly changed in the same cell type from two distinct regions, among which 332 were altered in concordant directions (**Figure 5C**, Pearson correlation coefficient r = 0.68, p-value < 2.2e-16). For example, oligodendrocytes in AD samples from both regions exhibited decreased expression of the oligodendrocyte terminal differentiation factor *OPALIN*^73^ and the oxidation stress protector *OXR1*^74^. Concurrently, we observed an upregulation of genes related to programmed cell death (*e.g., FLCN* and *RASSF2*)^75,76^, suggesting an elevated stress level in oligodendrocytes from AD brains. Other examples include the microglia-specific upregulation of *PTPRG*, a receptor protein tyrosine phosphatase that plays a key role in mediating AD-associated neuronal death^77^, and the downregulation of several transmembrane transporters (*e.g., AQP4* and *SLCO1C1*) and neurotransmitter metabolism enzymes (*e.g., GLUD1*) in astrocytes, indicating potential impairment of the blood-brain barrier^78^ and an altered metabolic state^79^.

Interestingly, some AD-associated gene signatures exhibited region-specific expression patterns. For example, *GPNMB*, a transmembrane glycoprotein associated with microglia activation in AD brains^80^, showed increased expression in the microglia from the hippocampus but not from the SMTG (**Figure 5D, top**). On the other hand, *MMP24*, a member of the metalloproteinase family implicated in AD pathogenesis^81^, showed increased expression in cortical projection neurons unique within the SMTG (**Figure 5D, bottom**). Notably, inhibition of MMP24 has been demonstrated to decrease amyloid-beta levels and promote cognitive functions in mouse models^82^, suggesting its potential role as a novel therapeutic target for AD.

Finally, we explored the human-mice relevance for AD-associated gene signatures and molecular pathways. Despite differences in the species and disease stages between the two datasets, several genes encoding heat shock proteins (*e.g., HSP90AA1*, *HSPH1*) were upregulated across multiple cell types in both EOAD mouse brains and the human hippocampus (**Figure 5E**). The elevated expression of the chaperon system potentially serves as a compensatory mechanism to reduce the formation of toxic oligomeric assemblies in AD brains^83^, further validating the dysfunction of proteostasis as a molecular marker of AD^84^. Meanwhile, we identified down-regulated genes in both human and mice (**Figure 5F**). One of the examples, *PLP1*, was reported as a subtype-specific driver gene contributing to AD pathogenesis^85^. Another gene, *PDE10A*, plays a key role in promoting neuronal survival, with its reduction detected in our datasets and multiple neurodegenerative diseases (*e.g.,* Huntington’s disease^86^, Parkinson’s disease^87^). Importantly, the above-mentioned trends were readily validated by another recently published single-cell dataset investigating Alzheimer’s disease in the human prefrontal cortex^6^ (**Figure S17**). In summary, the human-mice relevance analysis identified species-conserved genetic programs associated with Alzheimer’s pathogenesis.

## Discussion

In this study, we introduced EasySci, a cost-effectively technical framework for individual laboratories to generate gene expression and chromatin accessibility profiles from millions of single cells. With the technique, we generated a global view of aging and Alzheimer’s pathogenesis-associated cell population dynamics associated with aging and AD pathogenesis, leveraging the profiling of approximately 1.5 million single-cell transcriptomes with full gene body coverage and about 380,000 single-cell chromatin accessibility profiles across the entire mammalian brains spanning various age and genotype groups. The result datasets enable the identification of over 300 cellular subtypes throughout the brain, including highly rare cell types (*e.g.,* pinealocytes, tanycytes) representing less than 0.01% of the total brain cell population. Furthermore, we discovered region-specific effects attributable to aging and AD through high-resolution spatial transcriptomic analysis, and examined the manifestation of molecular signatures associated with aging and AD on a cell-type-specific basis.

As highlighted by our granular sub-cluster level analysis, the effects of aging and AD on the global brain cell population are profoundly cell-type-specific. While most brain cell types stay relatively stable under various conditions, we identified over fifty cell subtypes exhibiting significant alterations (exceeding two-fold change) in brains affected by aging and AD models. Many of these cell subtypes were rare and potentially overlooked in conventional “shallow” single-cell analysis. For example, the aging brain is characterized by the depletion of both rare neuronal progenitor cells and differentiating oligodendrocytes, associated with the enrichment of a *C4b+ Serpina3n+* reactive oligodendrocyte subtype surrounding the subventricular zone (SVZ), suggesting a potential interplay between oligodendrocytes, localized inflammatory signals, and the stem cell niche.

The lack of reliable mouse models remains one of the biggest challenges in studying late-onset Alzheimer’s disease. The novel APOE*4/Trem2*R47H model aims to overcome this limitation by introducing two of the strongest late-onset Alzheimer’s disease-associated mutations^88^. However, limited validation is available to assess whether this novel model shows any characteristics of Alzheimer’s disease. Moreover, the Trem2 mutation introduced in the LOAD model may lead to other anomalies, such as the expression of a novel splice variant of Trem2. Despite these potential limitations, we found consistent molecular and cellular population dynamics between the well-established 5xFAD and the novel APOE*4/Trem2*R47H model, underscoring that the novel LOAD model indeed manifests indicators of Alzheimer’s disease. For example, we observed shared subtypes that were depleted (*e.g., mt-Cytb+ mt-Rnr2+* choroid plexus epithelial cell) or enriched (*e.g., Col25a1+ Ndrg1+* interbrain and midbrain neuron) in both early- and late-onset AD mutant brains, validated by single-cell RNA-seq from both sexes as well as spatial transcriptomics analysis. On the other hand, differences were also observed between the two AD models, as expected by the different onset times. Most notably, the missing of the disease-associated microglia population increase in the LOAD model could be explained by the lack of amyloid deposition in the mouse model^89^ or by genetic perturbations, as both Trem2 and Apoe play a role in the activation of this cell population^52^.

In addition, we investigated AD-associated gene signatures in human brains by profiling over 100,000 single-nucleus transcriptomes derived from twenty-four human brain samples from control and AD patients, across two distinct anatomical locations. While most AD-associated gene dynamics are profoundly cell-type- and region-specific, we identified dysregulated genetic signatures that are conserved between different anatomical locations in the human brains. Moreover, integrating the human and mouse brain datasets further revealed molecular pathways shared between human AD patients and mouse AD models, which suggests that the mouse AD model can serve as a model system to investigate the function and regulation of these conserved features associated with AD or neuronal dysfunction.

Of note, there are several inherent limitations of the study. First, the analysis covers only around 2% of the total mouse brain population (estimated at approximately 100 million cells), which means extremely rare cell subtypes may still be overlooked. Additionally, the relatively shallow sequencing depth applied in our research might hinder the detection of lowly expressed transcripts or minor aging-related cellular state changes. Nevertheless, the validity of our key biological findings is reinforced by the consistent results across different sexes (male vs. female), genotypes (EOAD vs. LOAD), and orthogonal approaches (such as comparisons between single-cell transcriptome, chromatin accessibility, or spatial transcriptomics). This lends significant credence to our discoveries, even when considering the limitations of the study.

In summary, we have showcased the power of highly scalable single-cell genomics to delve into the dynamics of rare cell types, unearthing novel subtypes associated with development, aging, and disease. Though our focus was centered on brain tissues, the strategic approach we’ve employed could be readily extended to systematically explore cellular states across an entire organism. Such exploration could illuminate the cell populations most susceptible to aging and other diseases, opening up pathways to develop targeted therapeutic strategies.

## Acknowledgments

We thank all members of the Cao lab for helpful discussions and feedback. We thank Dr. Jay Shendure (University of Washington) for insightful feedback on this work. We also thank members of the Rockefeller University Genomics Resource Center (SCR_020986), High-Performance Computing Resource center, and Comparative Bioscience Center for their exceptional assistance with library sequencing and animal maintenance.

## Funding

This work was funded by grants from NIH (DP2HG012522, R01AG076932, and RM1HG011014 to J.C.; P30AG072946 and P01AG078116 to P.T.N.; R01AG072758 to L.G.) and the Sagol Network GerOmic Award for J.C.

## Author contributions

J.C. and W.Z. conceptualized and supervised the project. J.L. and A.S. developed the experimental and computation pipeline for *EasySci-RNA* profiling of all samples. G.B. and Z.L. developed the experimental and computation pipeline for *EasySci-ATAC* profiling of all samples. A.A. performed the 10x Visium spatial transcriptomics experiment. S.A. and P.N. processed the human brain samples for single-cell profiling experiments. A.S. and Z.L. performed the downstream analysis with assistance from E.M., A.L., A.E., Z.X., and Z.Z. J.B. and A.S. developed the website. J.C., W.Z., Z.L., and A.S. wrote the manuscript with input and biological insight from P.N., L.G. and other co-authors.

## Competing interests statement

J.C., W.Z., A.S. and J.L. are inventors on pending patent applications related to EasySci-RNA-seq. Other authors declare no competing interests.

## Supplementary Figures

**Figure S1.**
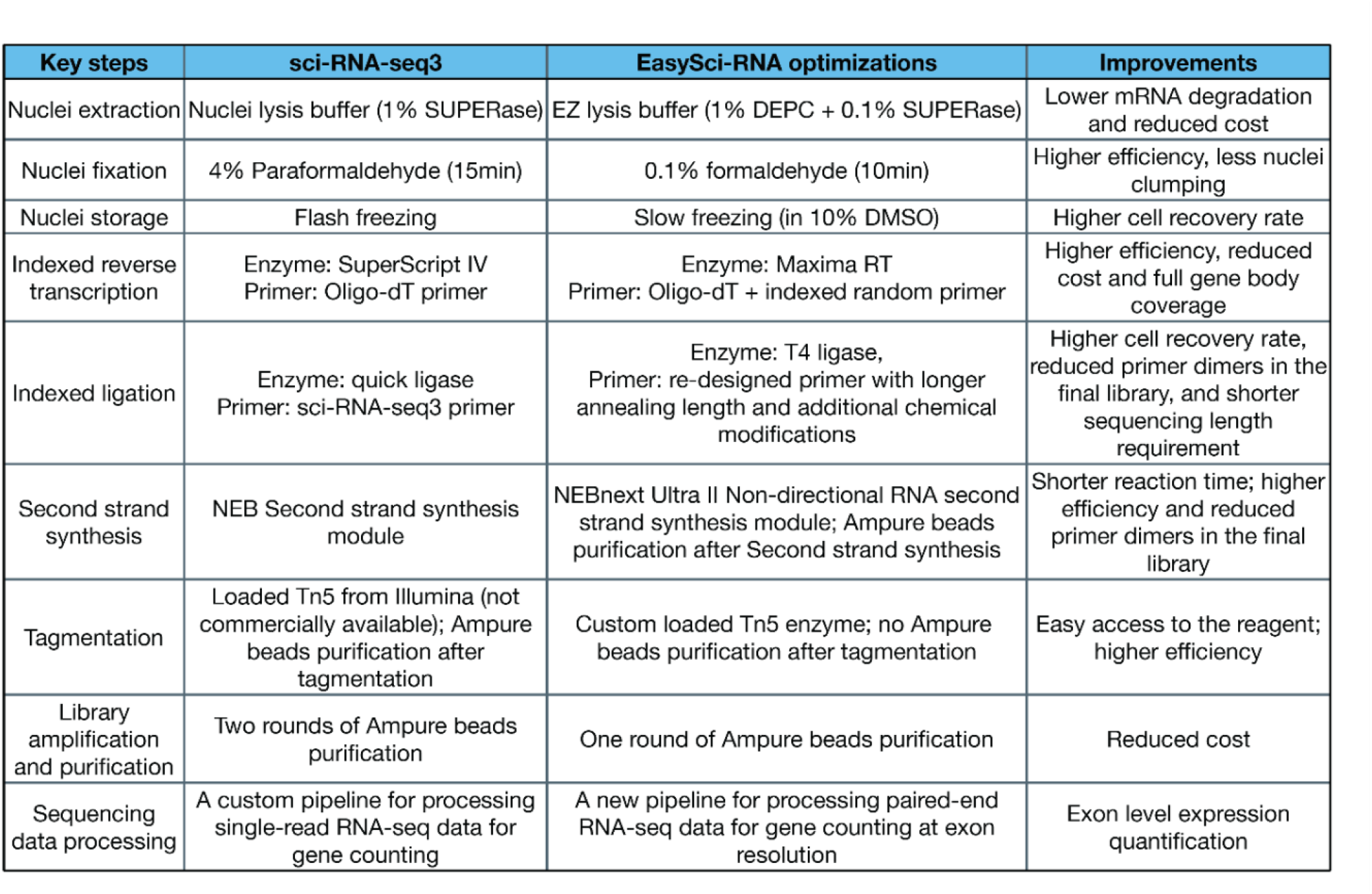
Summary of key optimizations of *EasySci-RNA* compared to published single-cell RNA-seq by combinatorial indexing (sci-RNA-seq3^11^).

**Figure S2.**
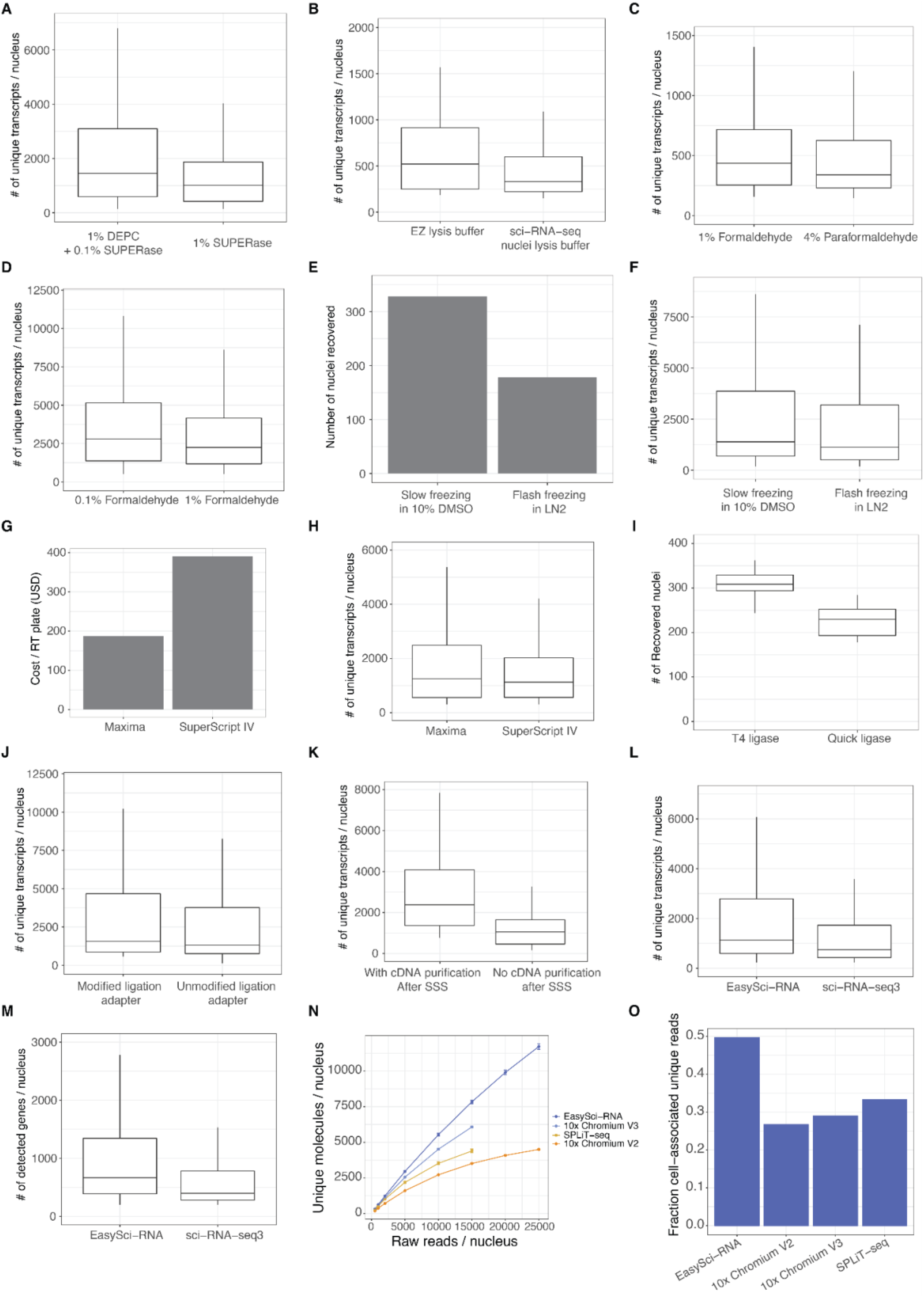
Representative examples showing the performance of optimized conditions of *EasySci-RNA*. (A-B) Box plots showing the number of unique transcripts detected per nucleus in different lysis conditions: 1% DEPC vs. no DEPC in lysis buffer (A); EZ lysis buffer vs. nuclei lysis buffer used in the published sci - RNA-seq3 ^11^ (B). For all box plots in this figure: middle lines, medians; upper and lower box edges, first and third quartiles, respectively; whiskers, 1.5 times the interquartile range. (C-D) Box plot showing the number of unique transcripts detected per nucleus across different fixation conditions: 1% formaldehyde vs 4% paraformaldehyde (C); 0.1% formaldehyde vs. 1% formaldehyde (D). (E-F) We compared two conditions for preserving the fixed nuclei. The slow freezing condition (in 10% DMSO) outperformed the flash freezing condition in sci-RNA-seq3 ^11^ by increasing the number of nuclei recovered in the experiment (E) and the number of unique transcripts detected per nucleus (F). (G-H) Maxima reverse transcriptase significantly reduces the enzyme cost (G) without affecting the number of transcripts detected per nucleus (H). (I) *EasySci-RNA* used T4 ligase instead of quick ligase for a higher recovery rate of nuclei. (J) We used chemically modified ligation primers in *EasySci* (Methods), which greatly reduced primer dimers in the following PCR reaction and slightly increased the number of unique transcripts detected per nucleus. (K) Additional cDNA purification step after second strand synthesis increased the number of unique transcripts per nucleus. (L-M) Comparison of the number of unique transcripts and the number of detected genes in EasySci-RNA and sci-RNA-seq3 with the same sequencing depth (∼2,500 reads/cell) (N) Line plots showing the median number (with standard error) of unique transcripts detected per cell from a deep sequenced EasySci-RNA, a published 10x Genomics Version 2 library ^14^, a 10x Genomics Version 3 library ^90^ and a SPLiT-seq library ^31^, both profiling mouse brains. To measure the reaction efficiency of the two methods in high-quality cells, we first selected the top 1000 highest-quality cells from both datasets. Then we subsampled these cells to different sequencing depths and calculated the number of unique transcripts detected per cell. (O) Comparison of the fraction of cell-associated unique transcripts from the total number of raw reads across EasySci-RNA, *10× genomics Version 2* ^14^, *10× genomics Version 3* ^90^ *and SPLiT-seq* ^31^. For this comparison, the first PCR batch of the large-scale EasySci-RNA dataset was compared to a publicly available 1*0X genomics* datasets and a SPLiT-seq dataset at the same sequencing depth (∼3,800 raw reads/cell).

**Figure S3.**
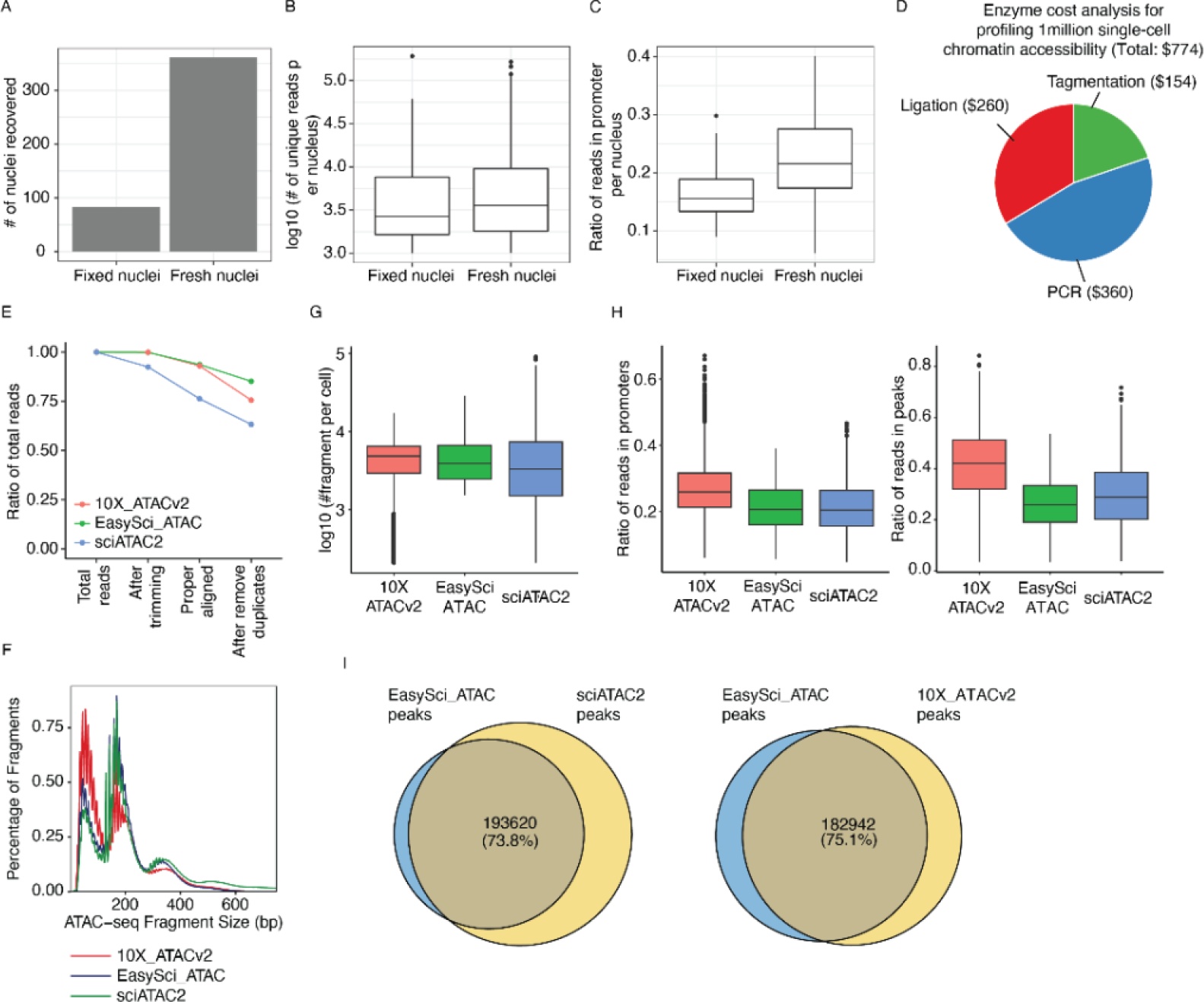
Representative examples showing the performance of optimized conditions of *EasySci-ATAC* and quality comparison with other single-cell ATAC protocols. (A-C) We compared two fixation conditions: nuclei were either fixed with 1% formaldehyde for 10 minutes at room temperature or directly used for tagmentation without fixation. The unfixed condition outperformed the fixed condition by increasing cell recovery (A), the number of reads (B), and the ratio of reads in promoters (C) per nucleus. For all box plots in this figure: middle lines, medians; upper and lower box edges, first and third quartiles, respectively; whiskers, 1.5 times the interquartile range; circles, outliers. (D) Pie chart showing the estimated enzyme cost compositions of library preparation for profiling 1 million single-cell chromatin accessibility profiles using *EasySci-ATAC*. (E) Lineplot showing the ratio of reads loss during each data processing step comparing EasySci -ATAC, 10×-ATACv2 and sci-ATAC-seq (refered as ‘sciATAC2’ in the figure). (F) Histogram showing the fragment length distributions across EasySci-ATAC, 10×-ATACv2 and sci-ATAC-seq. (G) Boxplot showing the number of unique fragments comparing EasySci-ATAC, 10×-ATACv2 and sci-ATAC-seq. (H) Boxplots showing and the number of reads mapped to promoters (left, defined as ±1kb around TSS) and peaks (right) comparing EasySci-ATAC, 10×-ATACv2 and sci-ATAC-seq. Peak calling was performed on each dataset separately and peaks were merged to a union peak set. (I) Pieplot showing the number of peaks that can be repeatedly identified between EasySci-ATAC and sci-ATAC-seq (left) and between EasySci-ATAC and 10×-ATACv2 (right).

**Figure S4.**
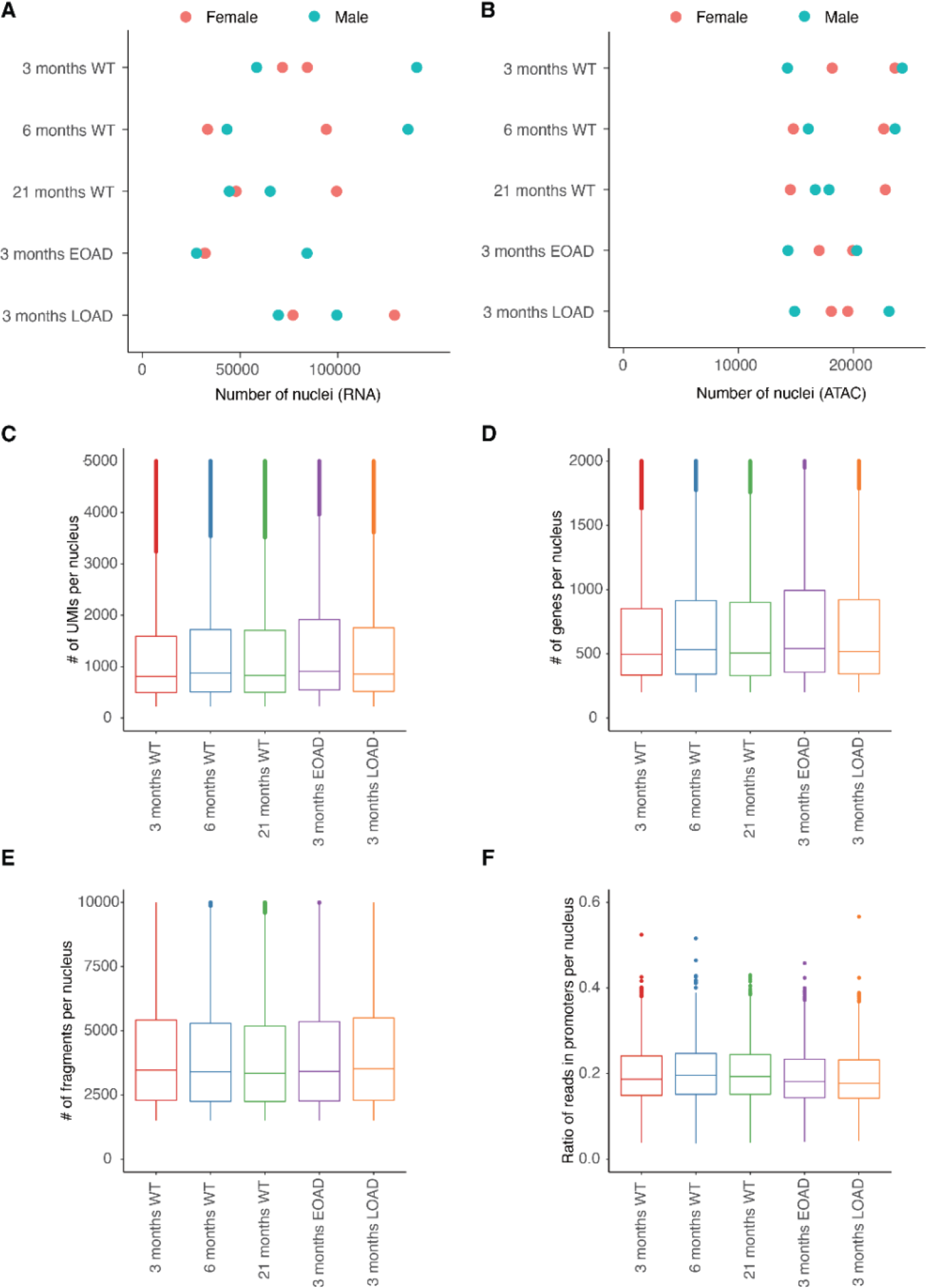
Performance of *EasySci-RNA* and *EasySci-ATAC* profiling of mouse brain samples. (A-B) Scatter plots showing the number of single-cell transcriptomes (A) and single-cell chromatin accessibility (B) profiled in each mouse individual across five conditions, colored by sex. Of note, the number of cells recovered from two mouse individuals in the EOAD model (RNA) are very close and can not be separated in the plot. (C-D) Box plots showing the number of unique transcripts (C) and genes (D) detected per nucleus in each condition profiled by *EasySci-RNA*. For all box plots: middle lines, medians; upper and lower box edges, first and third quartiles, respectively; whiskers, 1.5 times the interquartile range; and circles are outliers. (E-F) Box plots showing the number of unique fragments (E) and the ratio of reads in promoters (F) per cell in each condition profiled by *EasySci-ATAC*.

**Figure S5.**
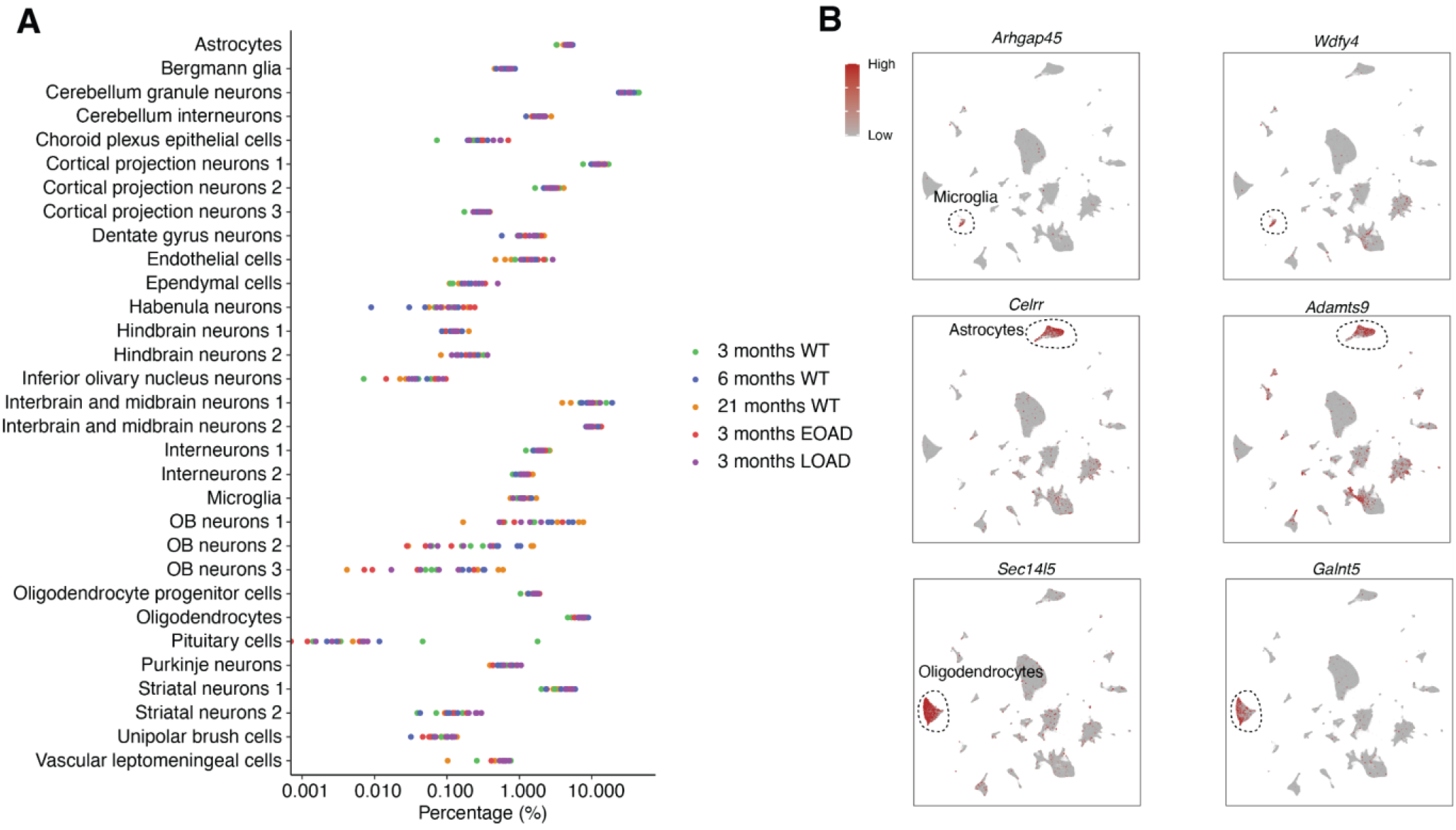
Identification of main brain cell types and cell-type-specific markers by *EasySci-RNA*. (A) The recovered cell percentage from every main cell type is shown across different replicates. The samples are color-coded based on the condition of origin. (B) UMAP plots showing the gene expression of identified novel markers for microglia (*Arhgap45*, *Wdfy4*), astrocytes (*Clerr*, *Adamts9*), and oligodendrocytes (*Sec14l5*, *Galnt5*). UMI counts for these genes are scaled by the library size, log-transformed, and then mapped to Z-scores.

**Figure S6.**
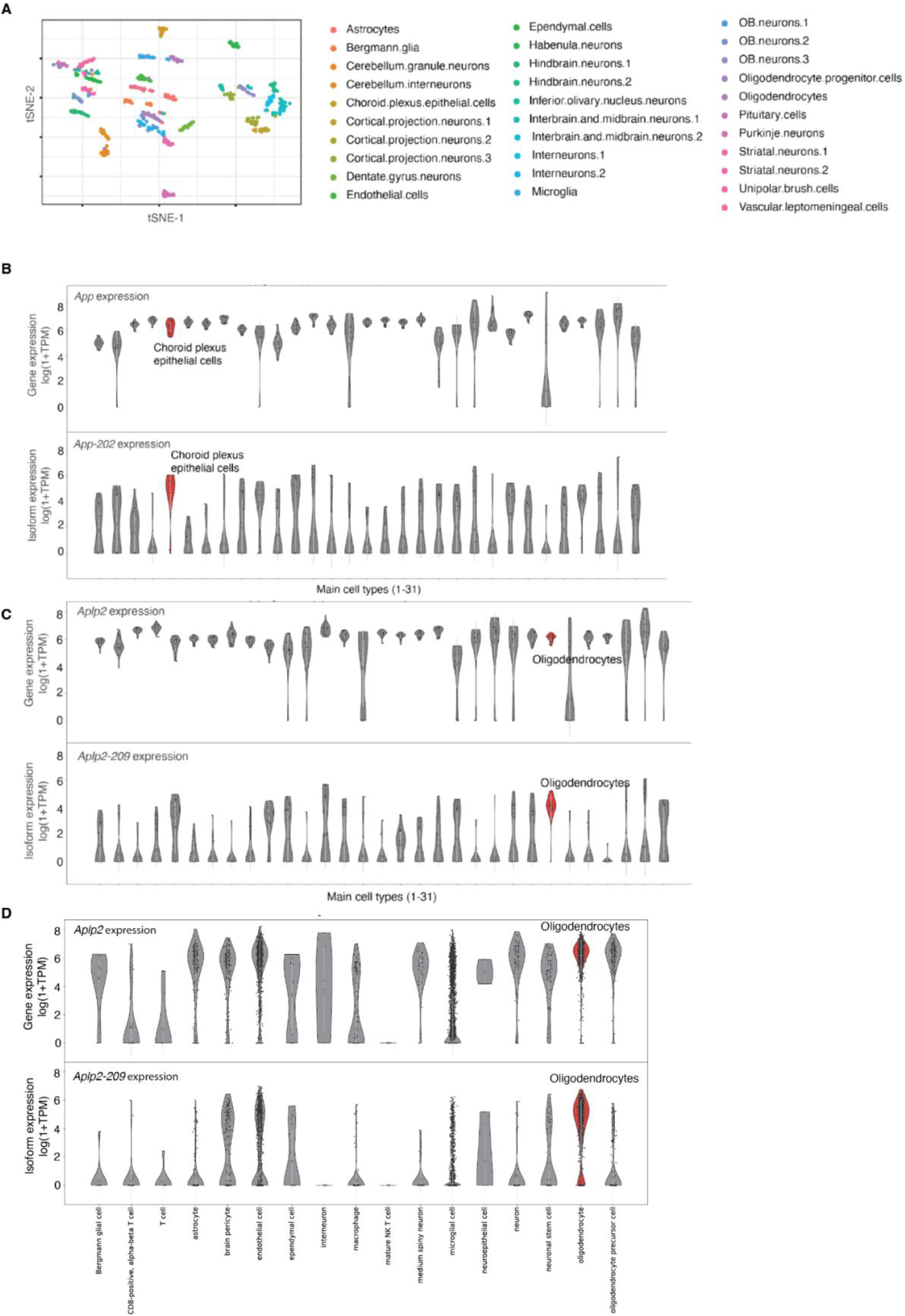
Identification of cell-type-specific isoforms in the mouse brain. (A) We aggregated randomN primed *EasySci-RNA* reads from each main cell type in every mouse individual, yielding 613 pseudocells. The t-SNE plot showed the separation of main cell types by isoform expression. (B) Violin plots showing the expression of gene *App* and isoform *App-202* across main cell types. (C-D) Violin plots showing the expression of gene *Aplp2* and isoform *Aplp2-209* across main cell types in the *EasySci* dataset (C) and the Tabula Muris Senis mouse aging atlas dataset ^7^ (D). White circles represent the normalized expression of genes and isoforms (log(1+TPM)). White bars represent the standard deviation.

**Figure S7.**
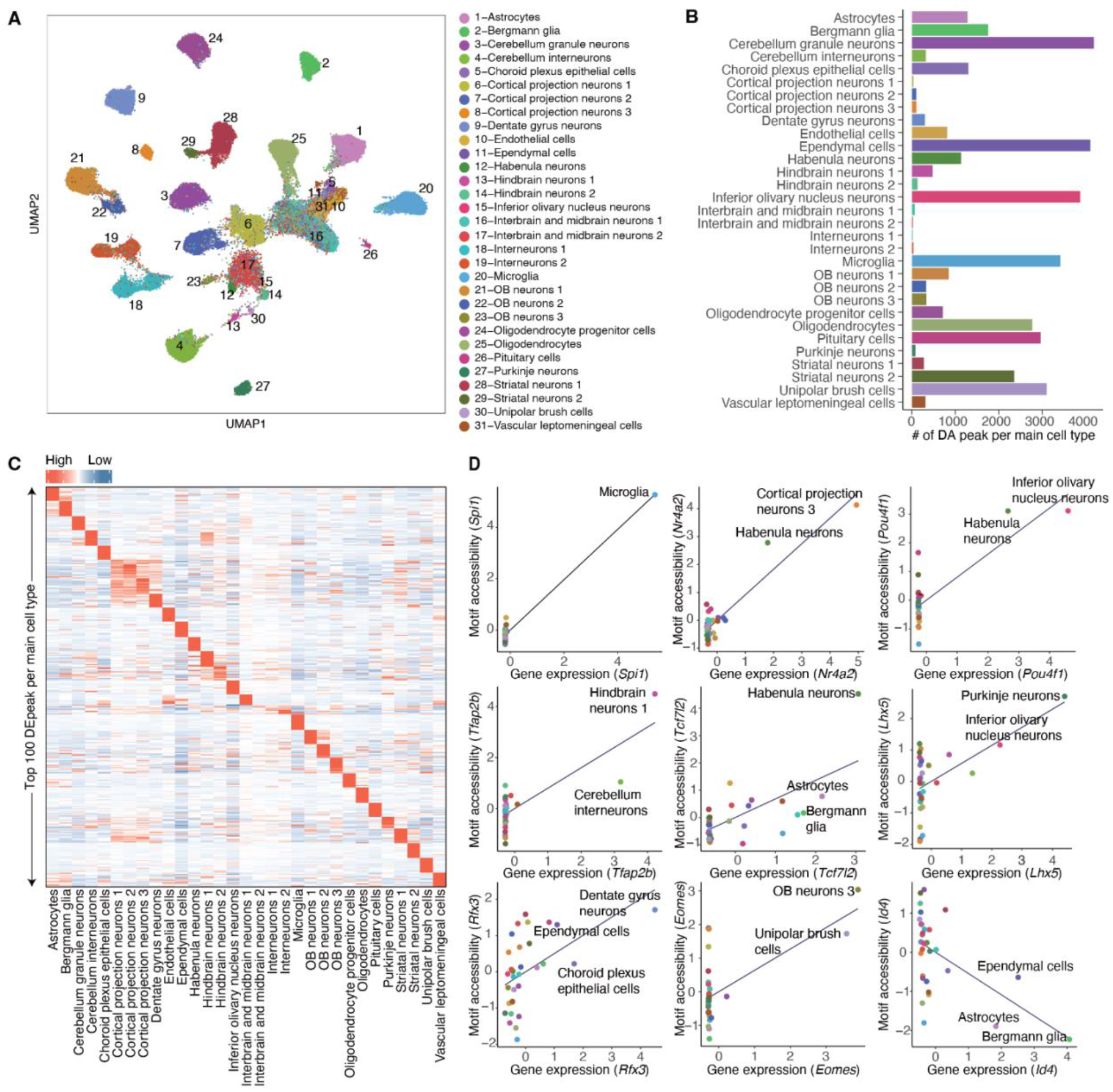
Characterization of cell-type-specific chromatin accessibility and key TF regulators using *EasySci-ATAC*. (A) UMAP plot of the *EasySci-ATAC* dataset subsampled to 5,000 cells per cell type (or all cells if the number of cells is less than 5,000), colored by main cell types in Figure 1H. The analysis used the peak-count matrix without integration with the *EasySci-RNA* dataset. (B) Bar plot showing the number of cell-type-specific peaks for each main cell type (defined as differential accessible (DA) peaks across main cell types with q-value < 0.05 and TPM > 20 in the target cell type). (C) Heatmap showing the aggregated accessibility of top 100 DA peaks per cell type (ranked by fold change between the maximum and the second accessible cell type). Unique counts for cell-type-specific peaks are first aggregated, normalized by the library size, and then mapped to Z-scores. (D) Scatter plots showing the correlation between gene expression and motif accessibility of cell -type specific TF regulators, together with a linear regression line. TF gene expressions are calculated by aggregating scRNA-seq gene counts for each main cluster, normalized by the library size, and then mapped to Z-scores. TF motif accessibilities are quantified by chromVar ^59^, then aggregated per main cell type and mapped to Z-scores (**Methods**).

**Figure S8.**
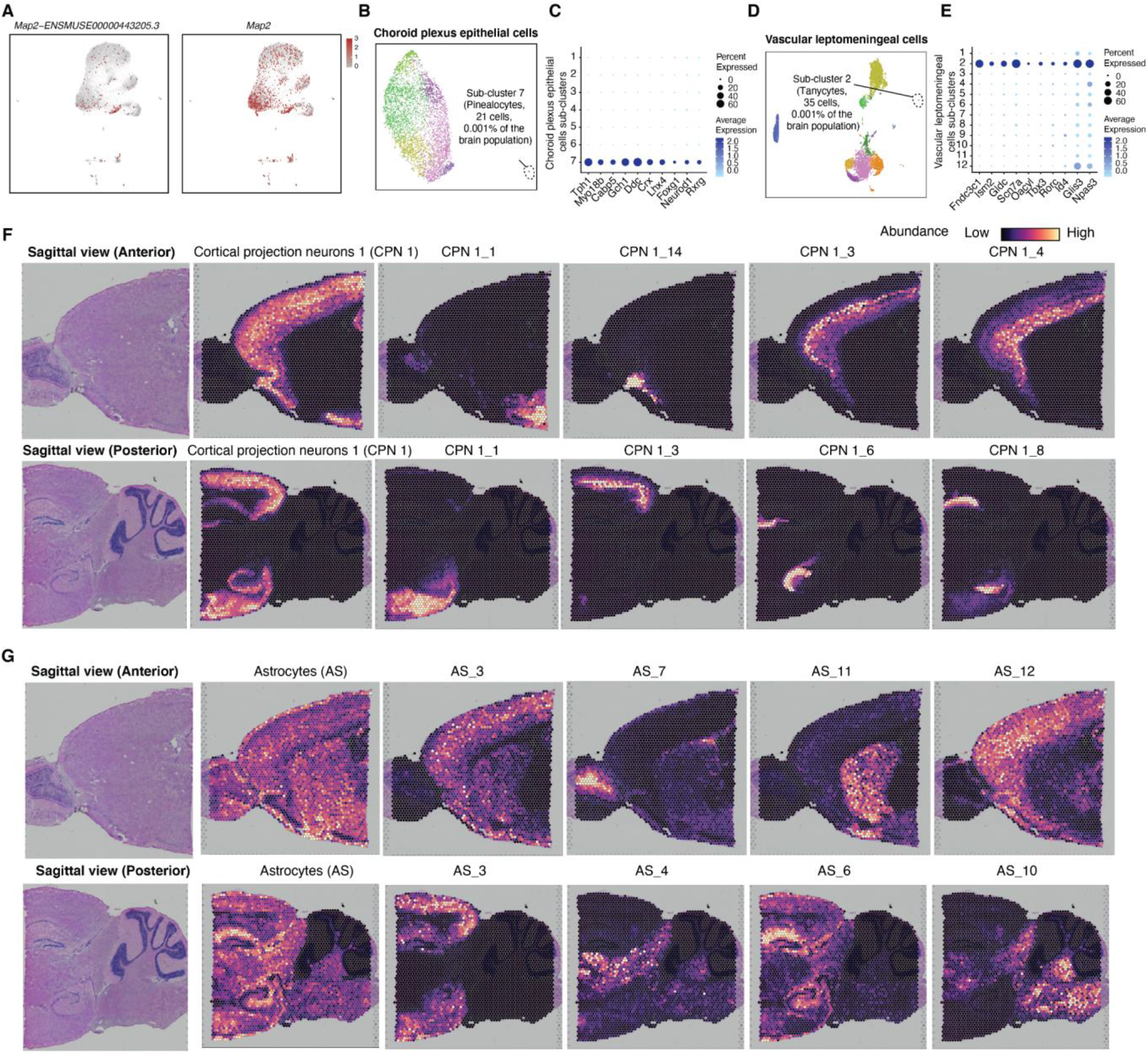
Validation of cellular subtypes in the mouse brain. (A) UMAP plots, same as Figure 2A based on both gene and exon-level expression, showing the specific expression of an example exon marker *Map2-ENSMUSE00000443205.3* (left) of microglia sub-cluster 8 and the lack of specificity of its corresponding gene *Map2* (right). Single-cell gene/exon expression was normalized first by library size, log-transformed, and then scaled to Z-scores. (B) UMAP visualizations showing sub-clustering analysis for choroid plexus epithelial cells colored by sub-cluster IDs, highlighting the rare sub-cluster shown in Figure 2C. (E) Dot plot showing the expression of selected marker genes for choroid plexus epithelial cells-7, including both normal genes (left five genes) and transcription factors (right five genes). (D) UMAP visualizations showing sub-clustering analysis for vascular leptomeningeal cells colored by sub-cluster IDs, highlighting two rare sub-clusters shown in Figure 2C. (E) Dot plot showing the expression of selected marker genes for vascular leptomeningeal cells-2, including both normal genes (left five genes) and transcription factors (right five genes). (F-G) Mouse brain sagittal sections showing spatial abundances of main cell types and related sub-clusters for cortical projection neurons 1 (F) and astrocytes (G) in anterior (top) and posterior (bottom) regions, estimated using the cell2location^35^.

**Figure S9.**
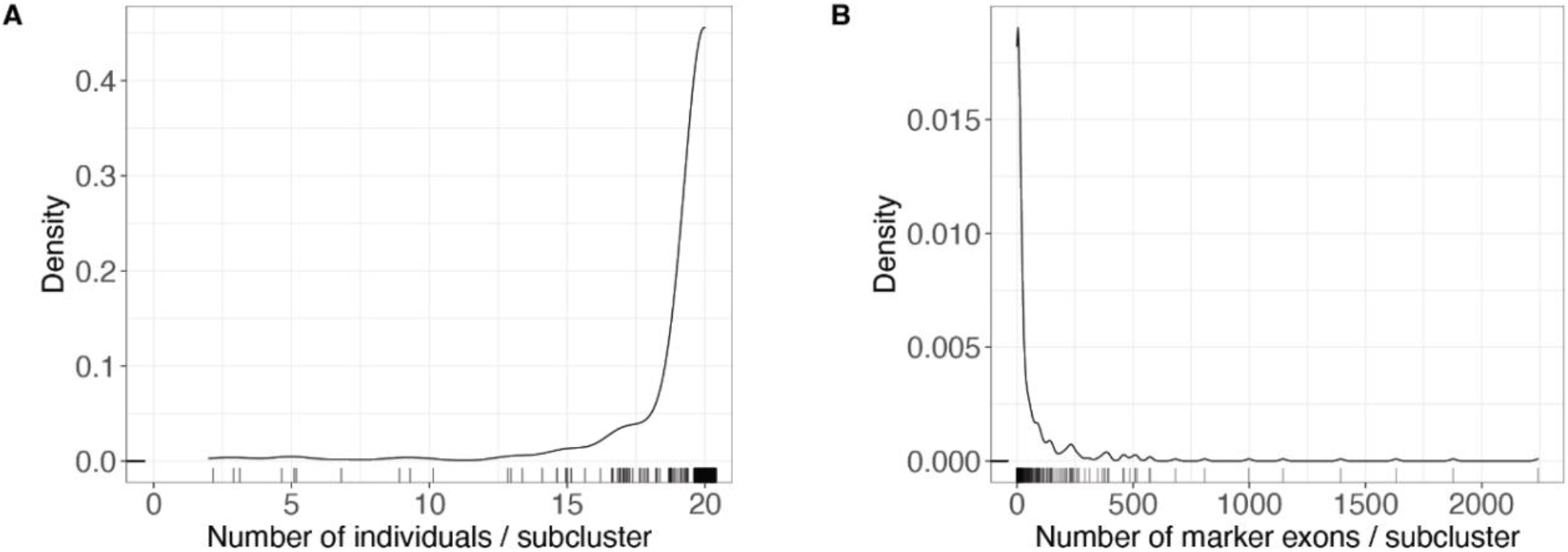
Characteristics of subclusters. (A) Density plot showing the number of individuals per subcluster. The rug plot below the density plot represents the individual subclusters. (B) Density plot of the number of marker exons per subcluster. The rug plot below the density plot represents the individual subclusters.

**Figure S10.**
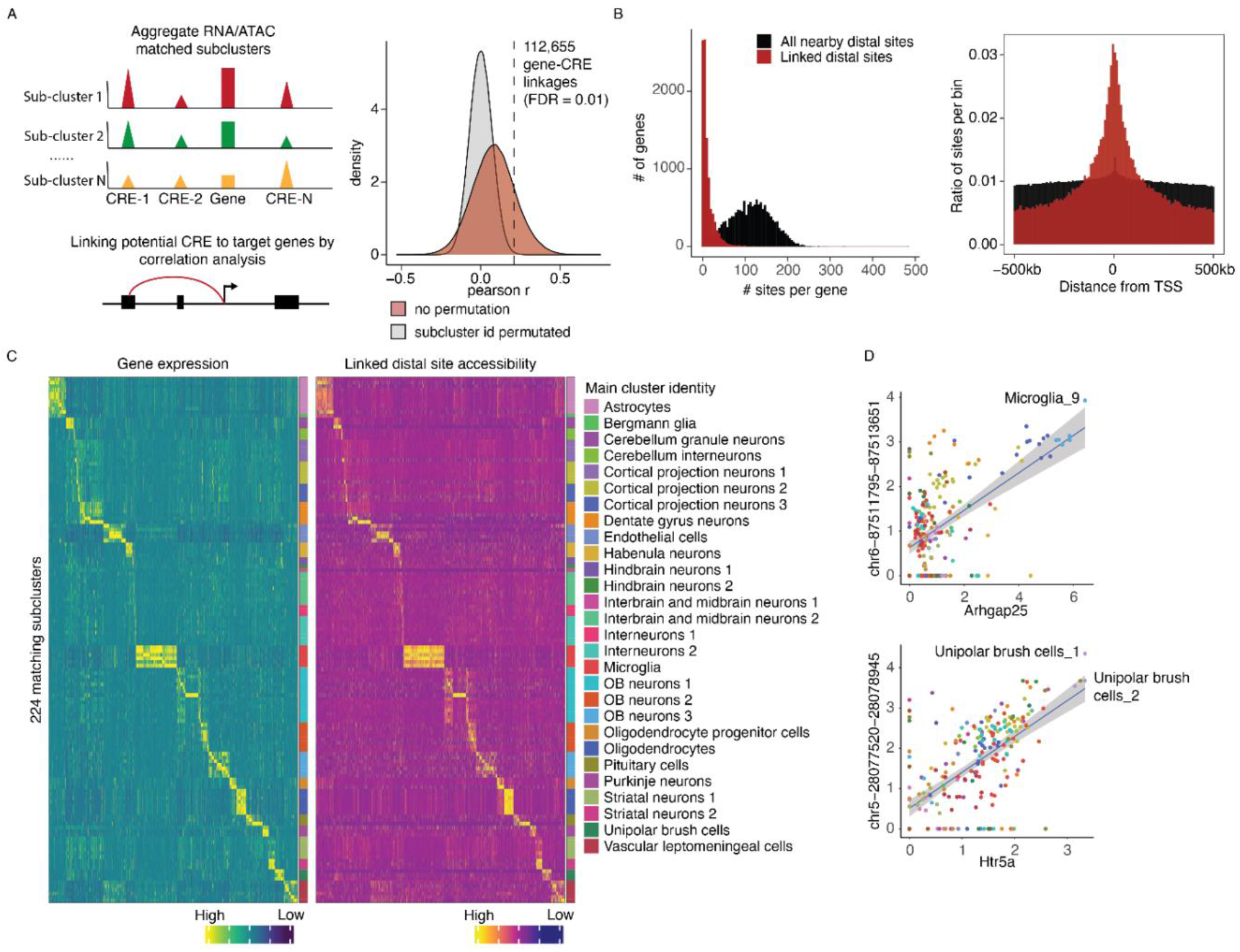
Cis-regulatory elements regulation of gene expression at the sub-cluster level. (A) Overview of analytical steps (left) and density plot showing the distribution of Pearson correlation coefficients between gene expression and the accessibility of distal linked site (colored in red) or all nearby accessible elements (within ±500 kb of the promoter, colored in gray) across pseudo-cells. Dash line indicates the Pearson correlation coefficient cutoff with 0.01 FDR. (B) Left, histogram showing the number of accessible sites per gene. Right, histogram shows the distance distribution of accessible sites within 500 kb of genes. Both plots include all nearby accessible sites (colored in black) and the linked accessible sites (colored in red). (C) Heatmaps showing the concordant expression of genes (left) and accessibility of putative distal linked sites (right) across all matched subclusters. Rows represent the aggregated gene expression or peak accessibility for a given sub-cluster. The raw aggregated RNA-seq and ATAC-seq data was normalized first by the total number of reads for each sub-cluster and then scaled to Z-score across all subtypes. (D) Correlation plot showing examples of sub-cluster-specific linkages for microglia-9 and unipolar brush cells-2.

**Figure S11.**
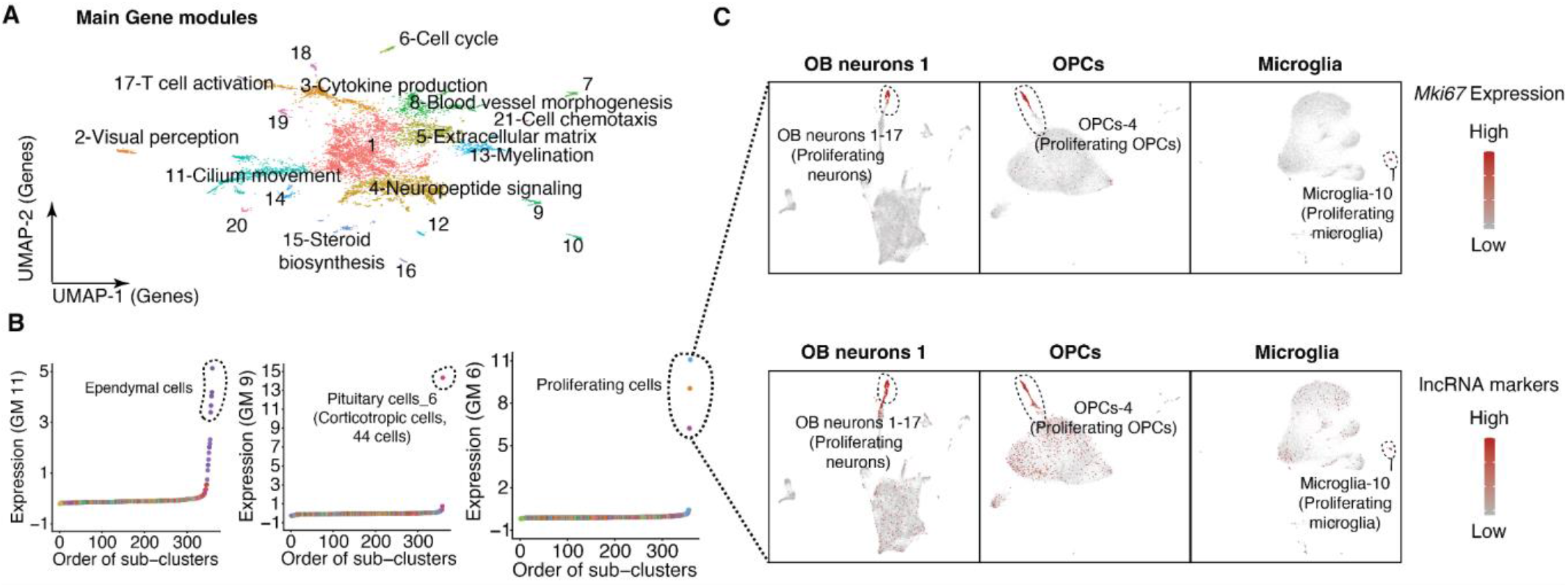
Identification of key molecular programs underlying cell type specificity in the mouse brain. (A) UMAP visualizations of genes colored by identified gene module IDs. (B) Scatter plots showing examples of gene modules and their expression levels across sub -clusters (ordered by the level of gene module expression): GM-11 is specific to ependymal cells; GM-9 is specific to pituitary cell-6 (corticotropic cells); GM-6 marks four proliferating sub-clusters from different main cell types. UMI counts for genes from each gene module are scaled for library size, log-transformed, mapped to Z-scores and then aggregated. (C) UMAP visualization showing four proliferating sub-clusters identified from OB neurons 1, astrocytes, oligodendrocyte progenitor cells, and microglia, colored by the normalized expression of canonical proliferating marker *Mki67* (top) and the aggregated expression of lncRNAs in GM-6 (bottom). UMI counts are first normalized by library size, log-transformed, and then mapped to Z-scores. OPCs, oligodendrocyte progenitor cells.

**Figure S12.**
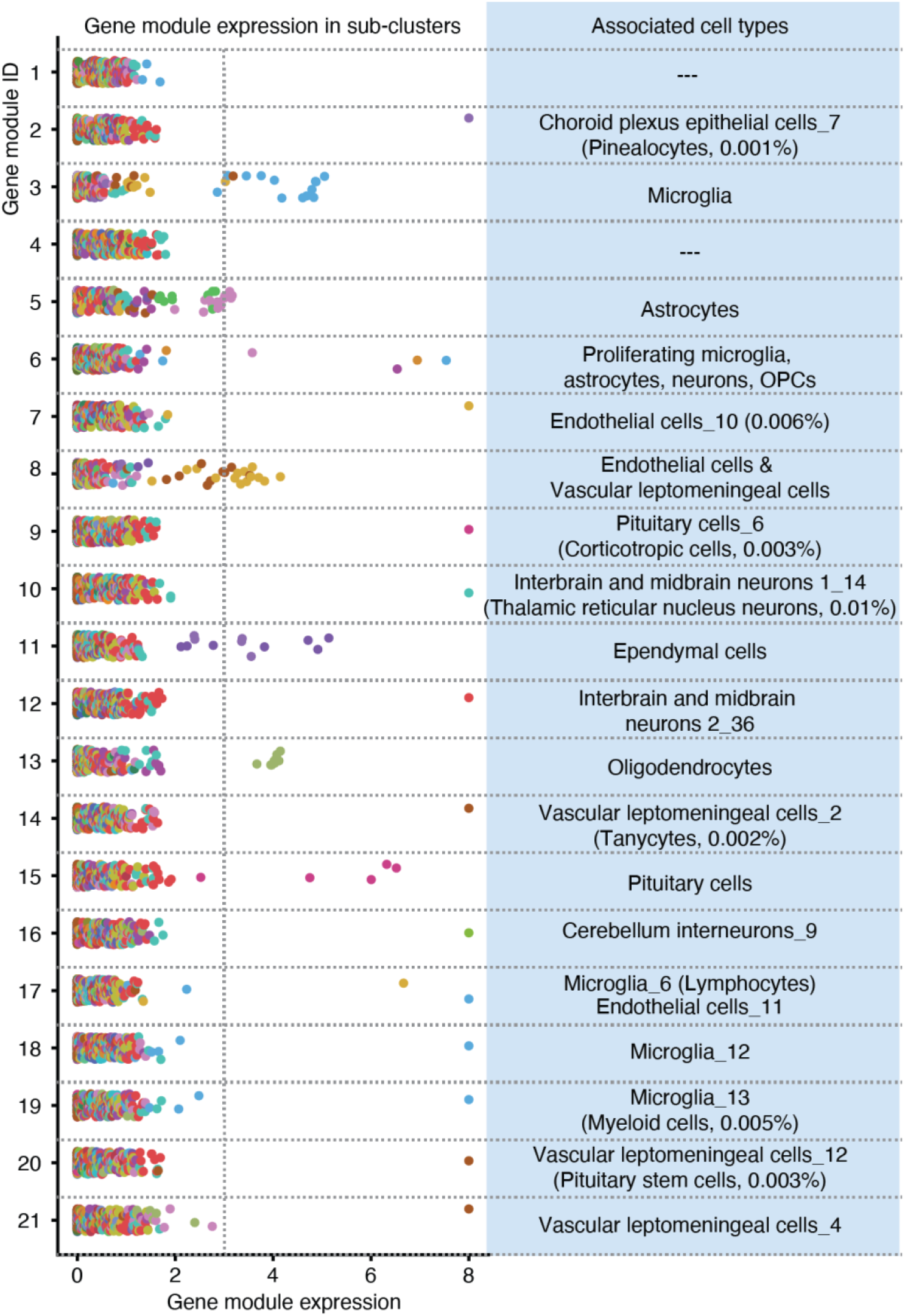
Characterization of cell types/subtypes by gene module expression. Scatter plot showing the expression of each gene module across 359 sub-clusters. The associated cell types were annotated on the plot. UMI counts for genes from each gene module are scaled for library size, log-transformed mapped to Z-scores and then aggregated.

**Figure S13.**
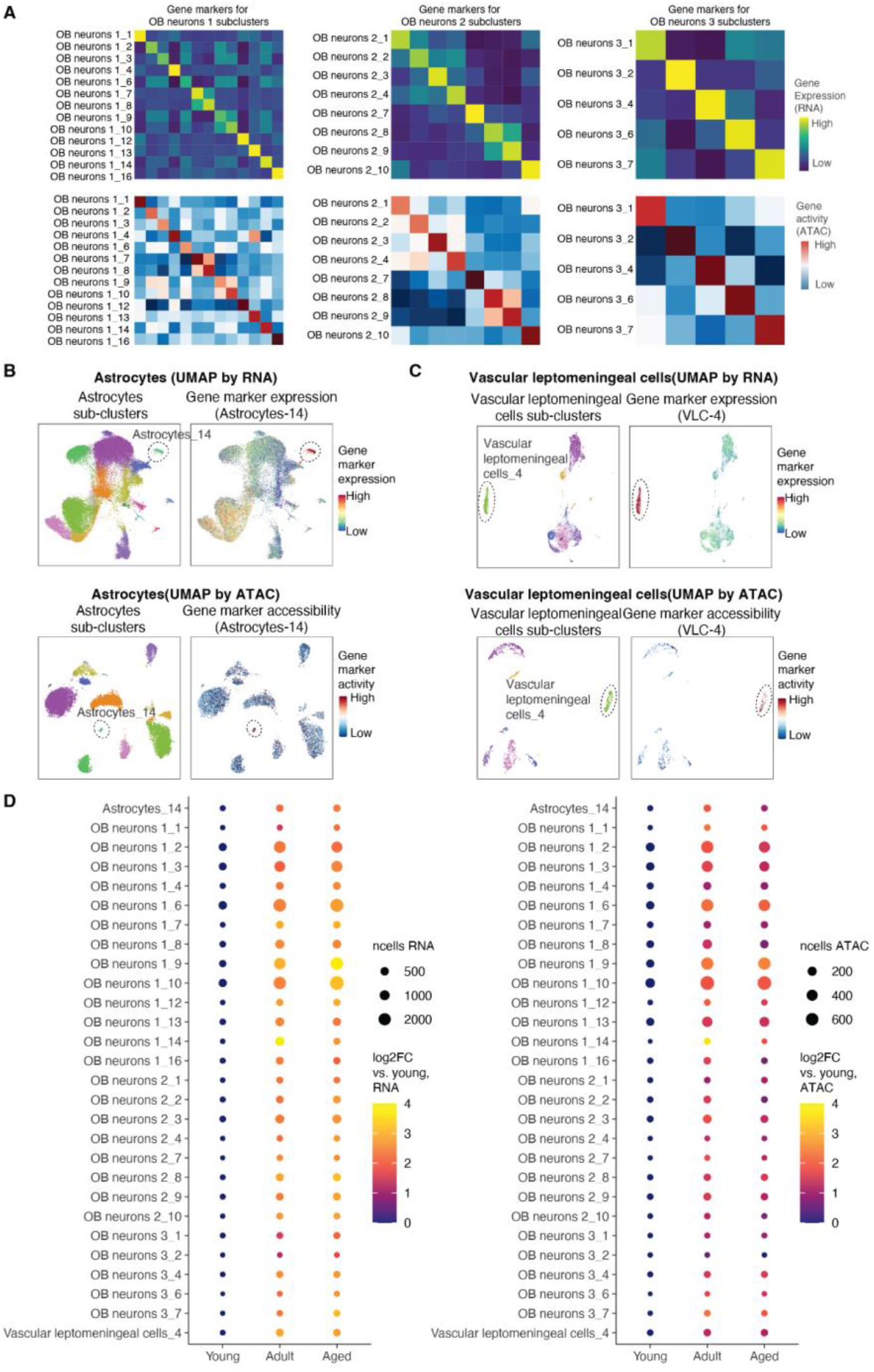
Identification of cell subtypes underlying olfactory bulb expansion from the young to adult stage in *EasySci-RNA* and *EasySci-ATAC*. (A) Heatmaps showing the aggregated gene expression (top) and gene body accessibility (bottom) of sub-cluster specific gene markers (columns) in OB expansion-associated sub-clusters (rows) from OB neurons 1 (left), OB neurons 2 (middle), and OB neurons 3 (right). UMI counts for genes or reads overlapping with gene bodies were aggregated for each sub-cluster, normalized first by the total number of reads, column centered, and scaled across all cell sub-clusters. (B-C) UMAP visualization showing astrocytes subtype 14 (B) and vascular leptomeningeal cells subtype 4 (VLC-4, C), colored by subcluster ID in *EasySci-RNA* (top left) and *EasySci-ATAC* (bottom left), the aggregated gene expression (top right) and gene body accessibility (bottom right) of sub -cluster specific gene markers. (D) For the OB expansion-related sub-clusters, we plotted their log2-transformed fold changes between each age group and the young mice, profiled by *EasySci-RNA* (left) and *EasySci-ATAC* (right).

**Figure S14.**
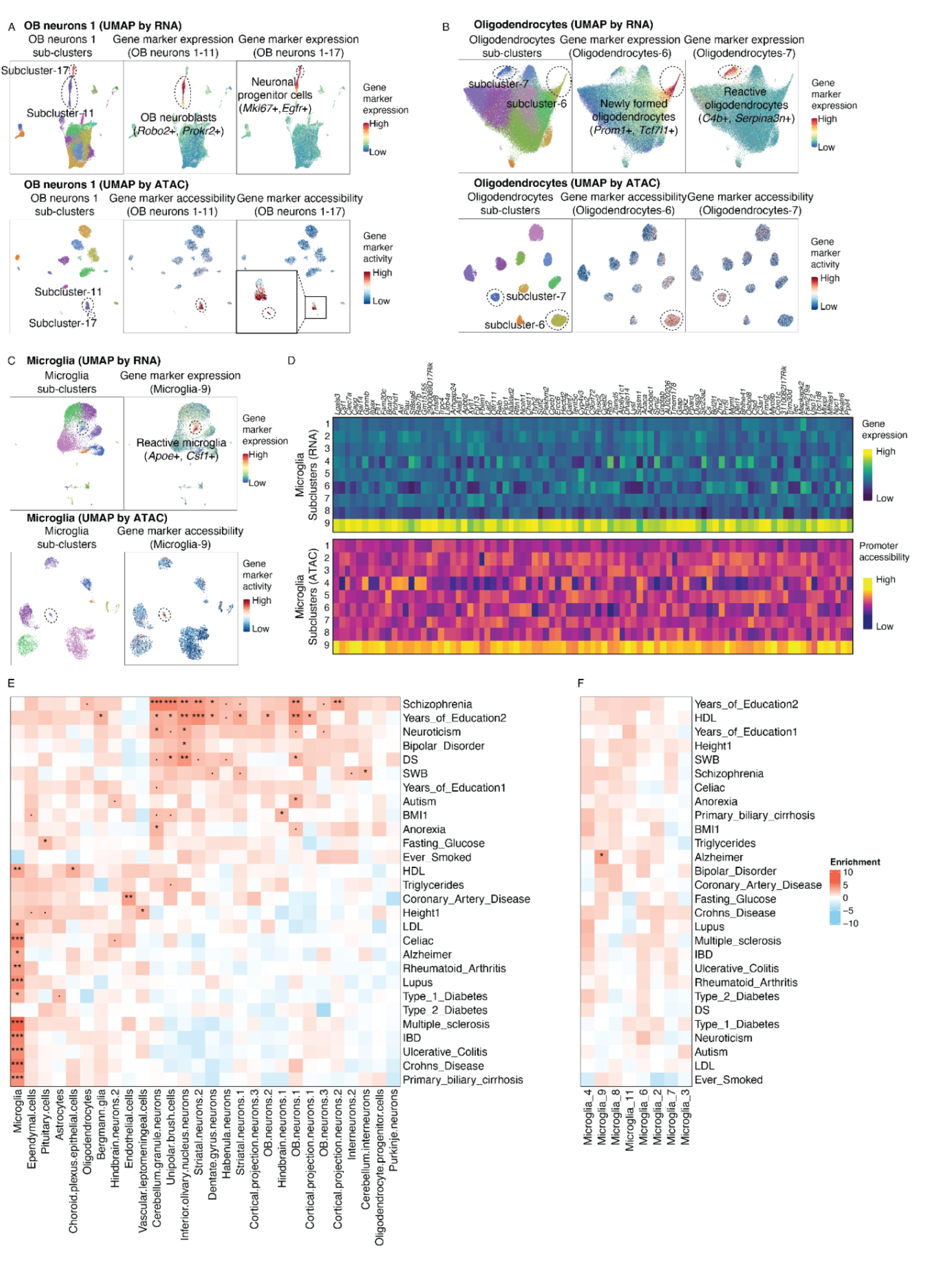
Identifying aging-associated sub-clusters in *EasySci-ATAC* and cell-type-specific enrichment of GWAS heritability of human phenotypes. (A) UMAP visualization showing OB neurons 1-11 and OB neurons 1-17 identified from *EasySci-RNA* (top) and *EasySci-ATAC* (bottom), colored by subcluster id (left), aggregated gene expression or gene activity of OB neurons 1-11 gene markers (middle) and OB neurons 1-17 gene markers (right). (B) UMAP visualization showing oligodendrocytes-6 and oligodendrocytes-7 identified from *EasySci-RNA* (top) and *EasySci-ATAC* (bottom), colored by subcluster id (left), aggregated gene expression or gene activity of oligodendrocytes-6 gene markers (middle) and oligodendrocytes-7 markers (right). (C) UMAP visualization showing microglia-9 identified from *EasySci-RNA* (top) and *EasySci-ATAC* (bottom), colored by subcluster id (left), aggregated gene expression or gene activity of microglia-9 gene markers (right). Subcluster marker genes were identified by differential expression analysis using scRNA-seq data (**Methods**). (D) Heatmap showing the gene expression (top) and the promoter accessibility (bottom) of microglia-9 enriched genes across subclusters. The *EasySci-RNA* data (UMI count matrix) and *EasySci-ATAC* data (read count matrix) were aggregated per sub-cluster, normalized by the total number of reads, column centered, and scaled. Of note, rare subclusters from RNA-seq data that were not detected in ATAC-seq data were not included in this analysis. (E) Heatmap showing the results of LDSC analysis of the SNPs associated with the indicated phenotypes in DEpeaks across main cell types, colored by z score of regression coefficient. *FDR < 0.05, **FDR < 0.01, ***FDR < 0.001. Only cell types that contain DEpeaks from all autosomes are included in this analysis. (F) Heatmap showing the results of LDSC analysis of the SNPs associated with the indicated phenotypes in DEpeaks across microglia subtypes, colored by z score of regression coefficient. *FDR < 0.05, **FDR < 0.01, ***FDR < 0.001. Only subtypes that contain DEpeaks from all autosomes are included in this analysis.

**Figure S15.**
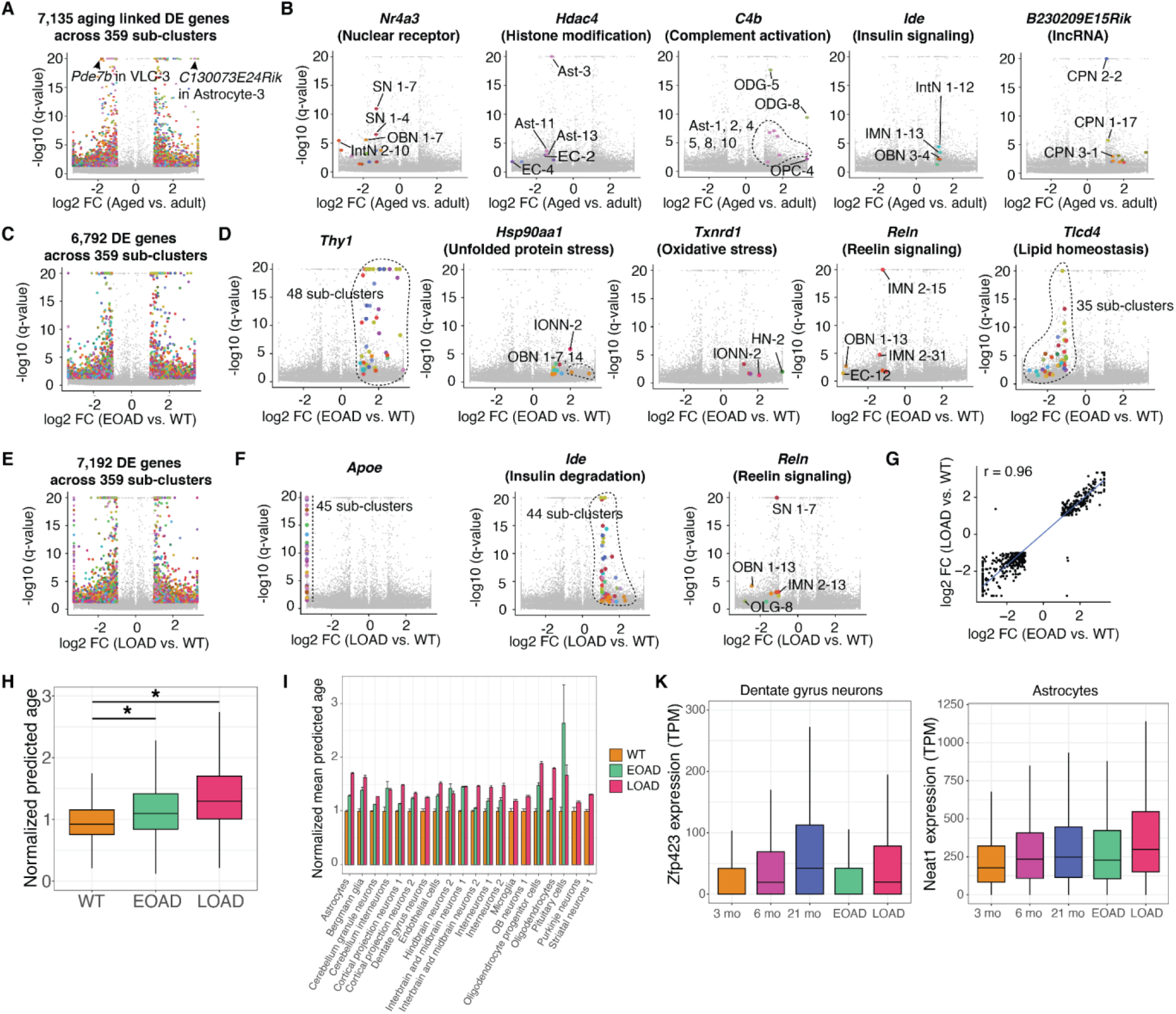
Identifying aging and AD pathogenesis-associated gene expression signatures. (A) Volcano plots showing the differentially expressed (DE) genes between adult (6 months) and aged (21 months) mice across all sub-clusters. Significantly changed genes are colored by the main cell type identity for the corresponding sub-cluster. (B) Volcano plot same as (A), highlighting example DE genes with concordant changes across multiple sub-clusters comparing adult and aged models, labeled with related biological pathways. (C) Volcano plots showing the differentially expressed (DE) genes between WT and EOAD models across all sub-clusters. Significantly changed genes are colored by the main cell type identity for the corresponding sub-cluster. (D) Volcano plot same as (C), highlighting example DE genes with concordant changes across multiple sub-clusters comparing WT and EOAD models, labeled with related biological pathways. (E) Volcano plots showing the differentially expressed (DE) genes between WT and LOAD models across all sub-clusters. Significantly changed genes are colored by the main cell type identity for the corresponding sub-cluster. (F) Volcano plot same as (E), highlighting example DE genes with concordant changes across multiple sub-clusters comparing WT and LOAD models, labeled with related biological pathways. (G) We detected 559 DE genes significantly changed within the same sub-cluster in both AD models (both compared with the wild-type). The scatter plot shows the correlation of the log2-transformed fold changes of these 559 shared DE genes in the EOAD model (x-axis) and the LOAD model (y-axis). (H) Boxplot displays the predicted biological age of 3-month-old wild-type mice against AD models across various cell types, normalized by the mean predicted age of the WT samples. A ridge regression model was employed to predict the ln(age) of pseudobulk cells (averaging 15 cells) utilizing 80% of cells from 3, 6, and 21-month-old mice. Predicted ages were subsequently calculated for the remaining 20% of WT mice and the entirety of the AD models, with individual models crafted for each cell type. (I) Barplot (supplemented with standard error) illustrates the mean predicted age of 3-month-old wild-type mice against AD models, focusing only on cell types and conditions demonstrating significant differences (Q-value < 0.05, Mann-Whitney U test). The values are normalized by the mean predicted age of the WT samples. (K) Boxplots represent the expression of the Zfp423 gene in Dentate gyrus neurons and the Neat1 gene in astrocytes across various conditions. Both genes, ranking among the top predictors of age in their respective cell types, exhibited increased expression in the 3-month-old AD models (Zfp423 solely in the LOAD model, Neat1 in both AD models).

**Figure S16.**
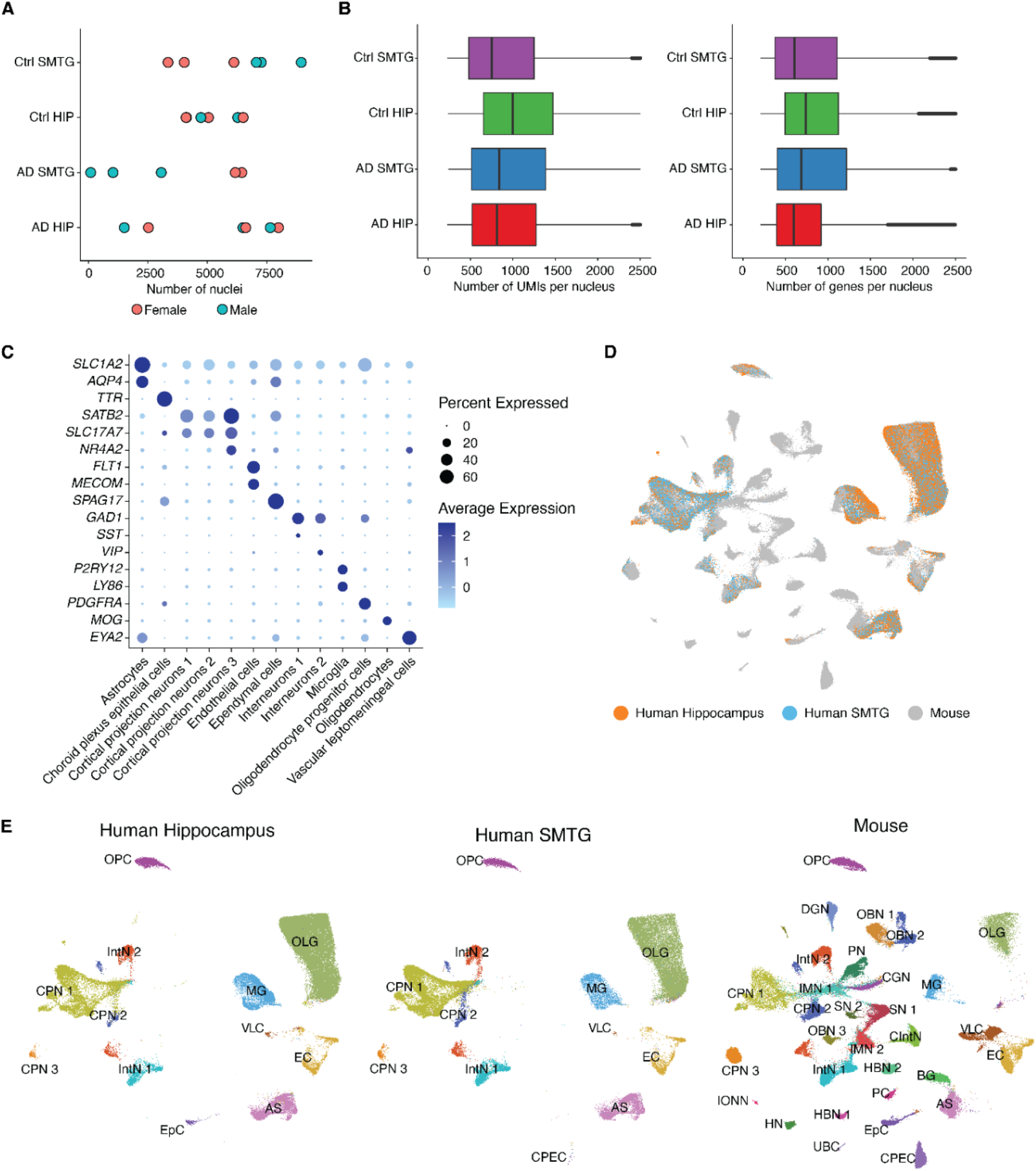
Performance and quality control of the human brain dataset. (A) Scatter plot showing the number of single-cell transcriptomes profiled in each human sample in two regions, colored by sexes. Of note, the number of cells recovered from two AD individuals in the SMTG are very close and cannot be separated in the plot. (B) Box plots showing the number of unique transcripts (left) and genes (right) detected per nucleus profiled in the human dataset. For all box plots: middle lines, medians; upper and lower box edges, first and third quartiles, respectively; whiskers, 1.5 times the interquartile range; and circles are outliers. (C) Dotplot showing the markers for the main cell types identified in the human dataset. (D-E) UMAP plot showing the integration between human and mouse cells, colored by the dataset (D) and main cell types (E). AS, astrocytes; BG, Bergmann glia; CGN, cerebellum granule neurons; CIntN, cerebellum interneurons; CPEC, choroid plexus epithelial cells; CPN 1, cortical projection neurons 1; CPN 2, cortical projection neurons 2; CPN 3, cortical projection neurons 3; DGN, dentate gyrus neurons; EC, endothelial cells; EpC, ependymal cells; HN, habenula neurons; HBN 1, hindbrain neurons 1; HBN 2, hindbrain neurons 2; IONN, Inferior olivary nucleus neurons; IMN 1, interbrain and midbrain neurons 1; IMN 2, interbrain and midbrain neurons 2; IntN1, interneurons 1; IntN2, interneurons 2; MG, microglia; OBN 1, OB neurons 1; OBN 2, OB neurons 2; OBN 3, OB neurons 3; OPC, oligodendrocyte progenitor cells; OLG, oligodendrocytes; PC, pituitary cells; PN, purkinje cells; SN 1, striatal neurons 1; SN 2, striatal neurons 2; UBC, unipolar brush cells; VLC, vascular leptomeningeal cells.

**Figure S17.**
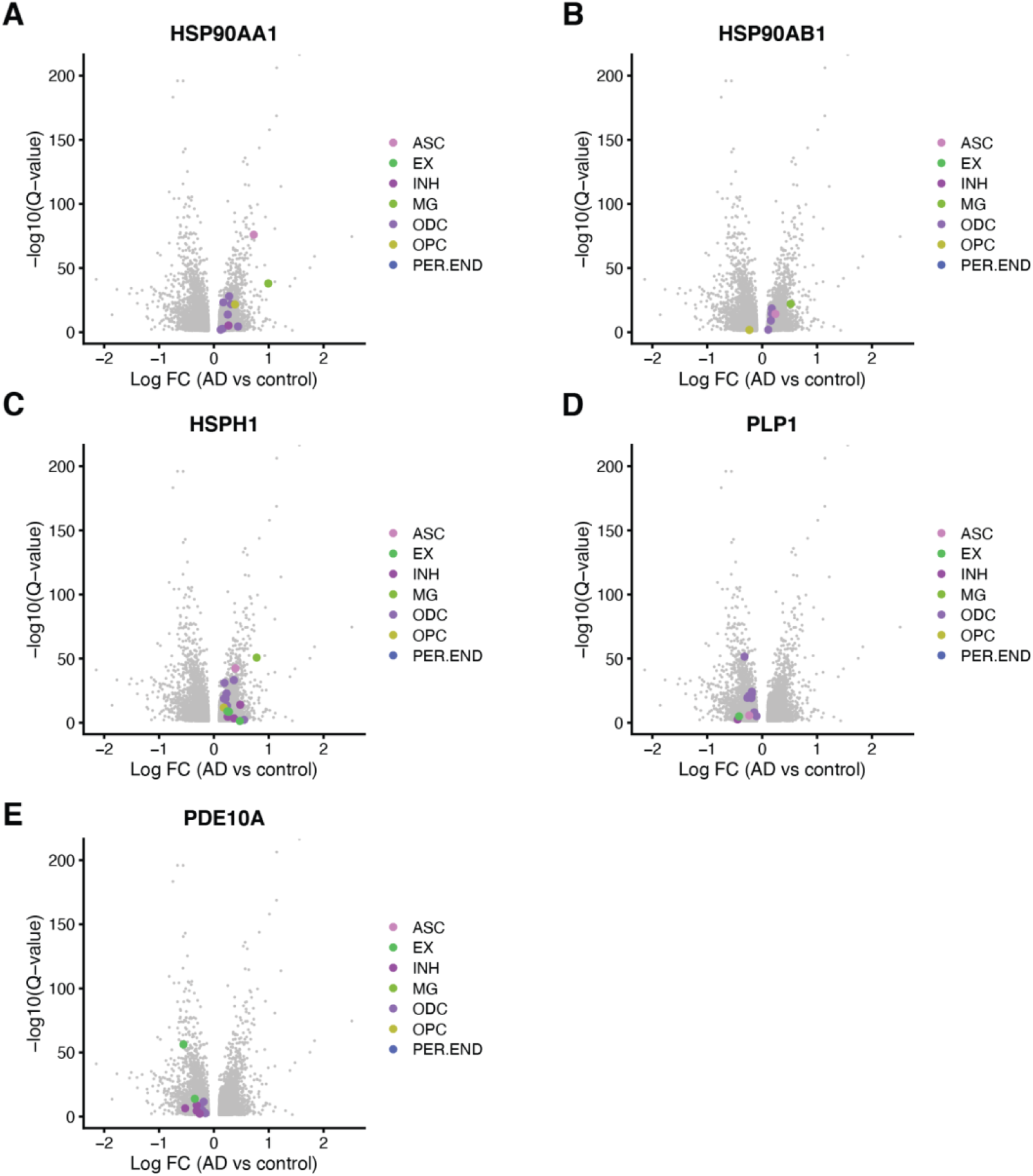
Identifying conserved gene expression changes across mouse AD models and human AD samples from the prefrontal cortex. (A-E) Volcano plots of genes from human prefrontal cortex samples ^6^. These genes show consistent changes in multiple cell subclusters between mouse AD models, human hippocampus and SMTG samples, and human prefrontal cortex samples from the above-mentioned publication. Dots on the figures represent genes across cell subclusters; abbreviations correspond to the following cell types: ASC, astrocytes; EX, excitatory neurons; INH, inhibitory neurons; MG, microglia; ODG, oligodendrocytes; OPC, oligodendrocyte progenitor cells; and PER.END, pericyte/endothelial cells.

## Materials and Methods

### Animals

C57BL/6 wild-type mouse brains at three months (n=4), six months (n=4), and twenty-one months (n=4) were collected in this study. These age points correspond to approximately 20, 30, and 62 years in humans. Furthermore, to gain insight into the early cellular state changes underlying the pathophysiology of Alzheimer’s disease, we added two AD models at 3-month-old from the same C57BL/6 background. These include an early-onset AD model (5xFAD, JAX stock #034840) that overexpresses mutant human amyloid-beta precursor protein (APP) with the Swedish (K670N, M671L), Florida (I716V), and London (V717I) Familial Alzheimer’s Disease (FAD) mutations and human presenilin 1 (PS1) harboring two FAD mutations, M146L and L286V. Brain-specific overexpression is achieved by neural-specific elements of the mouse *Thy1* promoter ^17^. The second, late-onset AD model (APOE*4/Trem2*R47H, JAX stock #028709) in this study carries two of the highest risk factor mutations of LOAD ^88^, including a humanized *APOE* knock-in allele, where exons 2, 3, and most of exon 4 of the mouse gene were replaced by the human ortholog including exons 2, 3, 4 and some part of the 3’ UTR. Furthermore, a knock-in missense point mutation in the mouse *Trem2* gene was also introduced, consisting of an R47H mutation, along with two other silent mutations. Two male and two female mice are included in each condition.

By studying 3-month-old animals, our goal was to gain insight into the early changes underlying the pathophysiology of the AD models. Mature adult mice start at the age of 3 months, but multiple AD hallmarks, including amyloid-beta plaques and gliosis, can be observed in the early-onset 5xFAD model^17^. Therefore, we decided that this age might be the most appropriate for our goal to study early contributors of Alzheimer’s disease pathomechanism.

### *EasySci-RNA* library preparation and sequencing

Extracted mouse brains were snap-frozen in liquid nitrogen and stored at -80°C. Detailed step-by-step *EasySci-RNA* protocol is included as a supplementary file (**Supplementary file 1**).

#### Initial Oligo Cost

For reverse transcription, we used two 96-well plate barcoded primers, one with shortdT and the other with random hexamers. Each well contains 700 uL of 100 uM primer, and each plate costs $2,756 and $2,445, respectively. For ligation, we used one plate of barcoded ligation oligos with 250 uL of 100 uM in each well, costing $518. For PCR, we used one plate of barcoded PCR primers with 200 uL of 100 uM in each well, costing $470. For a one million single-cell transcriptome profiling experiment, we used a total of four plates for reverse transcription, four plates for ligation, and two plates for PCR. Therefore, the total cost of the initial oligo purchase is $13,414.

### Human brain sample

Twenty-four post-mortem human brain samples across two regions (hippocampus and superior and middle temporal gyrus) and twelve individuals, including six controls and six Alzheimer’s disease patients ranging from 70 to 94 in age, were collected from the University of Kentucky Alzheimer’s Disease Center Tissue Bank. Each surveyed sample underwent rigorous quality control, including short PMI, and was subjected to *EasySci-RNA* profiling. The libraries were sequenced across four Illumina NextSeq™ 1000 sequencer runs.

### Computational procedures for processing *EasySci-RNA* libraries

A custom computational pipeline was developed to process the raw fastq files from the *EasySci* libraries. Similar to our previous studies ^10,11^, the barcodes of each read pair were extracted. Both adaptor and barcode sequences were trimmed from the reads. Second, an extra trimming step is implemented using Trim Galore ^91^ with default settings to remove the poly(A) sequences and the low-quality base calls from the cDNA. Afterward, the paired-end sequences were aligned to the genome with the STAR aligner ^92^, and the PCR duplicates were removed based on the UMI sequence and the alignment location. Finally, the reads are split into SAM files per cell, and the gene expression is counted using a custom script. At this level, the reads from the same cell originating from the short dT and the random hexamer RT primers were counted as independent cells. During the gene counting step, we assigned reads to genes if the aligned coordinates overlapped with the gene locations on the genome. If a read was ambiguous between genes and derived from the short dT RT primer, we assigned the read to the gene with the closest 3’ end; otherwise, the reads were labeled as ambiguous and not counted. If no gene was found during this step, we then searched for candidate genes 1000 bp upstream of the read or genes on the opposite strand. Reads without any overlapped genes were discarded.

We used a similar strategy to generate an exon count matrix across cells. Specifically, we counted the number of expressed exons based on the number of reads overlapping each exon. If one read overlapped with multiple exons, this read was split between the exons. Read overlapped with multiple genes were discarded, except if we can determine the exact gene based on the other paired-end read. For reads without overlapped genes, we checked if there are any overlapped exons on the opposite strand. Reads without any overlapped exons were discarded.

To compare the performance and library purity of *EasySci-RNA* with the commercial 10x Chromium system on mouse brain samples (number of UMI per nucleus and fraction of cell-associated unique reads), we subsampled ∼3,800 raw reads/cell from one randomly selected PCR batch of our large-scale mouse brain experiment and from the following 10x Chromium dataset^14^. After the subsampling, the *EasySci* data was processed with the custom computational pipeline, while the 10x Chromium data was processed with 10x Genomics’ Cell Ranger software^93^. We used the following criteria to remove low-quality cells: (i) the percentage of unassigned reads > 30%, (ii) the number of UMIs > 20,000, and (iii) the detected number of genes < 200. To compare the reaction efficiency of the two methods in high-quality cells, we first selected the top 1000 highest-quality cells from the 10x Chromium dataset^14^ and a deeply sequenced EasySci-RNA library profiling adult mouse brains. Subsequently, we subsampled these cells to different sequencing depths and quantified the unique transcripts/genes detected per cells.

Based on this comparison, we recommend a sequencing depth of no less than 5,000 raw reads per nucleus to ensure adequate coverage and detection of a substantial number of unique molecules (> 1,000 UMIs per cell) when conducting EasySci-RNA profiling. It should be noted that these are general guidelines, and the actual number of UMIs per cell may vary depending on specific experimental conditions. Moreover, other factors, such as the integrity of the cells and the method of cell fixation, should be considered when determining the ideal sequencing depth for a particular single-cell RNA sequencing experiment.

### Cell clustering and cell type annotation of single-cell RNA-seq data

After gene counting, we kept the cells with reads identified by both RT primers. We then merged the reads from the same cells. Low-quality cells were removed based on one of the following criteria: (i) the percentage of unassigned reads > 30%, (ii) the number of UMIs > 20,000, and (iii) the detected number of genes < 200. We then used the Scrublet ^94^ computational pipeline to identify and remove potential doublets, similar to our previous study ^10^. At the end of these filtering steps, we had around 1.5 million brain cells in the dataset.

To identify distinct clusters of cells corresponding to different cell types, we subjected the 1,469,111 single-cell gene expression profiles to UMAP visualization and Louvain clustering, similar to our previous study ^10^. We then co-embedded our data with the published datasets ^2,95,96^ through Seurat ^97^, and clusters were annotated based on overlapped cell types. The annotations were manually verified and refined based on marker genes. Differentially expressed genes across cell types were identified with the differentialGeneTest() function of Monocle 2 ^98^. To identify cell type-specific gene markers, we selected genes that were differentially expressed across different cell types (FDR of 5%, likelihood) and also with a > 2-fold expression difference between first and second-ranked cell types and TPM > 50 in the first-ranked cell types.

### Isoform expression analysis

Isoform expression was quantified in *EasySci* data using an adapted version of the pipeline built by Booeshaghi et al. ^22^ RandomN-primed reads for ∼1.5M single cells were merged into 613 pseudocells, grouping by individual mouse and cell types (31 cell types). The pseudocells were aligned to the mouse transcriptome with *kallisto* ^99^, generating a raw isoform count matrix. To filter and pre-process the raw data, isoform counts were normalized by length, and genes and isoforms with a dispersion of less than 0.001 were removed. The gene count matrix was produced by aggregating counts of all isoforms of a given gene. Both isoform and gene count matrices were normalized by dividing the counts in each cell by the sum of the counts for that cell, then multiplying by 1,000,000 and transforming with numpy’s log1p() function. The filtered data contained 33,361 isoforms corresponding to 12,636 genes. Highly variable isoforms and genes were identified using scanpy, by binning into 20 bins and scaling the dispersion for each feature to zero mean and unit variance within each bin. The top 5,000 gene and isoforms in each matrix were retained based on normalized dispersion. Neighborhood components analysis was performed on the filtered and normalized isoform matrix after scaling the log(1+TPM) expression to zero mean and unit variance, training on cell type labels from each pseudocell with random state 42, and visualized using t-SNE with perplexity 10, 5,000 iterations and random state 42. Differentially expressed isoforms were identified by looking for isoforms that were upregulated across a given cell type, while the genes containing those isoforms were not significantly expressed more among that cell type than its complement (the rest of the dataset). Isoforms expressed in less than 90% of pseudocells within a cell type were discarded. T-tests used a significance level of 0.01 with Bonferroni correction for multiple comparisons.

### Sub-cluster analysis of the single-cell RNA-seq data

To identify cell subtypes, we selected each main cell type and applied PCA, UMAP and Louvain clustering similarly to the major cluster analysis, based on a combined matrix including the 30 principal components derived from the gene-level expression matrix and the first 10 principal components derived from the exon-level expression matrix. We then merged sub-clusters that were not readily distinguishable in the UMAP space through an intra-dataset cross-validation procedure described before ^10^. A total of 359 cell subtypes were identified, with a median of 1,038 cells in each group. All subtypes were contributed by at least two individuals (median of twenty). Differentially expressed genes and exons across cell types were identified with the differentialGeneTest() function of Monocle 2 ^98^. To identify sub-cluster-specific differentially expressed genes associated with aging or AD models, we sampled a maximum of 5,000 cells per condition for downstream DE gene analysis using the differentialGeneTest function of the Monocle 2 package ^98^. The sex of the animals was included as a covariate to reduce sex-specific batch effects.

To detect cellular fraction changes at the subtype level across various conditions, we first generated a cell count matrix by computing the number of cells from every sub-cluster in each reverse transcription well profiled by *EasySci-RNA*. Each RT well was regarded as a replicate comprising cells from a specific mouse individual. We then applied the likelihood-ratio test to identify significantly changed sub-clusters between different conditions, with the differentialGeneTest() function of Monocle 2 ^98^. Sub-clusters were removed if they had less than 20 cells in either the male or female samples. The fold change was calculated manually by first normalizing the number of cells in a cluster by the total number of cells in the corresponding condition, then dividing the normalized values in the case and control conditions after adding a small number (10^-5^) to reduce the effect of the very small clusters. In addition, we considered subclusters to change significantly only if there was at least a two-fold change between two groups and the q-value was less than 0.05.

### Spatial transcriptomic analysis to estimate spatial abundances of cell subtypes

To spatially map *EasySci* cell subtypes, we integrated the *EasySci-RNA* data with a publicly available 10x Visium spatial transcriptomics dataset ^28–30^ using cell2location, a Bayesian model designed to map fine-grained cell types. We first aggregated ∼50 single-cell transcriptomes identified by k-means clustering (k = 50) of cells in the UMAP space of sub-clustering analysis. The cell2location model first used negative binomial regression to estimate reference cell type signatures from the *EasySci-RNA* data. In a second step, cell2location decomposed spatial mRNA counts from 10x Visium data into the reference signatures to estimate cell type spatial abundances. Training of the model utilized durations of 25 and 15,000 epochs for the negative binomial regression and spatial mapping steps, respectively. We repeated this analysis on the dataset with 75 coronal sections ^36^. To establish the corresponding regions of EasySci subclusters, we utilized the regional annotation of the spatial pixels and we manually reviewed the anatomical regions of the top 10 pixels with the highest mapping score. Only mapping scores above 1 were considered to remove low quality spatial mappings.

### Gene module analysis

We performed gene module analysis to identify the molecular programs underlying different cell types in the brain. First, we aggregated the gene expression across all sub-clusters. The aggregated gene count matrix was then normalized by the library size and then log-transformed (log10(TPM / 10 + 1)). Genes were removed if they exhibited low expression (less than 1 in all sub-clusters) or low variance of expression (*i.e.,* the gene expression fold change between the maximum expressed sub-cluster and the median expression across sub-clusters is less than 5). The filtered matrix was used as input for UMAP/0.3.2 visualization ^20^ (metric = “cosine“, min_dist = 0.01, n_neighbors = 30). We then clustered genes based on their 2D UMAP coordinates through densityClust package (rho = 1, delta = 1) ^100^.

### *EasySci-ATAC* library preparation and sequencing

Mouse brain samples were snap-frozen in liquid nitrogen and stored at -80°C. For nuclei extraction, thawed brain samples were minced in PBS using a blade, re-frozen, stored at -80°C, and processed in multiple batches. The detailed step-by-step protocol is included as a supplementary file (**Supplementary file 4**).

### Data processing for *EasySci-ATAC*

Base calls were converted to fastq format and demultiplexed using Illumina’s bcl2fastq/v2.19.0.316 tolerating one mismatched base in barcodes (edit distance (ED) < 2). Downstream sequence processing was similar to sci-ATAC-seq ^101^. Indexed Tn5 barcodes and ligation barcodes were extracted, corrected to its nearest barcode (edit distance (ED) < 2) and reads with uncorrected barcodes (ED >= 2) were removed. Tn5 adaptors were removed from 5’-end and clipped from 3’-end using trim_galore/0.4.1 ^91^. Trimmed reads were mapped to the mouse genome (mm39) using STAR/v2.5.2b ^92^ with default settings. Aligned reads were filtered using samtools/v1.4.1 ^102^ to retain reads mapped in proper pairs with quality score MAPQ > 30 and to keep only the primary aligment. Duplicates were removed by picard MarkDuplicates/v2.25.2 ^103^ per PCR sample. Deduplicated bam files were converted to bedpe format using bedtools/v2.30.0 ^104^, which were further converted to offset-adjusted (+4 bp for plus strand and -5 bp for minus) fragment files (.bed). Deduplicated reads were further split into constituent cellular indices by further demultiplexing reads using the Tn5 and ligation indexes. For each cell, we also created sparse matrices counting reads falling into promoter regions (±1 kb around TSS) for downstream analysis.

To compare the performance of EasySci-ATAC with other sc-ATAC-seq protocols, we utilized a 10x ATAC-v2 dataset (https://www.10xgenomics.com/resources/datasets/8k-adult-mouse-cortex-cells-atac-v2-chromium-x-2-standard) and a conventional sci-ATAC-seq dataset ^16^, all profiling adult mouse brains. We first extracted reads containing barcodes from cells passing quality control (3,636 cells from one PCR well of EasySci-ATAC library, 8067 nuclei from 10×-ATACv2 library; 5494 nuclei from sci-ATAC-seq library). To ensure a balanced comparison, we normalized for sequencing depth differences by subsampling reads from the 10×-ATACv2 and sci-ATAC-seq library, resulting on average 6,360 raw reads per cell across all three libraries. We processed the data through the same computation pipeline described above. Peak calling was performed on each dataset separately with these parameters: --nomodel --extsize 200 --shift - 100 -q 0.05. For peak counting, a union peak set was generated by merging the peaks called from three datasets. Cells were determined to be accessible at a given peak if a read from a cell overlapped with the peak. The peak count matrix was generated by a custom python script with the HTseq package^105^.

### Cell filtering, clustering and annotation for EasySci-ATAC

We used SnapATAC2/v1.99.99.3^106^; ^107^ to perform preprocessing steps for the *EasySci-ATAC* dataset. Cells with less than 1500 fragments and less than 2 TSS Enrichment were discarded. Potential doublet cells and doublet-derived subclusters were detected using an iterative clustering strategy ^10^ modified to suit for scATAC-seq data. Briefly, cells were splitted by individual animals to overcome the large memory use when simulating doublets for the full dataset, and doublet scores were calculated using snap.pp.scrublet() ^94^. Then, all cells were combined, followed by clustering and sub-clustering analysis with spectral embedding and graph-based clustering implemented in SnapATAC2. Cells labeled as doublets (defined by a doublet score cutoff of 0.2) or from doublet-derived sub-clusters (defined by a doublet ratio cutoff of 0.4) were filtered out. In addition, cells with high fragment numbers in each main cluster (defined as cells with fragments number higher than the 95th quantile within the main cluster) were also filtered out. We then generated a gene activity matrix using snap.pp.make_gene_matrix() for the following integration analysis.

We used a deep-learning-based framework scJoint ^24^ to annotate main ATAC-seq cell types using the *EasySci-RNA* dataset as a reference. First, we subsampled 5,000 cells from each main cell type of the *EasySci-RNA* dataset, and selected genes detected in more than 10 cells. Then, the gene count matrix and cell type labels of *EasySci-RNA*, along with the gene activity matrix of *EasySci-ATAC* were input into the scJoint pipeline with default parameters. Jointed embedding layers calculated from scJoint were used for UMAP visualizations using python package umap/v0.5.3 ^108^. Louvain clusters were identified using the Seurat function FindNeighbors() and FindClusters() based on the UMAP coordinates. Cells were assigned to the prediction label with the highest abundance within each louvain cluster. Clusters with low purities (*i.e.,* less than 80% cells were from the highest abundant cell type) were removed upon inspections. Finally, to validate the integration-based annotations, we selected differentially expressed genes identified from the RNA-seq data with the following criteria: fold change between the maximum and the second maximum expressed cell type > 1.5, q-value < 0.05, TPM (transcripts per million) > 20 in the maximum RNA group and RPM (reads per million) > 50 in the maximum ATAC group. Top 10 genes ranked by fold change between the maximum and the second maximum expressed group were selected using RNA-seq data for each cell type. If there were less than 10 genes passing the cutoff, we selected the top genes ranked by the fold change between the maximum expressed cell type and the mean expression of other cell types. We then calculated the aggregated gene count and gene body accessibility (gene activity) for each cell type.

Subcluster level integrations were similar to the main cluster level integrations with mild modifications. For the majority of main cell typescells, we used all cells from the *EasySci-RNA* dataset as input for the integrations. For cortical projection neurons 1, interbrain and midbrain neurons 2, and oligodendrocytes, we subsampled 2,000 cells from each subcluster from the *EasySci-RNA* data for integration analysis. Subcluster id for each ATAC-seq cell was predicted based on the transfer learning implemented in the scJoint pipeline^24^. Similarly, we validated the subcluster level integrations by inspecting the aggregated gene activity of subcluster-specific gene markers in the predicted ATAC subclusters. Subcluster marker genes were identified by differential expression analysis using scRNA-seq data and selected by the following criteria: fold change between the maximum expressed sub-cluster and the mean of all the other subclusters within the same main cell type > 2, FDR < 0.05, TPM (transcripts per million) > 50 in the maximum expressed RNA group and RPM (reads per million) > 50 in the maximum accessible ATAC group. Predicted sub-clusters in the ATAC-seq data passing the enrichment cutoff defined by the log2FC of gene activity > 0.25 between the target subcluster to the rest of cells and containing more than 10 cells were considered matched.

### Peak calling, peak-based dimension reduction and identifications of differential accessible peaks

To define peaks of accessibility, we used MACS2/v2.1.1 ^109^. Nonduplicate ATAC-seq reads of cells from each main cell type were aggregated and peaks were called on each group separately with these parameters: --nomodel --extsize 200 --shift -100 -q 0.05. To correct for differences in read depth or the number of nuclei per cell type, we converted MACS2 peak scores (−log10(q-value)) to ‘score per million’ ^110^ and filtered peaks by choosing a score-per-million cut-off of 1.3. Peak summits were extended by 250bp on either side and then merged with bedtools/v2.30.0. Cells were determined to be accessible at a given peak if a read from a cell overlapped with the peak. The peak count matrix was generated by a custom python script with the HTseq package ^105^.

We used R package Signac/v1.7.0 ^111^ to perform the dimension reduction analysis using the peak-count matrix. We subsampled 5,000 cells from each main cell type and performed TF-IDF normalization using RunTFIDF(), followed by singular value decomposition using RunSVD() and retained the 2nd to 30th dimensions for UMAP visualizations using RunUMAP().

Differentially accessible peaks across cell types were identified using monocle 2 ^98^ with the differentialGeneTest() function. 5,000 cells were subsampled from each cell type for this analysis. Peaks detected in less than 50 cells were filtered out. We selected peaks that were differentially accessible across cell types by the following criteria: 5% FDR (likelihood ratio test), and with TPM > 20 in the target cell type.

### Transcription factor motif analysis

We used ChromVar/v1.16.0 ^59^ to access the TF motif accessibility using a collection of the cisBP motif sets curated by chromVARmotifs/v0.2.0 ^112^; ^59^. To investigate TF regulators at the main cluster level, we subsampled 5,000 cells from each main cell type, and calculated the motif deviation score for each single cell using the Signac wrapper RunChromVAR(). The motif deviation scores of each single cell were rescaled to (0, 10) using R function rescale() and then aggregated for each cell type. In addition, we also aggregated the gene expression of each TF in each cell type. We then computed the Pearson correlations between the aggregated motif matrix and aggregated TF expression matrix after scaling across all main cell types. TF analysis at the subcluster level was performed similarly with modifications. For each cell type of interest, we selected peaks detected in more than 20 cells and only kept cells with more than 500 reads in peaks. Peaks were resized to 500 bp (± 250 bp around the center) and motif occurrences were identified using matchMotifs() function from motifmatchr/v1.16.0 ^113^. The motif deviation matrix was calculated using the ChromVar function computeDeviations(). Then, the motif deviation scores were rescaled to (0, 10) and aggregated per subcluster. Pearson correlation was calculated between the aggregated motif activity and aggregated TF expression across subclusters after scaling. ATAC-seq subclusters with less than 20 cells were excluded from the correlation analysis.

### LDSC analysis

To estimate enrichments of heritability for human traits in differentially accessible peaks, we applied LDSC, which takes summary statistics from a given GWAS as input and quantifies the enrichment of heritability in an annotated set of SNPs conditioned on a baseline model that accounts for the non-random distribution of heritability across the genome. The LDSC computational pipeline was modified from ^16^ and based on the LDSC software (https://github.com/bulik/ldsc). Specifically, to integrate human and mouse data, we first used the UCSC utility liftOver to lift all GWAS SNPs that are used to the mouse genome. We then took the set of differentially accessible peaks (identified from differential accessible analysis using monocle 2 differentialGeneTest() function, defined as q-value < 0.05 and with TPM > 10 in the target cell type) across main clusters and across microglia subclusters, and annotated each SNP according to whether or not it overlapped one of these peaks. We then followed the recommended workflow for running LDSC using HapMap SNPs, precomputed files corresponding to 1000 genomes phase 3, excluding the MHC region to generate an LDSC model for each chromosome and peak set. Only main cell types or subclusters containing DEpeaks in every chromosome are included in the following analysis.

To calculate enrichments based on each model, we first regenerated the baseline model (version 1.1) provided via the LDSC website and used this as the reference for enrichment calculation. We obtained full summary statistics for most GWAS from the Broad LD Hub (https://data.broadinstitute.org/alkesgroup/sumstats_formatted/). Results for all trait/cluster pairs were gathered into a single file. P-values were calculated from z-scores assigned to coefficients reported by ldsc.py and coefficients were divided by the average per-SNP heritability for trait associated with a given test (as calculated from the number of SNPs and overall heritability reported in the .log files from ldsc.py), as recommended by the LDSC authors, producing scaled coefficients. Tests were corrected for multiple hypothesis testing using the Benjamini-Hochberg method and only tests with a q-value of 0.05 or lower were considered significant.

### Spatial gene expression profiling of mouse brains

Spatial gene expression analysis experimental protocol was followed according to Visium Spatial Gene Expression User Guide (catalog no. CG000160), Visium Spatial Tissue Optimization User Guide (catalog no. CG000238 Rev A, 10x Genomics) and Visium Spatial Gene Expression User Guide (catalog no. CG000239 Rev A, 10x Genomics). Briefly, mice were sacrificed, and brains were extracted and frozen with liquid nitrogen. Frozen brain was embedded in OCT (Tissue TEK O.C.T compound) and cryosectioned at -15C (Leica cryostat). Coronally placed brains were cut halfway, to place half coronally sectioned brains at 10um on Visium tissue optimization, or gene expression analysis slides capture areas. User guide CG000160 from 10x Genomics was followed for methanol fixation and H&E stain. After fixation and staining, imaging was performed using Leica DMI8, and images were stitched using Leica Application Suite X and saved into .tiff format. After tissue fixation and staining, Visium Spatial Tissue Optimization User Guide (catalog no. CG000238 Rev A, 10x Genomics) or Visium Spatial Gene Expression User Guide (catalog no. CG000239 Rev A, 10x Genomics) were followed for either protocol optimization, or gene expression analysis, respectively. Tissue optimization was performed according to CG000238, and according to optimization experiments, 18 min permeabilization provided the most optimal signal, and was followed for gene expression library preparation as well. Libraries were prepared according to Visium Spatial Gene Expression User Guide (CG000239, 10x Genomics)

### Analysis for linking cis-regulatory elements (CRE) to putative target genes

We aim to identify links between cis-regulatory elements and putative regulated genes based on their covariance across matched subclusters between RNA and ATAC data. We first constructed pseudo-cells by aggregating the RNA-seq and ATAC-seq profiles of the same subclusters. Aggregated count matrices for RNA-seq and ATAC-seq were normalized to transcripts per million (TPM) and log-transformed after adding one pseudocount. We only retained genes and peaks with TPM value greater than 10 in the maximum expressed pseudo-cells. Then, for each gene, we calculated the Pearson Correlation Coefficient (PCC) between its gene expression and the chromatin accessibility of its nearby accessible sites (minus/plus 500 kb from the TSS) across aggregated subclusters. To define a threshold at PCC score, we also generated a set of background pairs by permuting the subcluster id of the ATAC-seq matrix and with an empirically defined significance threshold of FDR < 0.01, to select significant positively correlated cCRE-gene pairs. We only keep the one top linked gene with the highest PCC for each peak and distal peaks overlapping with the promoters for other genes were filtered out.

### Library preparation and data processing of spatial transcriptomics

Libraries were sequenced using a NextSeq1000 system. BCL files were converted to FASTQ, and raw FASTQ files and .tiff histology images were processed with spaceranger-1.2.2 software. Spaceranger-1.2.2 uses STAR for RNA reads genome alignment, and utilized the GRCm38 (mouse mm10) as the reference genome provided from 10x Genomics. We performed the downstream visualization and clustering analysis of the spatial transcriptomic data following the tutorial of Seurat ^97^ (https://satijalab.org/seurat/articles/spatial_vignette.html) with default parameters.

### Non-negative least squares approach to integrate *EasySci* with external datasets and to locate the spatial distributions of main cell types and subtypes

To annotate the spatial locations of main cell types, we integrated the *EasySci-RNA* data with publicly available 10x Visium spatial transcriptomics dataset (https://satijalab.org/seurat/articles/spatial_vignette.html^28–30^ through a non-negative least squares (NNLS) approach modified from our previous study ^10^. We first aggregated cell-type-specific UMI counts, normalized by the library size, multiplied by 100,000, and log-transformed after adding a pseudo-count. A similar procedure was applied to calculate the normalized gene expression in each spatial spot captured in the 10x Visium dataset. We then applied non-negative least squares (NNLS) regression to predict the gene expression of each spatial spot in 10x Visium data using the gene expression of all cell types recovered in Easy-RNA data:

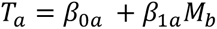

where *T*_*a*_and *M*_*b*_ represent filtered gene expression for target spatial spot from 10x Visium dataset A and all cell types from *EasySci-RNA* dataset B, respectively. To improve accuracy and specificity, we selected cell type-specific genes for each target cell type by: 1) ranking genes based on the expression fold-change between the target cell type vs. the median expression across all cell types, and then selecting the top 200 genes. 2) ranking genes based on the expression fold-change between the target cell type vs. the cell type with maximum expression among all other cell types, and then selecting the top 200 genes. 3) merging the gene lists from step (1) and (2). *β*_1*a*_is the correlation coefficient computed by NNLS regression.

Similarly, we then switch the order of datasets A and B, and predict the gene expression of target cell type (*T*_*b*_) in dataset B with the gene expression of all spatial spots (*M*_*a*_) in dataset A:

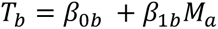

Thus, each spatial spot a in 10x Visium dataset A and each cell type b in *EasySci* dataset B are linked by two correlation coefficients from the above analysis: *β*_*ab*_ for predicting the gene expression in each spatial spot a using b, and *β*_*ba*_for predicting gene expression in each cell type b using a. We combine the two values by:

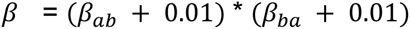

The *β* is then capped to [1, 3]. We find *β* reflects the cell-type-specific abundance across different spatial spots in 10x Visium datasets with high specificity. We thus use *β* as the alpha value (*i.e.,* the opacity of a geom) to plot the spatial distribution of different cell types.

To characterize the expression of sub-cluster specific gene markers, we first normalized the gene expression in each spatial spot of 10x Visium data by the library size, multiplied by 100,000, and log-transformed after adding a pseudo-count. The expression of genes from sub-cluster specific gene markers was aggregated, scaled to z-score and capped to [3, 6]. Of note, the sub-cluster specific gene markers were selected by differentiation expression analysis described above and only DE genes (FDR of 5%, with a >2-fold expression difference between first and second ranked sub-clusters, expression TPM > 50 in at least one sub-cluster) were selected as gene markers. In addition, we examined the aggregated expression of the selected gene markers across all 359 sub-clusters to further validate the specificity of gene markers for labeling target sub-clusters.

We used our NNLS approach to integrate our *EasySci-RNA* dataset with a large single-cell dataset featuring highly detailed cell type annotations^2^. After calculating the combined *β* scores as described above, we established a cutoff value to remove the abundant low matching scores (cutoff: *β* > 0.15) and we established the paired cell types between the two datasets using the highest combined *β* scores.

### Clustering, annotation and differential analysis for human brain samples

A digital gene expression matrix was constructed from the raw sequencing data as described before. To identify distinct clusters of cells corresponding to different cell types in the human brain samples, we co-embedded the human cells from both regions with our mouse brain dataset (up to 5,000 cells randomly sampled from each of 31 cell types), and clusters were annotated based on overlapped cell types. The annotations were manually verified and refined based on marker genes. Following on, the hippocampus and SMTG human dataset were integrated together to construct the same low-dimensional space with only human cells.

Differentially expressed genes between AD and control samples for each cell type in each region were identified using Monocle 2 (Qiu et al., 2017) with the differentialGeneTest() function. Main cell types with less than 50 cells were excluded from the analysis (i.e, choroid plexus epithelial cells and vascular leptomeningeal cells in the SMTG). DE genes were filtered based on the following cutoffs: q-value < 0.05, with FC > 1.5 between the maximum and second expressed condition, and with transcripts per million (TPM) > 50 in the highest expressed condition. To further validate human-mouse shared gene expression changes, we used a recently published Alzheimer’s disease single-cell dataset from the human prefrontal cortex ^6^.

## Data Availability

The data generated by this study can be downloaded in raw and processed forms from the NCBI Gene Expression Omnibus (GSE212606, reviewers’ token: sxitgiusjjwjvuh).

## Code Availability

The detailed experimental protocol and computation scripts of *EasySci* were included as supplementary files. Furthermore, the EasySci-RNA computational processing pipeline is available on GitHub at this repository: https://github.com/JunyueCaoLab/EasySci.

## Supplementary Tables (provided as Microsoft Excel files)

**Supplementary Table 1:** Differentially expressed genes across main cell types. For each gene, the “Cell type” is the cell type with the highest expression, with the expression level quantified by transcripts per million in “TPM in cell type”. The “Q-value” is the false detection rate (one-sided likelihood ratio test with adjustment for multiple comparisons) for the differential expression test across different cell clusters. The “Fold change” is the fold change between the max expressed cell type and the second expressed cell type.

**Supplementary Table 2:** Differentially expressed isoforms across main cell types. For each isoform (“Isoform”), the “Cell type” is the cell type with the highest expression. The “P-value” is the raw p-value for the differential expression test across different cell types; and the “Q-value” is the false detection rate (one-sided likelihood ratio test with adjustment for multiple comparisons). The “Effect size” is the effect size between the max expressed cell type and the second expressed cell type.

**Supplementary Table 3:** Differentially accessible sites for main cell types. For each peak (“Peak”), the “Max cell type” is the cell type with the highest accessibility (“Peak accessibility in max cell type”). The “Second cell type” is the cell type with the second highest accessibility (“Peak accessibility in second cell type”). The “Fold change” is the fold change between the max accessibility and the second max accessibility. The “P-value” is the raw p-value for the differential accessibility test across different cell types, and the “Q-value” is the false detection rate (one-sided likelihood ratio test with adjustment for multiple comparisons).

**Supplementary Table 4:** Differentially expressed exons across sub-clusters within each main cell type. For each sub-cluster (“Cell sub-cluster ID”), the following features of marker genes are listed: gene symbol (“Gene name”); Ensembl ID of the gene and the exon (“Exon ID”); false detection rate (one-sided likelihood ratio test with adjustment for multiple comparisons) for the differential expression test across different cell sub-clusters within each main cell type (“Q-value”); fold change of the marker exon expression between the max and second expressed cell sub-cluster (“Fold change”); expression level of the marker exon quantified by transcripts per million in max sub-cluster (“TPM in max sub-cluster”). Marker exons are defined by Q-value < 0.05, Fold change > 2 and TPM in max subcluster > 50.

**Supplementary Table 5:** Validation of EasySci-RNA subclusters using integration analysis with EasySci-ATAC, Zeisel et al^2^ and Ortiz el al^36^. For the EasySci-ATAC integration, we show if the subcluster was validated by this analysis. For the Zeisel at al integration, we indicate the top matching cell type, if we could validate the subcluster. For the Ortiz et al integration, we show the consensus anatomical region from the top 10 pixels with the highest mapping score above 1, if we could validate the subcluster.

**Supplementary Table 6:** Gene module analysis results. For each gene module (“Gene module ID”), the following information about the genes belonging to that gene module is listed: Ensembl ID (“Gene ID”); type of gene (“Gene type”); gene symbol (“Gene name”); UMAP visualization coordinates of the genes based on their expression variance across all 359 cell sub-clusters (“UMAP 1”, “UMAP 2”).

**Supplementary Table 7:** Differentially abundant sub-clusters between adult and young samples. Sub-clusters (“Cell sub-clusters”) abundances were compared between the 3 vs 6 months old groups (“Condition”), and the following statistical values are listed: false detection rate (likelihood ratio test with adjustment for multiple comparisons) for the differential abundance test across age groups (“Q-value”); log2 fold change of the cell sub-cluster abundance between the age groups (“Log2(Fold change)”); the number of cells compared in the sub-cluster (“Number of cells”); whether the sub-cluster is upregulated, downregulated or there is no significant change (“Final change”, significance was determined by Fold change > 2, Q-value < 0.05 and more than 20 cells in both male and female samples).

**Supplementary Table 8:** Differentially abundant sub-clusters between aged and adult samples. Sub-clusters (“Cell sub-clusters”) abundances were compared between the 6 vs 21 months old groups (“Condition”), and the following statistical values are listed: false detection rate (likelihood ratio test with adjustment for multiple comparisons) for the differential abundance test across age groups (“Q-value”); log2 fold change of the cell sub-cluster abundance between the age groups (“Log2(Fold change)”); the number of cells compared in the sub-cluster (“Number of cells”); whether the sub-cluster is upregulated, downregulated or there is no significant change (“Final change”, significance was determined by Fold change > 2, Q-value < 0.05 and more than 20 cells in both male and female samples).

**Supplementary Table 9:** Differentially expressed genes between aged and adult for all sub-clusters. For each subcluster in the dataset (“Cell subcluster”), the following information is listed: gene symbol (“Gene name”); gene Ensembl ID (“Gene ID”); the false detection rate (one-sided likelihood ratio test with adjustment for multiple comparisons) for the differential expression test across the age groups (“Q-value”); fold change between the max expressed age group and second expressed age group (“Fold change ”); expression level quantified by transcripts per million in the max age group (“TPM in max condition”); the age group where the max expression was detected (“Max condition”). Only significant genes are listed in the table, which is defined by Q-value < 0.05, Fold change > 2 and TPM in max condition > 50.

**Supplementary Table 10:** Differentially abundant sub-clusters between wild-type and EOAD model. Sub-clusters (“Cell sub-clusters”) abundances were compared between the wild-type and EOAD model (5xFAD) groups (“Condition”), and the following statistical values are listed: false detection rate (likelihood ratio test with adjustment for multiple comparisons) for the differential abundance test between conditions (“Q-value”); log2 fold change of the cell sub-cluster abundance between conditions (“Log2(Fold change)”); the number of cells compared in the sub-cluster (“Number of cells”); whether the sub-cluster is upregulated, downregulated or there is no significant change (“Final change”, significance was determined by Fold change > 2, Q-value < 0.05 and more than 20 cells in both male and female samples).

**Supplementary Table 11:** Differentially abundant sub-clusters between wild-type and LOAD model. Sub-clusters (“Cell sub-clusters”) abundances were compared between the wild-type and LOAD model (APOE*4/Trem2*R47H) groups (“Condition”), and the following statistical values are listed: false detection rate (likelihood ratio test with adjustment for multiple comparisons) for the differential abundance test between conditions (“Q-value”); log2 fold change of the cell sub-cluster abundance between conditions (“Log2(Fold change)”); the number of cells compared in the sub-cluster (“Number of cells”); whether the sub-cluster is upregulated, downregulated or there is no significant change (“Final change”, significance was determined by Fold change > 2, Q-value < 0.05 and more than 20 cells in both male and female samples).

**Supplementary Table 12:** Differentially expressed genes between wild-type and EOAD model (5xFAD) for all sub-clusters. For each subcluster in the dataset (“Cell subcluster”), the following information is listed: gene symbol (“Gene name”); gene Ensembl ID (“Gene ID”); the false detection rate (one-sided likelihood ratio test with adjustment for multiple comparisons) for the differential expression test across conditions (“Q-value”); fold change between the max expressed group and second expressed group (“Fold change”); expression level quantified by transcripts per million in the max group (“TPM in max condition”); the condition where the max expression was detected (“Max condition”). Only significant genes are listed in the table, which is defined by Q-value < 0.05, Fold change > 2 and TPM in max condition > 50.

**Supplementary Table 13:** Differentially expressed genes between wild-type and LOAD model (APOE*4/Trem2*R47H) for all sub-clusters. For each subcluster in the dataset (“Cell subcluster”), the following information is listed: gene symbol (“Gene name”); gene Ensembl ID (“Gene ID”); the false detection rate (one-sided likelihood ratio test with adjustment for multiple comparisons) for the differential expression test across conditions (“Q-value”); fold change between the max expressed group and second expressed group (“Fold change”); expression level quantified by transcripts per million in the max group (“TPM in max condition”); the condition where the max expression was detected (“Max condition”). Only significant genes are listed in the table, which is defined by Q-value < 0.05, Fold change > 2 and TPM in max condition > 50.

**Supplementary Table 14:** Metadata of human brain samples included in this study.

**Supplementary Table 15:** Differentially expressed genes between control and AD human brain samples for each main cell type in each region. For each main cell type (“Main cluster name”) in each region (“Region”), the following information is listed: gene symbol (“Gene name”); the max and the second expressed group (“Max condition”, “Second condition”) along with expression level quantified by transcripts per million (“TPM in max condition”, “TPM in second condition”) and the fold change (“Fold change”); the false detection rate (one-sided likelihood ratio test with adjustment for multiple comparisons) for the differential expression test across the two conditions (“Q-value”). Only significant genes are listed in the table, which is defined by Q-value < 0.05, Fold change > 1.5 and TPM in max condition > 50.

## Supplementary files

**Supplementary file 1:** Detailed experiment protocols for *EasySci-RNA*, including all materials and equipment needed, step-by-step descriptions, and representative gel images.

**Supplementary file 2:** Primer sequences used in the *EasySci-RNA* experiment, including multiple plates of short dT RT primers, random hexamer RT primers, ligation primers and P7 PCR primers. The columns indicate the positions on the 96-well plate (Well position), an identifier of the sequence (Name), the full primer sequence (Sequence) and the barcode sequence (Barcode).

**Supplementary file 3:** Computational pipeline scripts and notes for processing *EasySci-RNA* data, from sequencer-generated files to single-cell gene count matrix.

**Supplementary file 4:** Detailed experiment protocols for *EasySci-ATAC*, including all materials and equipment needed, step-by-step descriptions, and representative gel images.

**Supplementary file 5:** Primer sequences used in the *EasySci-ATAC* experiment, including N5/N7 oligos used in indexed Tn5 assembly, ligation primers and P7 PCR primers. The columns indicate the positions on the 96-well plate (Well position), an identifier of the sequence (Name), the full primer sequence (Sequence) and the barcode sequence (Barcode).

**Supplementary file 6:** Computational pipeline scripts and notes for processing *EasySci-ATAC* data, from sequencer-generated files to single-cell read files.

## Notes

### Competing Interest Statement

The authors have declared no competing interest.

### Summary of Updates

The revised version of the manuscript contains further benchmarking of the novel EasySci method, further validation and annotation of detected cell types and novel biological observations. Furthermore, we corrected a typo in a catalog ID in Supplementary file 1.

## Reference

1. Erö, C., Gewaltig, M.-O., Keller, D. & Markram, H. A Cell Atlas for the Mouse Brain. Front. Neuroinform. 12, 84 (2018).

2. Zeisel, A., et al. Molecular Architecture of the Mouse Nervous System. Cell 174, 999–1014.e22 (2018).

3. Mathys, H., et al. Single-cell transcriptomic analysis of Alzheimer’s disease. Nature 570, 332–337 (2019).

4. Xia, X., Jiang, Q., McDermott, J. & Han, J.-D. J. Aging and Alzheimer’s disease: Comparison and associations from molecular to system level. Aging Cell 17, e12802 (2018).

5. Ximerakis, M., et al. Single-cell transcriptomic profiling of the aging mouse brain. Nat. Neurosci. 22, 1696–1708 (2019).

6. Morabito, S., et al. Single-nucleus chromatin accessibility and transcriptomic characterization of Alzheimer’s disease. Nature Genetics vol. 53 1143–1155 Preprint at https://doi.org/10.1038/s41588-021-00894-z (2021).

7. Tabula Muris Consortium. A single-cell transcriptomic atlas characterizes ageing tissues in the mouse. Nature 583, 590–595 (2020).

8. Wang, R., et al. Construction of a cross-species cell landscape at single-cell level. Nucleic Acids Res. (2022) doi:10.1093/nar/gkac633.

9. Cao, J., et al. Comprehensive single-cell transcriptional profiling of a multicellular organism. Science 357, 661–667 (2017).

10. Cao, J., et al. A human cell atlas of fetal gene expression. Science 370, (2020).

11. Cao, J., et al. The single-cell transcriptional landscape of mammalian organogenesis. Nature 566, 496–502 (2019).

12. Ma, S., et al. Chromatin Potential Identified by Shared Single-Cell Profiling of RNA and Chromatin. Cell (2020) doi:10.1016/j.cell.2020.09.056.

13. Martin, B. K., et al. An optimized protocol for single cell transcriptional profiling by combinatorial indexing. (2021) doi:10.48550/arXiv.2110.15400.

14. Ding, J., et al. Systematic comparison of single-cell and single-nucleus RNA-sequencing methods. Nat. Biotechnol. 38, 737–746 (2020).

15. Domcke, S., et al. A human cell atlas of fetal chromatin accessibility. Science 370, (2020).

16. Cusanovich, D. A., et al. A Single-Cell Atlas of In Vivo Mammalian Chromatin Accessibility. Cell 174, 1309–1324.e18 (2018).

17. Oakley, H., et al. Intraneuronal beta-amyloid aggregates, neurodegeneration, and neuron loss in transgenic mice with five familial Alzheimer’s disease mutations: potential factors in amyloid plaque formation. J. Neurosci. 26, 10129–10140 (2006).

18. Desimone, A., et al. The influence of ApoE4 on the clinical outcomes and pathophysiology of degenerative cervical myelopathy. JCI Insight 6, (2021).

19. Xiang, X., et al. The Trem2 R47H Alzheimer’s risk variant impairs splicing and reduces Trem2 mRNA and protein in mice but not in humans. Mol. Neurodegener. 13, 49 (2018).

20. McInnes, L., Healy, J., Saul, N. & Großberger, L. UMAP: Uniform Manifold Approximation and Projection. Journal of Open Source Software vol. 3 861 Preprint at https://doi.org/10.21105/joss.00861 (2018).

21. Blondel, V. D., Guillaume, J.-L., Lambiotte, R. & Lefebvre, E. Fast unfolding of communities in large networks. Journal of Statistical Mechanics: Theory and Experiment vol. 2008 P10008 Preprint at https://doi.org/10.1088/1742-5468/2008/10/p10008 (2008).

22. Booeshaghi, A. S., et al. Isoform cell-type specificity in the mouse primary motor cortex. Nature 598, 195–199 (2021).

23. Bray, N. L., Pimentel, H., Melsted, P. & Pachter, L. Near-optimal probabilistic RNA-seq quantification. Nat. Biotechnol. 34, 525–527 (2016).

24. Lin, Y., et al. scJoint integrates atlas-scale single-cell RNA-seq and ATAC-seq data with transfer learning. Nat. Biotechnol. 40, 703–710 (2022).

25. Yeh, H. & Ikezu, T. Transcriptional and Epigenetic Regulation of Microglia in Health and Disease. Trends Mol. Med. 25, 96–111 (2019).

26. Watakabe, A., et al. Comparative analysis of layer-specific genes in Mammalian neocortex. Cereb. Cortex 17, 1918–1933 (2007).

27. McEvilly, R. J., et al. Requirement for Brn-3.0 in differentiation and survival of sensory and motor neurons. Nature 384, 574–577 (1996).

28. Genomics, 10×. Mouse Brain Section (Coronal). (2019).

29. Genomics, 10×. Mouse Brain Serial Section 1 (Sagittal-Posterior). (2019).

30. Genomics, 10×. Mouse Brain Serial Section 1 (Sagittal-Anterior). (2019).

31. Rosenberg, A. B., et al. Single-cell profiling of the developing mouse brain and spinal cord with split-pool barcoding. Science 360, 176–182 (2018).

32. Stoeckius, M., et al. Cell Hashing with barcoded antibodies enables multiplexing and doublet detection for single cell genomics. Genome Biol. 19, 224 (2018).

33. Mays, J. C., et al. Single-cell RNA sequencing of the mammalian pineal gland identifies two pinealocyte subtypes and cell type-specific daily patterns of gene expression. PLoS One 13, e0205883 (2018).

34. Campbell, J. N., et al. A molecular census of arcuate hypothalamus and median eminence cell types. Nat. Neurosci. 20, 484–496 (2017).

35. Kleshchevnikov, V., et al. Cell2location maps fine-grained cell types in spatial transcriptomics. Nat. Biotechnol. 1–11 (2022).

36. Ortiz, C., et al. Molecular atlas of the adult mouse brain. Sci Adv 6, eabb3446 (2020).

37. Kuleshov, M. V., et al. Enrichr: a comprehensive gene set enrichment analysis web server 2016 update. Nucleic Acids Res. 44, W90–7 (2016).

38. Liu, J., et al. Tbx19, a tissue-selective regulator of POMC gene expression. Proc. Natl. Acad. Sci. U. S. A. 98, 8674–8679 (2001).

39. Tufo, C., et al. Development of the mammalian main olfactory bulb. Development 149, (2022).

40. Sokolowski, J. D., et al. Brain-specific angiogenesis inhibitor-1 expression in astrocytes and neurons: implications for its dual function as an apoptotic engulfment receptor. Brain Behav. Immun. 25, 915– 921 (2011).

41. Tepe, B., et al. Single-Cell RNA-Seq of Mouse Olfactory Bulb Reveals Cellular Heterogeneity and Activity-Dependent Molecular Census of Adult-Born Neurons. Cell Rep. 25, 2689–2703.e3 (2018).

42. Barraud, P., et al. Neural crest origin of olfactory ensheathing glia. Proc. Natl. Acad. Sci. U. S. A. 107, 21040–21045 (2010).

43. Monavarfeshani, A., Knill, C. N., Sabbagh, U., Su, J. & Fox, M. A. Region- and Cell-Specific Expression of Transmembrane Collagens in Mouse Brain. Front. Integr. Neurosci. 11, 20 (2017).

44. Puverel, S., Nakatani, H., Parras, C. & Soussi-Yanicostas, N. Prokineticin receptor 2 expression identifies migrating neuroblasts and their subventricular zone transient-amplifying progenitors in adult mice. J. Comp. Neurol. 512, 232–242 (2009).

45. Pastrana, E., Cheng, L.-C. & Doetsch, F. Simultaneous prospective purification of adult subventricular zone neural stem cells and their progeny. Proc. Natl. Acad. Sci. U. S. A. 106, 6387– 6392 (2009).

46. Kumar, A., et al. Transcriptomic analysis of the signature of neurogenesis in human hippocampus suggests restricted progenitor cell progression post-childhood. IBRO Rep 9, 224–232 (2020).

47. Marques, S., et al. Transcriptional Convergence of Oligodendrocyte Lineage Progenitors during Development. Dev. Cell 46, 504–517.e7 (2018).

48. Zhang, Y., et al. An RNA-sequencing transcriptome and splicing database of glia, neurons, and vascular cells of the cerebral cortex. J. Neurosci. 34, 11929–11947 (2014).

49. Lu, Z. et al. A comprehensive view of cell-type-specific temporal dynamics in human and mouse brains. bioRxiv 2022.10.01.509820 (2022) doi:10.1101/2022.10.01.509820.

50. Graham, V., Khudyakov, J., Ellis, P. & Pevny, L. SOX2 functions to maintain neural progenitor identity. Neuron 39, 749–765 (2003).

51. Li, J., et al. Transcription Factors Sp8 and Sp9 Coordinately Regulate Olfactory Bulb Interneuron Development. Cereb. Cortex 28, 3278–3294 (2018).

52. Keren-Shaul, H., et al. A Unique Microglia Type Associated with Restricting Development of Alzheimer’s Disease. Cell vol. 169 1276–1290.e17 Preprint at https://doi.org/10.1016/j.cell.2017.05.018 (2017).

53. Zhou, Y., et al. Human and mouse single-nucleus transcriptomics reveal TREM2-dependent and TREM2-independent cellular responses in Alzheimer’s disease. Nat. Med. 26, 131–142 (2020).

54. Kenigsbuch, M., et al. A shared disease-associated oligodendrocyte signature among multiple CNS pathologies. Nat. Neurosci. 25, 876–886 (2022).

55. See, A. P., et al. The role of STAT3 activation in modulating the immune microenvironment of GBM. J. Neurooncol. 110, 359–368 (2012).

56. Paillasse, M. R. & de Medina, P. The NR4A nuclear receptors as potential targets for anti-aging interventions. Med. Hypotheses 84, 135–140 (2015).

57. Di Giorgio, E., et al. HDAC4 degradation during senescence unleashes an epigenetic program driven by AP-1/p300 at selected enhancers and super-enhancers. Genome Biol. 22, 129 (2021).

58. Zhang, Y. & Wang, P. Age-Related Increase of Insulin-Degrading Enzyme Is Inversely Correlated with Cognitive Function in APPswe/PS1dE9 Mice. Med. Sci. Monit. 24, 2446–2455 (2018).

59. Schep, A. N., Wu, B., Buenrostro, J. D. & Greenleaf, W. J. chromVAR: inferring transcription-factor-associated accessibility from single-cell epigenomic data. Nat. Methods 14, 975–978 (2017).

60. Hashimoto, Y., et al. A rescue factor abolishing neuronal cell death by a wide spectrum of familial Alzheimer’s disease genes and Abeta. Proc. Natl. Acad. Sci. U. S. A. 98, 6336–6341 (2001).

61. Cavalcante, G. C., et al. Mitochondrial Genetics Reinforces Multiple Layers of Interaction in Alzheimer’s Disease. Biomedicines 10, (2022).

62. Mielke, M. M. & Lyketsos, C. G. Alterations of the sphingolipid pathway in Alzheimer’s disease: new biomarkers and treatment targets? Neuromolecular Med. 12, 331–340 (2010).

63. Tong, Y., Xu, Y., Scearce-Levie, K., Ptácek, L. J. & Fu, Y.-H. COL25A1 triggers and promotes Alzheimer’s disease-like pathology in vivo. Neurogenetics 11, 41–52 (2010).

64. Butler, T., et al. Volume of the human septal forebrain region is a predictor of source memory accuracy. J. Int. Neuropsychol. Soc. 18, 157–161 (2012).

65. Oeckinghaus, A. & Ghosh, S. The NF-kappaB family of transcription factors and its regulation. Cold Spring Harb. Perspect. Biol. 1, a000034 (2009).

66. Liu, X.-F., et al. Nrf2 as a target for prevention of age-related and diabetic cataracts by against oxidative stress. Aging Cell 16, 934–942 (2017).

67. Bommer, G. T. & MacDougald, O. A. Regulation of lipid homeostasis by the bifunctional SREBF2-miR33a locus. Cell Metab. 13, 241–247 (2011).

68. Finucane, H. K., et al. Partitioning heritability by functional annotation using genome-wide association summary statistics. Nat. Genet. 47, 1228–1235 (2015).

69. Seripa, D., et al. The RELN locus in Alzheimer’s disease. J. Alzheimers. Dis. 14, 335–344 (2008).

70. Herring, A., et al. Reelin depletion is an early phenomenon of Alzheimer’s pathology. J. Alzheimers. Dis. 30, 963–979 (2012).

71. Attwood, M. M. & Schiöth, H. B. Characterization of Five Transmembrane Proteins: With Focus on the Tweety, Sideroflexin, and YIP1 Domain Families. Front Cell Dev Biol 9, 708754 (2021).

72. Buckley, M. T., et al. Cell-type-specific aging clocks to quantify aging and rejuvenation in neurogenic regions of the brain. Nat Aging 3, 121–137 (2023).

73. de Faria, O., Jr et al. TMEM10 Promotes Oligodendrocyte Differentiation and is Expressed by Oligodendrocytes in Human Remyelinating Multiple Sclerosis Plaques. Sci. Rep. 9, 3606 (2019).

74. Volkert, M. R. & Crowley, D. J. Preventing Neurodegeneration by Controlling Oxidative Stress: The Role of OXR1. Front. Neurosci. 14, 611904 (2020).

75. Schmidt, L. S. & Linehan, W. M. FLCN: The causative gene for Birt-Hogg-Dubé syndrome. Gene 640, 28–42 (2018).

76. Cooper, W. N., et al. RASSF2 associates with and stabilizes the proapoptotic kinase MST2. Oncogene 28, 2988–2998 (2009).

77. Luo, J., et al. PTPRG activates m6A methyltransferase VIRMA to block mitochondrial autophagy mediated neuronal death in Alzheimer’s disease. bioRxiv (2022) doi:10.1101/2022.03.11.22272061.

78. Silva, I., Silva, J., Ferreira, R. & Trigo, D. Glymphatic system, AQP4, and their implications in Alzheimer’s disease. Neurol Res Pract 3, 5 (2021).

79. Kulijewicz-Nawrot, M., Syková, E., Chvátal, A., Verkhratsky, A. & Rodríguez, J. J. Astrocytes and glutamate homoeostasis in Alzheimer’s disease: a decrease in glutamine synthetase, but not in glutamate transporter-1, in the prefrontal cortex. ASN Neuro 5, 273–282 (2013).

80. Hüttenrauch, M., et al. Glycoprotein NMB: a novel Alzheimer’s disease associated marker expressed in a subset of activated microglia. Acta Neuropathol Commun 6, 108 (2018).

81. Zipfel, P., Rochais, C., Baranger, K., Rivera, S. & Dallemagne, P. Matrix Metalloproteinases as New Targets in Alzheimer’s Disease: Opportunities and Challenges. J. Med. Chem. 63, 10705–10725 (2020).

82. Baranger, K., et al. MT5-MMP is a new pro-amyloidogenic proteinase that promotes amyloid pathology and cognitive decline in a transgenic mouse model of Alzheimer’s disease. Cell. Mol. Life Sci. 73, 217–236 (2016).

83. Arawaka, S., Machiya, Y. & Kato, T. Heat shock proteins as suppressors of accumulation of toxic prefibrillar intermediates and misfolded proteins in neurodegenerative diseases. Curr. Pharm. Biotechnol. 11, 158–166 (2010).

84. Cornejo, V. H. & Hetz, C. The unfolded protein response in Alzheimer’s disease. Semin. Immunopathol. 35, 277–292 (2013).

85. Neff, R. A., et al. Molecular subtyping of Alzheimer’s disease using RNA sequencing data reveals novel mechanisms and targets. Sci Adv 7, (2021).

86. Niccolini, F., et al. Altered PDE10A expression detectable early before symptomatic onset in Huntington’s disease. Brain 138, 3016–3029 (2015).

87. Niccolini, F., et al. Loss of phosphodiesterase 10A expression is associated with progression and severity in Parkinson’s disease. Brain 138, 3003–3015 (2015).

88. Karch, C. M. & Goate, A. M. Alzheimer’s disease risk genes and mechanisms of disease pathogenesis. Biol. Psychiatry 77, 43–51 (2015).

89. Kotredes, K. P., et al. Uncovering Disease Mechanisms in a Novel Mouse Model Expressing Humanized APOEε4 and Trem2*R47H. Front. Aging Neurosci. 13, 735524 (2021).

90. Website. https://www.10xgenomics.com/resources/datasets/5k-adult-mouse-brain-nuclei-isolated-with-chromium-nuclei-isolation-kit-3-1-standard.

91. Krueger, F., James, F., Ewels, P., Afyounian, E. & Schuster-Boeckler, B. GitHub - FelixKrueger/TrimGalore: A wrapper around Cutadapt and FastQC to consistently apply adapter and quality trimming to FastQ files, with extra functionality for RRBS data. (2021) doi:10.5281/zenodo.5127899.

92. Dobin, A., et al. STAR: ultrafast universal RNA-seq aligner. Bioinformatics 29, 15–21 (2013).

93. Zheng, G. X. Y., et al. Massively parallel digital transcriptional profiling of single cells. Nat. Commun. 8, 14049 (2017).

94. Wolock, S. L., Lopez, R. & Klein, A. M. Scrublet: Computational Identification of Cell Doublets in Single-Cell Transcriptomic Data. Cell Syst 8, 281–291.e9 (2019).

95. Yao, Z., et al. A transcriptomic and epigenomic cell atlas of the mouse primary motor cortex. Nature 598, 103–110 (2021).

96. Kozareva, V., et al. A transcriptomic atlas of mouse cerebellar cortex comprehensively defines cell types. Nature 598, 214–219 (2021).

97. Stuart, T., et al. Comprehensive Integration of Single-Cell Data. Cell 177, 1888–1902.e21 (2019).

98. Qiu, X., et al. Reversed graph embedding resolves complex single-cell trajectories. Nat. Methods 14, 979–982 (2017).

99. Melsted, P., et al. Modular, efficient and constant-memory single-cell RNA-seq preprocessing. Nat. Biotechnol. 1–6 (2021).

100. Rodriguez, A. & Laio, A. Machine learning. Clustering by fast search and find of density peaks. Science 344, 1492–1496 (2014).

101. Cao, J., et al. Joint profiling of chromatin accessibility and gene expression in thousands of single cells. Science 361, 1380–1385 (2018).

102. Li, H., et al. The Sequence Alignment/Map format and SAMtools. Bioinformatics 25, 2078–2079 (2009).

103. Broad Institute. Picard toolkit. Broad Institute, GitHub repository (2019).

104. Quinlan, A. R. & Hall, I. M. BEDTools: a flexible suite of utilities for comparing genomic features. Bioinformatics 26, 841–842 (2010).

105. Anders, S., Pyl, P. T. & Huber, W. HTSeq--a Python framework to work with high-throughput sequencing data. Bioinformatics 31, 166–169 (2015).

106. Fang, R., et al. Comprehensive analysis of single cell ATAC-seq data with SnapATAC. Nat. Commun. 12, 1337 (2021).

107. Zhang, K. SnapATAC2: Single Nucleus Analysis Pipeline for ATAC-seq — SnapATAC2 2.0.0 documentation, https://kzhang.org/SnapATAC2/index.html. (2022).

108. McInnes, L. UMAP: Uniform Manifold Approximation and Projection for Dimension Reduction — umap 0.5 documentation, https://umap-learn.readthedocs.io/en/latest/. (2018).

109. Zhang, Y., et al. Model-based analysis of ChIP-Seq (MACS). Genome Biol. 9, R137 (2008).

110. Corces, M. R. et al. The chromatin accessibility landscape of primary human cancers. Science 362, (2018).

111. Stuart, T., Srivastava, A., Madad, S., Lareau, C. A. & Satija, R. Single-cell chromatin state analysis with Signac. Nat. Methods 18, 1333–1341 (2021).

112. Weirauch, M. T., et al. Determination and inference of eukaryotic transcription factor sequence specificity. Cell 158, 1431–1443 (2014).

113. Schep, A. motifmatchr: Fast Motif Matching in R, https://github.com/GreenleafLab/motifmatchr/. (2017).

