## Supplementary file 4 for "A global view of aging and Alzheimer’s pathogenesis-associated cell population dynamics and molecular signatures in the human and mouse brains"

**EasySci-ATAC protocol**

**Abstract**

Single-cell combinatorial indexing ('sci-') is a methodological framework that employs split-pool barcoding to uniquely label the nucleic acid contents of large numbers of single cells or nuclei. Here, we present a new 3-level sci-ATAC-seq method (EasySci-ATAC), optimized from the recently published single-cell chromatin accessibility profiling method by combinatorial indexing (sci-ATAC-seq3). The new EasySci-ATAC method allows for the preparation of 1,000,000 nuclei for under $1,000.

**Protocol workflow**

- Buffer Preparation (Steps 1-6)
- Ligation Primer Loading (Step 7-10)
- Tn5 loading (Step 11)
- Nuclei Extraction (Step 12-18)
- Nuclei Wash (Steps 19-21)
- Nuclei Counting (Step 22-23)
- Tagmentation (Steps 24-28)
- Pool/Centrifuge/Resuspend/Redistribute (Steps 29-34)
- Ligation (Steps 35-40)
- Pool/Centrifuge/Resuspend/Redistribute/Quantify (Steps 41-46)
- Proteinase K treatment (Step 47-50)
- PCR (Step 51-55)
- Sequencing Library Purification (Steps 56-61)

To validate the experimental setup, it is recommended to start with a species-mixing experiment. We normally start with a mixture of human (HEK 293T) and mouse (NIH/3T3) cells. A good run normally yields single-cell chromatin accessibility with 3000~4000 unique fragments (with over 10,000 sequencing reads) per cell and >90% purity.

**Primer Sequences used**

All primer sequences including Tn5 oligos/Ligation/PCR primers are attached as a separate excel file (EasySci-ATAC_primer_sequences.xlsx). All primers are ordered from IDT. Most are with standard desalting, except Tn5 oligos and universal P5 primers which we use HPLC purifications.

**Required Equipments**

- Hemocytometers (Neubauer Improved, Bulldog Bio VWR #102966-632)
- Centrifuge (Eppendorf 5702 RH)
- Eppendorf Mastercycler (4x)
- Freezer (-20C, -80C) and Refrigerator (4C)
- Gel Imager
- Ice Buckets
- Microscope
- Multi-channel Pipettes (2-20μL, 20-200μL) (Rainin Instruments)
- Pipettors
- Liquid nitrogen tank for sample storage
- FreezeCell Cell Freezing Container (GeneSeeSci, catalog number: 27-802) Eppendorf ThermoMixer C (5382000023) OR Fisherbrand Nutating Mixer (88861043) ]

**Materials used**

- 1 M Tris-HCl, pH 7.5 (VWR, 97062-936)
- 5 M NaCl (VWR, 97062-858)
- 1 M MgCl2 (VWR, 97062-848)
- Tween-20 (Sigma, P9416)
- cOmplete™, EDTA-free Protease Inhibitor Cocktail (Sigma, 11873580001)
- IGEPAL CA-630 (VWR, IC0219859650)
- Dimethylformamide (Fisher, AC327175000)
- 0.5 M EDTA (VWR, 97062-656)
- Sperimidine (Sigma, S3256-1g)
- DMSO (VWR, 97063-136)
- T4 DNA Ligase (NEB, M0202L)
- EB buffer (Qiagen, 19086)
- TE Buffer (IDTE, 11-05-01-05)
- 10% SDS (VWR, E719-100ML)
- Proteinase K (Sigma, 3115828001)
- Tween-20 (Sigma, P9416)
- NEBNext® High-Fidelity 2X PCR Master MixNEB (NEB, M0541L)
- DNA Clean & Concentrator kit (Zymoresearch, D4014)
- Zymoclean Gel DNA Recovery Kit (Zymoresearch, D4007)
- 6-cm Cell culture dishes(Genesee, 25-260)
- Razor Blades (VWR, 100491-872)
- 1.5 mL Eppendorf™ DNA LoBind Tubes (Eppendorf, 22431021)
- 40 µm cell strainers (VWR, 470236-276)
- 5 mL Syringes (Fisher, 309603)
- pluriStrainer Mini 5 µm filter (Pluriselect, 43-10005-70)
- pluriStrainer Mini 40 µm filter (Pluriselect, 43-10040-70)
- 96-well plates (Genesee, 24-302)
- 2% E-Gel™ EX Agarose Gels (Invitrogen, G402022)
- E-Gel™ 50 bp DNA Ladder (Invitrogen, 10488099)
- Tn5 MEr oligos: /5Phos/CTGTCTCTTATACACATCT
- 12 indexed N5 oligos: (100 μM, /5Phos/ACGACGCTCTTCCGATCT[6bp-barcode]AGATGTGTATAAGAGACAG; IDT, HPLC purification)
- 32 indexed N7 oligos: (100 μM, CGTGTGCTCTTCCGATCT[6bp-barcode]AGATGTGTATAAGAGACAG; IDT, HPLC purification)
- Ligation Adapter Primer (100 μM, AGATCGGAAGAGCGTCGTGTAGGGAAAGAGTGT; IDT)
- 384 indexed ligation primers (100 μM, 5’-AATGATACGGCGACCACCGAGATCTACAC[10bp barcode]ACACTCTTTCCCTAC-3’; IDT)
- Universal P5 primer (10 μM, 5′-AATGATACGGCGACCACCGAGATCTACAC-3′; IDT, HPLC purification)
- 96 indexed P7 primers (10 μM, 5′-CAAGCAGAAGACGGCATACGAGAT[10bp-i7-barcode]GTGACTGGAGTTCAGACGTGTGCTCTTCCGATCT-3′; IDT)

**Buffer Preparation**

1. **500 mL Nuclei Buffer (NB, Stored in 4 °C)**

10 mM Tris-HCl, pH 7.5; 10 mM NaCl; 3 mM MgCl_2_ in nuclease free water:

| Reagent | Stock concentration | Final concentration | Volume (ml) |
| --- | --- | --- | --- |
| Tris-HCl (pH 7.5) | 1 M | 10 mM | 5 |
| NaCl | 5 M | 10 mM | 1 |
| MgCl_2_ | 1 M | 3 mM | 1.5 |
| Nuclease-free water | NA | NA | 492.5 |
| Final volume |  |  | 500 |

Filter the buffer through a 0.22 µm filter and store the buffer in 4C for up to 1 year.

2. **Nuclei Isolation Buffer (NIB)**

Nuclei buffer with 0.1% (volume) Tween-20 and 1X Protease Inhibitor Cocktail. To make a 25X Protease Inhibitor solution, dissolve 1 tablet of cOmplete™, EDTA-free Protease Inhibitor Cocktail into 2 mL nuclease free water.

3. **Nuclei Isolation Buffer + 0.1% IGEPAL-630**

Nuclei Isolation Buffer with 0.1% (volume) IGEPAL-630. For each sample, combine 1980 μL Nuclei Isolation Buffer and 20 μL 10% IGEPAL-630.

4. **2X Tagmentation Buffer (Stored in -20 °C)**

Prepare 200 mL of Tagmentation Buffer (filtered):

- 1M Tris HCl (pH 7.5): 4mL
- 1M MgCl_2_: 2mL
- DMF: 40mL
- H_2_O: 154mL

Aliquot the solution into 15 mL or 1.5 mL tubes for storage at -20 °C.

5. **2X Stop Buffer (Prepared fresh everytime, ~1.5 mL needed for one 96-well plate )**

| Reagent | Stock Con. | Volume |
| --- | --- | --- |
| EDTA | 40 mM | 400μL |
| Spermidine |  | 0.78μL |
| Nuclease free water |  | 4.6 mL |

6. **Tn5 storage buffer**

50 mM Tris-HCl, pH 7.45; 800 mM NaCl; 0.2 mM EDTA; 2 mM DTT; 10% glycerol.

Filter the buffer through a 0.22 µm filter, aliquot into 15mL/1.5 mL tubes, and store the buffer in -20 °C.

| Reagent | Stock concentration | Final concentration | Volume (ml) |
| --- | --- | --- | --- |
| Tris-HCl (pH 7.5) | 1 M | 50 mM | 5 |
| NaCl | 5 M | 800 mM | 16 |
| EDTA | 40 mM | 0.2 mM | 0.5 |
| DTT | 500 mM | 2 mM | 0.4 |
| Glycerol |  | 10% (volume) | 10 |
| Nuclease-free water |  |  | 68.1 |
| Final Volume |  |  | 100 |

**Ligation Primer Loading (1 hour)**

7. Resuspend and dissolve the Ligation Adaptor Primer Oligo to 100 μM in TE Buffer.
8. In each well of an empty 96-well plate, add 5 μL of 100 μM dissolved Ligation Adaptor Primer and 5 μL 100 μM Barcoded Ligation Primers - make sure to add the Barcoded Ligation Primers to their correct wells.
9. Anneal the adaptor and ligation primers together by running the following thermocycler program:

- 95 °C for 2 minutes
- Cool to 20 °C at a rate of -1 °C per minute
- Hold at 4 °C

The final annealed concentration will be 50 μM.

10. Dilute the primers to 3.3 μM by adding 140 μL of TE buffer. The resulting product is in stable, double-stranded form and can be stored at 4 °C or frozen. In 4 °C, the annealed primers should be stable for roughly three months and are suitable for short-term testing experiments.

**Tn5 Loading (2 hours)**

11. Protocol Derived from Hennig et al. 2018, Large-Scale Low-Cost NGS Library Preparation Using a Robust Tn5 Purification and Tagmentation Protocol - purified Tn5 protein is also generated following this publication. The Tn5 loading procedure is listed below. Volumes are calculated to make 4 96-well barcoded Tn5 plates.
1) For each N5 barcoded oligo, mix the following:

- 11.52 μL 100 μM N5 barcoded oligo
- 11.52 μL 100 μM MEr oligo
- 48.96 μL Tn5 Storage buffer

2) For each N7 barcoded oligos, mix the following:

- 5.76 μL 100 μM N7 barcoded oligo
- 5.76 μL 100 μM MEr oligo
- 24.48 μL Tn5 Storage buffer

3) Run the following PCR program in a thermocycler for the annealing of the oligonucleotides:

- 95°C 5 min --> slowly cool down to 65 °C (0.1°C/sec or 2%)
- 65°C 5 min --> slowly cool down to 4 °C (0.1°C/sec or 2%)
- 4°C

4) Plate indexed oligos in 96-well plate:

- Mix 2.5 μL of N5/MEr (16 μM) and 2.5 μL of N7/MEr (16 μM) in each well.
- Result: 5 μl of 8 μM Tn5 oligo mix.

5) Prepare glycerol Tn5 stock:
   - Thaw 600 μL Tn5 extract on ice (4mg/ml original concentration)
   - Prepare 1800 μL glycerol buffer: 600ul Tn5 Storage buffer + 1200 ul 100% glycerol (800 mM NaCl) + 7.2 μL 500 mM DTT (final DTT concentration at 2 mM).
   - Add the thawed Tn5 to the glycerol buffer and mix by gentle rotation at 4 °C.
   - Once it is mostly mixed together, use a pipet to make it completely homogeneous. and store at -20 °C until needed.

6) Load indexed Tn5:

- Add 5.5 μL of 8 μM Tn5 stock (1 mg/ml) to each well, mix by pipetting gently.
- Incubate at room temperature for 30 minutes with gentle shaking at 300 RPM.
- Centrifuge the plates and store at -20 °C until needed.

**Tissue preparation and nuclei isolations (~2 hours for 4 samples)**

11. Cool centrifuge to 4 °C - make sure to use a bucket centrifuge for all centrifuging steps unless otherwise stated, as normal centrifuges may have difficulty making a neat pellet at the bottom of the tube, which is necessary to maximize nuclear recovery. In a 6-cm dish on ice, cut each tissue section (0.1 g - 0.5 g) into small pieces (< 1 mm^3^) using a razor blade (pre-chilled) in 1 mL PBS on ice. Transfer the tissue and solution into a 1.5 mL tube and spin for 5 minutes at 200g at 4 °C.

NOTES:

- Do not process too many samples at the same time to reduce over-lysis.
- Do not let the samples stand on ice for too long, start the nuclei extraction immediately after taking them out of the freezer.

12. Dump Supernatant and add 1 mL ice-cold NIB + 0.1% IGEPAL CA-630 to the tissue for nuclei extraction. Pipet the tissue up and down with a 1 mL pipet tip 10 times (cut the top of 1 mL pipet tip if needed for easier pipetting). Incubate on ice for 5 minutes.

NOTES:

- From this point on, use 1 mL pipet tips or wide bore tips when working with nuclei to avoid stress on nuclei.

13. Filter tissue with a 40 μm cell strainer into a 6 cm dish and grind tissue on the strainer using a 5 mL syringe plunger. Add another 500 µL NIB + 0.1% IGEPAL CA-630 and continue grinding tissue on the strainer. Move solution into a 1.5 mL microcentrifuge tube.

NOTES:

It is not necessary to push the whole tissue through the filter! Use your own discretion in deciding when to stop grinding the tissue through the filter, but make sure not to tear through the filter!

14. Transfer the isolated nuclei into 1.5 mL Lo-bind tubes and centrifuge at 500g, 4°C for 5 minutes.
15. Remove supernatants and wash nuclei with 1 mL NIB and centrifuge at 500g, 4°C for 5 minutes.
16. Remove supernatants and resuspend nuclei in ~900 µL NIB (final volume will be higher).
17. Take a few µL samples for counting using nuclear staining (e.g. DAPI). Adjust final concentration to ~5,000-10,000 nuclei/µL.
18. PERFORM THIS STEP IF YOU WISH TO STORE NUCLEI FOR LATER USE - OTHERWISE, SKIP TO THE SECOND PART OF THE NEXT STEP:

For long-term storage, resuspend nuclei in 100-500 μL NIB + 10% DMSO, and split them into 100 μL aliquots into 1.5 mL cryo tubes. Place into slow-freeze FreezeCell containers and stored at -80°C. Nuclei should be stored in liquid nitrogen to reduce degradation due to temperature fluctuations. (**STOP POINT**).

**Nuclei Wash and filtering (~15 minutes for 4 samples)**

19. PERFORM BELOW IF YOU ARE WORKING WITH PREVIOUSLY FROZEN, STORED NUCLEI:

- Transfer nuclei from liquid nitrogen to 37°C water bath to thaw as quickly as possible. Place on ice immediately after being completely thawed.
- Add 1000 µL NIB to each sample for washing and mix gently by vortexing.
- Centrifuge nuclei at 500g, 4°C for 5 minutes, and remove supernatant by pipetting.

20. Add ~500 µL NIB and filter through a pluriStrainer Mini 40 µm filter into clean 1.5 mL LoBind tubes. Briefly centrifuge at 300g, 4°C for 15 seconds to let all the nuclei pass through.
21. Resuspend the nuclei in 100μL NIB.

**Nuclei counting (~30 minutes for 4 samples)**

22. Count the nuclei concentration for each sample.

Optimally, use a buffer with DAPI and a fluorescent microscope to distinguish between actual nuclei and debris.

Dissolve 10 mg DAPI in 2 ml of deionized water (dH2O) with a final concentration of 5 mg/ml. Split the DAPI solution into multiple tubes (100 ul per tube).

Take out one tube (100 ul, 5 mg/ml DAPI), add 1.9 ml deionized water (dH2O). Split the diluted DAPI solution into multiple tubes (100 ul per tube, 0.25 mg/ml DAPI).

Store the DAPI solution in the -20 °C freezer.

Make the DAPI counting solution: in 500 μL of Nuclei Buffer, add 0.5 μL - 1μL of 0.25mg/mL DAPI solution.

Take 1 μL of the sample and combine it with 9μL of the counting solution. Mix the solution and take 6 μL to dispense into a hemocytometer for counting the concentration of nuclei.

23. Adjust final concentration to 1000 nuclei/µL using NIB.

**Tagmentation (~1 hour for 4 plates)**

24. Dispense 5000 nuclei per well into 96-well plates by mixing the prepared nuclei with 2X TD buffer at 1:1 ratio.
    a. e.g. for each sample, mix 200 µL nuclei (1000 nuclei/µL) with 200 µL 2X TD buffer. Nuclei generally distributed into PCR strips and then distributed into wells - make sure not to pipet up and down in 96-plate wells to avoid introducing extra stress to the nuclei
25. Add 1 µL barcoded Tn5 enzyme in each well and cover with plastic foil.
26. Perform the tagementation at 55°C for 10 minutes with gentle shaking at 300 RPM.
27. Briefly spin down the plates (15 seconds) to collect all liquid from the sides.
28. Place on ice immediately to prevent over-tagmentation. Add 11 µL of 2X Stop Buffer. Mix gently by pipetting.

**Pool/Centrifuge/Resuspend/Redistribute (30 minutes)**

29. Pool nuclei and transfer them to 1.5 mL tubes.
30. Centrifuge the nuclei at 500g, 4°C for 5 minutes. Remove supernatant gently by pipetting. Leave a little excess to save as much nuclei as possible.
31. Wash nuclei with 1 mL NIB.
32. Centrifuge nuclei at 500g, 4°C for 5 minutes. Remove supernatant gently by pipetting.
33. Resuspend nuclei in 2150 µL NIB (1920 µL + ~10% excess for pipetting). Mix gently by vortexing.
    a. Notes:

- Nuclei concentration can be counted at this step for sample loss calculations.

34. Load 5 µL resuspend nuclei (~5000 nuclei) into each well in 96-well plates.
    a. Note: use 20 µL low retention tips, not wide-bore tips for higher pipetting accuracy.

**Ligation (~2 hours for 4 plates)**

35. To each well, add 2 µL DNA ligation primer/adaptor complex (3.3 μM).
36. To each well, add 3 µL T4 DNA ligase master mix in each well containing the following:
    a. 1 µL 10X T4 DNA Ligase Buffer
    b. 1 µL T4 DNA Ligase Enzyme
    c. 1 µL nuclease-free water
37. Cover plates with metal foil, and centrifuge briefly at 300g, 4°C for 15 seconds to collect all liquid from the sides.
38. Incubate plates for 30 minutes at room temperature with gentle shaking (300 rpm with Thermomixer, 50 rpm on Fisherbrand Nutating Mixer).
39. Place the plate on ice immediately and centrifuge briefly at 300g, 4°C for 15 seconds to collect all liquid from the sides.
40. Add 2 µL 18 mM EDTA to each well to stop the ligation reaction, pool all samples and transfer to 1.5 mL LoBind tubes.

**Pool/Centrifuge/Resuspend/Redistribute/Quantify (30 minutes)**

41. Centrifuge nuclei at 500g, 4°C for 5 minutes. Remove supernatant gently by pipetting. Leave a little excess to save as much nuclei as possible.
42. Wash nuclei with 1 mL NIB.
43. Centrifuge the tube at 500g, 4°C for 5 minutes. Remove supernatant gently by pipetting.
44. (Optional) Nuclei filtering.
    a. Resuspend nuclei in ~600 µL NIB and transfer them into a clean 1.5 mL LoBind tube.
    b. Filter the nuclei using a Pluristrainer Mini 5µm filter.
    c. Centrifuge nuclei briefly at 300g, 4°C for 15 seconds to allow all the nuclei to pass though.
    d. Wash the filter again 3 times with 100 µL NIB , followed by centrifugation at 300g, 4°C for 15 seconds to improve recovery rate. (~900 µL final sample)
45. Resuspend the nuclei in ~100 µL NIB, count nuclei and adjust the concentration to ~2000 nuclei/µL.
46. Distribute the nuclei into a 96 well plate with 10,000 nuclei per well.

NOTES:

- You can directly freeze and store cells at this point.
- If you choose to freeze the plate, it is okay to place it directly in -80 °C freezer without flash-freezing
- You can also store nuclei directly into PCR strips if you don't need to profile a whole plate of cells

**Proteinase K treatment (overnight)**

47. For each well, add 1 µL Proteinase K master mix:

- 0.5 µL EB buffer

- 0.25 µL 1% SDS

- 0.25 µL Proteinase K (18.9 mg/ml)

48. Mix the samples by vortexing, followed by a brief centrifugation to collect all liquid from the sides, and vortex again to mix up the nuclei from the bottom.
49. Incubate samples at 65°C for 16 hours and keep at 4 °C until the following step.
50. Add 2 µL 10% Tween-20 to quench the SDS and mix well by pipetting.

Note:

- Adjust final volume to 8 µL using EB if sample evaporation occured in certain wells.

- Samples can be stored at -20°C or -80°C until the PCR amplification step.

**PCR Amplification (~1 hour)**

51. Centrifuge the plates to collect all liquid from the sides.

Notes:

- We perform test qPCR runs on a few wells to determine appropriate PCR cycle number by adding dsGreen to the master mix.

- Expected optimal cycle number is around mid-exponential phase, but higher cycle number is safer to use in order to ensure enough material for Novaseq sequencing.

- Higher cycle number can increase yield, but it can also result in higher adapter dimer concentration.

52. To each well, add the following:

- 1 µL 10 µM Universal P5 primer

- 1 µL 10 µM Barcoded P7 primer

- 10 µL NEBNext® High-Fidelity 2X PCR Master Mix.

53. Mix wells by pipetting. Centrifuge to collect all liquid from sides and remove bubbles.
54. Perform the following PCR reaction:
    a. 72°C for 5 minutes
    b. 98°C for 30 seconds
    c. 8 cycles of (98°C for 10 seconds, 66°C for 30 seconds, 72°C for 30 seconds)
    d. 72°C for 5 min
55. Store samples at -20°C until the sequencing library purification step (STOP POINT).

**Sequencing Library Purification (1 hour)**

56. Combine PCR products into a 1.5 mL tube.
57. Perform column purification using Zymo DNA Clean & Concentrator kit. Elute in 20 µL EB.
58. Run samples on 2% E-Gel™ EX Agarose Gels. Perform gel purification using Zymo Gel DNA Recovery Kit to remove primer dimers. Elute in 16 µL EB.
59. Quantify the library concentration and visualize the library via electrophoresis (performed using a Qubit and a 2% Agarose E-Gel). An example library is shown below:

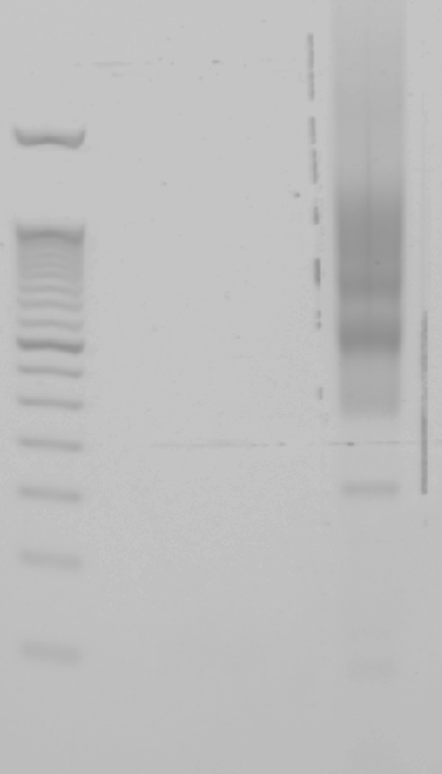

Lanes from left to right:

1. E-Gel™ 50 bp DNA Ladder
2. Pooled PCR product after column purification (Gel extraction will be performed afterwards to get rid of primer dimer at ~150bp)
60. Prepare Illumina sequencing library depending on the instrument recommendations:

- NextSeq 1000: 2 nM DNA library, minimum 12 µL

- NovaSeq 6000: 2.6 nM DNA library, minimum 310 µL

Important: To increase library complexity, add minimum 20% PhiX before sequencing.

61. Sequence the library on the Novaseq Platform (Read1:100 bp, Read2: 100 bp, Index 1: 10 bp, Index 2: 10 bp).

NOTE: Perform a small-scale sequencing run before NovaSeq sequencing to prevent possible clustering and quality issues.

**References**

Hennig, Bianca P., Lars Velten, Ines Racke, Chelsea Szu Tu, Matthias Thoms, Vladimir Rybin, Hüseyin Besir, Kim Remans, and Lars M. Steinmetz. 2018. “Large-Scale Low-Cost NGS Library Preparation Using a Robust Tn5 Purification and Tagmentation Protocol.” *G3* 8 (1): 79–89.
