## Supplementary file 6 for "A global view of aging and Alzheimer’s pathogenesis-associated cell population dynamics and molecular signatures in the human and mouse brains": EasySci_ATAC_pipeline.sh

```
########################################################## MODIFY THIS
fastq_folder=""
sample_ID=""
all_output_folder=""
PCR_sample_barcode_file=''
########################################################## MODIFY THIS


core=16 # Define number of cores used 
# define the number of unique reads cutoff for splitting single cell; cells with unique reads less than this number will be discarded
cutoff=500

###Choose the mouse STAR index, GTF file, GFF file
index="/ru-auth/local/home/asziraki/projects/AS_20200820_single_cell_pipeline/Raw_data/AS_20200820_genome_files/original_pipeline/Reference_ALL/index/index/STAR/mouse-latest-gencode-release-m27/STAR_mouse_M27"
gtf_file="/ru-auth/local/home/asziraki/projects/AS_20200820_single_cell_pipeline/Raw_data/AS_20200820_genome_files/original_pipeline/Reference_ALL/index/GTF/mouse-latest-gencode-release-m27/gencode.vM27.primary_assembly.annotation.gtf.gz"
gff_file='/rugpfs/fs0/cao_lab/store/zlu/Reference/GTF/gencode.vM27.primary_assembly.annotation.gff3.gz'
chromosome_length_file=$index/chrNameLength.txt

################ STEP 1: Barcode extraction, trimming, alignment, filter low quality alignments and remove duplicates

################ Barcode extraction(Extract ligation barcdoe, N5 and N7 barcode and attach those to the reads name)
script_folder='/Scripts/'
input_folder=$fastq_folder
output_folder=$all_output_folder/UMI_attach
script=$script_folder/UMI_attach_EasySci_ATAC3.py
echo "Changing the name of the fastq files..."
for sample in $(cat $sample_ID); do echo changing name $sample; mv $input_folder/*$sample*R1*fastq.gz $input_folder/$sample.R1.fastq.gz; mv $input_folder/*$sample*R2*fastq.gz $input_folder/$sample.R2.fastq.gz; mv $input_folder/*$sample*R3*fastq.gz $input_folder/$sample.R3.fastq.gz; done
echo "Attaching barcode...."
mkdir -p $output_folder
barcode_N5=$script_folder/HPLC_BC_N5.pickle2
barcode_N7=$script_folder/HPLC_BC_N7.pickle2
ligation_barcode=$script_folder/EasySci-ATAC_P5_BC.pickle2
source /rugpfs/fs0/cao_lab/scratch/zlu/anaconda3/etc/profile.d/conda.sh
conda activate sc-01
python $script $input_folder $sample_ID $output_folder $barcode_N5 $barcode_N7 $ligation_barcode $core
echo "Barcode transformed"
conda deactivate

################# Trimming the reads
bashrc_location=/ru-auth/local/home/asziraki/projects/AS_20200820_single_cell_pipeline/Scripts/original_pipeline/sci-RNA-seq3_pipeline/.bashrc
source $bashrc_location
conda activate ATAC_pipeline
echo "Start trimming the read file..."
mkdir -p $all_output_folder/trimmed_fastq
trimmed_fastq=$all_output_folder/trimmed_fastq
for sample in $(cat $sample_ID); do echo trimming $sample; trim_galore $all_output_folder/UMI_attach/$sample.R1.fastq.gz $all_output_folder/UMI_attach/$sample.R2.fastq.gz --paired -a CTGTCTCTTATACACATCT -o $trimmed_fastq; done
echo "All trimmed file generated."

############ Align the reads with STAR
input_folder=$trimmed_fastq
STAR_output_folder=$all_output_folder/STAR_alignment
filtered_sam_folder=$all_output_folder/filtered_sam
rmdup_sam_folder=$all_output_folder/rmdup_sam
#align read1, read2 to the index file using STAR with default setting
echo "Start alignment using STAR..."
mkdir -p $STAR_output_folder
STAR --genomeDir $index --genomeLoad Remove
#start the alignment
for sample in $(cat $sample_ID); do echo aligning $sample; STAR --runThreadN $core --outSAMstrandField intronMotif --genomeDir $index --readFilesCommand zcat --readFilesIn $input_folder/$sample*R1*.gz $input_folder/$sample*R2*.gz --outFileNamePrefix $STAR_output_folder/$sample --genomeLoad LoadAndKeep; done
#remove the index from the memory
STAR --genomeDir $index --genomeLoad Remove
echo "All alignment done."

############ Filter the alinged reads

echo "Start filter and sort the sam files..."
mkdir -p $filtered_sam_folder
for sample in $(cat $sample_ID); do echo Filtering $sample; \
samtools view -H $STAR_output_folder/$sample*.sam > $filtered_sam_folder/$sample.sam;\
samtools view -bh -q 30 -F 4 -f 0x2 -@ 10 -F 0x100 $STAR_output_folder/$sample*.sam|samtools sort -@ 10 -|samtools view -|awk '$3 ~ /^chr/' ->>$filtered_sam_folder/$sample.sam; \
done

################################# remove duplicates using Picard
echo Remove duplicates
mkdir -p $rmdup_sam_folder
for sample in $(cat $sample_ID); do echo removing duplicates $sample; picard MarkDuplicates INPUT=$filtered_sam_folder/$sample.sam OUTPUT=$rmdup_sam_folder/$sample.bam REMOVE_DUPLICATES=true ASSUME_SORTED=True METRICS_FILE=$rmdup_sam_folder/Picard_metrics_file.$sample.txt VALIDATION_STRINGENCY=LENIENT; done

################################# calculate the reads number in each file

report_folder=$all_output_folder/report
mkdir -p $report_folder
fastq_folder=$fastq_folder
trimmed_folder=$all_output_folder/trimmed_fastq
alignment=$all_output_folder/STAR_alignment
filtered_sam=$all_output_folder/filtered_sam
rm_dup_sam=$all_output_folder/rmdup_sam
echo sample,Total reads,Barcode matching,After trimming,Uniquely aligned reads,Filtered reads,After remove duplicates>$report_folder/read_number.csv
for sample in $(cat $sample_ID); do echo calculating $sample; echo $sample,$(expr $(zcat $fastq_folder/$sample.R[13].fastq.gz|wc -l) / 4),$(expr $(zcat $all_output_folder/UMI_attach/$sample*.gz|wc -l) / 4),$(expr $(zcat $trimmed_folder/$sample*.gz|wc -l) / 4),$(samtools view $alignment/$sample*.sam|wc -l),$(samtools view $filtered_sam/$sample.sam|wc -l),$(samtools view $rm_dup_sam/$sample.bam|wc -l)>>$report_folder/read_number.csv; done
echo "Read number calculation is done."


################ STEP 2: Generate bedpe file, fragment files and single-cell read count file for downstream analysis

################ Sort sam files by readname and make bedpe and fragment files
echo Sort sam by readnames make bedpe and fragments files
sam_folder=$all_output_folder/rmdup_sam
sam_folder_ordered_by_read_name=$all_output_folder/rmdup_sam_ordered_by_read_name
fragmentfile_new=$all_output_folder/fragmentfile_new
mkdir -p $sam_folder_ordered_by_read_name
bash_script=$script_folder/EasySci_ATAC3_sam_order.sh
R_script=$script_folder/sci3_bash_input_ID_output_core.R
Rscript $R_script $bash_script $sam_folder $sample_ID $sam_folder_ordered_by_read_name $core
input_folder=$sam_folder_ordered_by_read_name
output_folder=$fragmentfile_new

mkdir -p $fragmentfile_new
for sample in $(cat $sample_ID);do echo bam to bedpe $sample; \
samtools view -S -b $input_folder/$sample.sam > $input_folder/$sample.bam; \
bedtools bamtobed -i $input_folder/$sample.bam -bedpe >  $input_folder/$sample.bedpe; \
done

for sample in $(cat $sample_ID);do echo bedpe to fragment $sample; \
PCRbarcode=$(cat $PCR_sample_barcode_file | grep $sample | cut -f 2); \
cat $input_folder/$sample.bedpe | cut -f 1,2,6,7 | awk '{sub (/,/, OFS)} 1' OFS='\t' | cut -f 1,2,3,4 | awk -v barcode=$PCRbarcode '{$2=$2+4;$3=$3-5;$4=(barcode)$4; print}' OFS='\t' | awk '$1 ~ /^chr/' | awk -F '\t' -v OFS='\t' '{$(NF+1)=1;print}' > $output_folder/$sample.fragments.bed; \
done
cat $fragmentfile_new/*.bed > $fragmentfile_new/fragments.bed
conda deactivate

source /rugpfs/fs0/cao_lab/scratch/zlu/anaconda3/etc/profile.d/conda.sh
conda activate R4.1
bgzip -@ 10 --force -c $fragmentfile_new/fragments.bed > $fragmentfile_new/fragments.bed.gz
conda deactivate 

########################### split the sam file based on the barcode, and move the result to the report folder
source $bashrc_location
conda activate ATAC_pipeline
echo Split sam files 
sam_folder=$all_output_folder/rmdup_sam_ordered_by_read_name
output_folder=$all_output_folder/sam_splitted
mkdir -p $output_folder


echo "Start splitting the sam file..."
echo samfile folder: $sam_folder
echo sample list: $sample_ID
echo ouput folder: $output_folder
## echo barcode file: $barcodes
echo cutoff value: $cutoff

bash_script=$script_folder/sci3_split_paired_end.sh
Rscript $R_script $bash_script $sam_folder $sample_ID $output_folder $core $cutoff
cat $output_folder/*sample_list.txt>$output_folder/All_samples.txt
cp $output_folder/All_samples.txt $output_folder/../barcode_samples.txt


################ STEP 3: Counting reads overlapping with promoter regions

##Make promoter reference bed files
reference_folder=$all_output_folder/reference_folder/
promoter_length_upstream=1000 ### Here you can modify the upstream and downstream distance for defining the promoter region
promoter_length_downstream=1000
python_use="/rugpfs/fs0/cao_lab/scratch/asziraki/anaconda3/bin/python"
mkdir -p $reference_folder
promoter_file=$reference_folder/promoter_gtf.bed
script=$script_folder/gtf_to_promoter_bed.r

bashrc_location=/ru-auth/local/home/asziraki/projects/AS_20200820_single_cell_pipeline/Scripts/original_pipeline/sci-RNA-seq3_pipeline/.bashrc
source $bashrc_location
conda activate ATAC_pipeline_createreference
Rscript $script $gtf_file $promoter_file $promoter_length_upstream $promoter_length_downstream
conda deactivate

conda activate ATAC_pipeline
bedtools sort -i $reference_folder/promoter_gtf.bed | bedtools merge -c 4 -o distinct -i - > $reference_folder/promoter_gtf.overlappingmerged.bed 
promoter_bed=$reference_folder/promoter_gtf.overlappingmerged.bed

all_output=$all_output_folder/peak_count
sam_folder=$all_output_folder/sam_splitted/
sample_list=$all_output_folder/barcode_samples.txt
echo Start promoter counting
out_folder=$all_output/peak_count_promoters/
mkdir -p $out_folder
ref_bed=$promoter_bed
dis=100
script=$script_folder/peak_map_new.py
python $script $sample_list $sam_folder $ref_bed $out_folder $dis $core
rm $out_folder/cell_count.MM
rm $out_folder/cell_nread.txt
for sample in $(cat $sample_list); do echo combining $sample; cat $out_folder/$sample.count >>$out_folder/cell_count.MM; rm $out_folder/$sample.count; done
for sample in $(cat $sample_list); do cat $out_folder/$sample.nread.txt >> $out_folder/cell_nread.txt; rm $out_folder/$sample.nread.txt; done
conda deactivate

conda activate ATAC_pipeline_createreference
peak_count_folder=$all_output/peak_count_promoters/
summary_folder=$all_output/summary_count_onlypromoter/
mkdir -p $summary_folder
script=$script_folder/peak_count_summary_no_report_promoter.r
Rscript $script $peak_count_folder $reference_folder $summary_folder
conda deactivate
echo Promoter counting finished

#### Count total reads per cell
rm $all_output_folder/total_read_per_cell.csv
for VAR in $(cat $all_output_folder/barcode_samples.txt);do echo $VAR,$(grep -v "@" $all_output_folder/sam_splitted/$VAR.sam | wc -l) >> $all_output_folder/total_read_per_cell.csv ;done

################ STEP 4: Generate h5ad file for SnapATAC2 analysis

source /rugpfs/fs0/cao_lab/scratch/zlu/anaconda3/etc/profile.d/conda.sh
conda activate snapatac2_old
python_script=$script_folder/snapATAC2_make_h5ad.py

working_folder=$all_output_folder/fragmentfile_new
fragment_file=$all_output_folder/fragmentfile_new/fragments.bed
min_tsse=2 ### Define the minumum TSS Enrichment score for pre-filtering
min_nfragment=1000 ### Define the minimum number of fragment for pre-filtering

echo start preparing h5ad for running snapatac2
python $python_script $working_folder $fragment_file $gff_file $min_tsse $min_nfragment $chromosome_length_file
conda deactivate
```
