## Supplementary file 6 for "A global view of aging and Alzheimer’s pathogenesis-associated cell population dynamics and molecular signatures in the human and mouse brains": generate_barcode_dictionary.py

```
N5_bc_file = "/Scripts/HPLC_N5_BC.txt"
N7_bc_file = "/Scripts/HPLC_N7_BC.txt"
lig_bc_file = "/Scripts/EasySci-ATAC_P5_BC.txt"


pickle_N5_temporary="/Scripts/HPLC_BC_N5.pickle"
pickle_N7_temporary="/Scripts/HPLC_BC_N7.pickle"
ligation_pickle_temporary = "/Scripts/EasySci-ATAC_P5_BC.pickle"

pickle_N5_final="/Scripts/HPLC_BC_N5.pickle2"
pickle_N7_final="/Scripts/HPLC_BC_N7.pickle2"
ligation_pickle_final = "/Scripts/EasySci-ATAC_P5_BC.pickle2"


import gzip
import itertools
from Levenshtein import distance
import pickle

# for accept a list of characters ["A", "C", "G", "T"] and a length generate all permutations 
def combination_seqs(input_char, output_length):
    all_combine = list(map(lambda x: list(map(lambda x: "".join(x), set(itertools.permutations("".join(x))))), 
                           list(itertools.combinations_with_replacement(input_char, output_length))))
    return(list(itertools.chain.from_iterable(all_combine)))

# Here I am going to accept the permuted sequence list and the target list and then generate
# a dictionary including the permuted sequence and the matching sequence (within 1 edit distance)
# if there is no sequence matching, then return ""

def find_bc(seq, bc_list, mismatch_rate = 1):
    # in this script, I am going to accept a sequence and a ligation barcode list, and then
    # find if there is a matching barcode in the bc_list matching the first several bases of
    # the seq
    result = ""
    for bc in bc_list:
        mismatch = distance(bc, seq)
        if mismatch <= mismatch_rate:
            result = bc
            break
    return(result)


def find_lig_bc(x, lib_dic):
    if x in lib_dic.keys():
        return(lib_dic[x])
    else:
        return(x)

# add "C" to 9bp ligation barcode
def add_c(x):
    if len(x) == 9:
        return(x+"C")
    else:
        return(x)
    
# generate all possible sequences of 6bp
print("Generating all possible sequences...")
all_seq_6bp = combination_seqs(["A", "C", "G", "T", "N"], 6)
N5_bc = open(N5_bc_file)
N5 = list(map(lambda x: x.strip(), N5_bc.readlines()))
N5_bc.close()

N7_bc = open(N7_bc_file)
N7 = list(map(lambda x: x.strip(), N7_bc.readlines()))
N7_bc.close()

print("Generating the dictionary for N5 barcodes...")
# generat the dictionary for RT barcodes
all_seq_match = list(map(lambda x: find_bc(x, N5), all_seq_6bp))
all_seq_dic = dict(zip(all_seq_6bp, all_seq_match))
# save the dictionary into a file
pickle_out = open(pickle_N5_temporary,"wb")
pickle.dump(all_seq_dic, pickle_out)
pickle_out.close()

print("Generating the dictionary for N7 barcodes...")
# generat the dictionary for RT barcodes
all_seq_match = list(map(lambda x: find_bc(x, N7), all_seq_6bp))
all_seq_dic = dict(zip(all_seq_6bp, all_seq_match))
# save the dictionary into a file
pickle_out = open(pickle_N7_temporary,"wb")
pickle.dump(all_seq_dic, pickle_out)
pickle_out.close()


pickle_in = open(pickle_N5_temporary, "rb")
N5_dic = pickle.load(pickle_in)
N5_filter = {k: v for k, v in N5_dic.items() if v != ""}
pickle_out = open(pickle_N5_final, "wb")
pickle.dump(N5_filter, pickle_out, 2)
pickle_in.close()
pickle_out.close()

pickle_in = open(pickle_N7_temporary, "rb")
N7_dic = pickle.load(pickle_in)
N7_filter = {k: v for k, v in N7_dic.items() if v != ""}
pickle_out = open(pickle_N7_final, "wb")
pickle.dump(N7_filter, pickle_out, 2)
pickle_in.close()
pickle_out.close()


# generate all possible sequences of 10bp
print("Generating all possible sequences...")
all_seq_10bp = combination_seqs(["A", "C", "G", "T", "N"], 10)
# save the dictionary into a file
print("Generating the dictionary for ligation barcodes...")
# also generate the dictionary for ligation barcodes
lig_bc = open(lig_bc_file)
lig = list(map(lambda x: x.strip(), lig_bc.readlines()))
lig_bc.close()
lib_bc_addC = list(map(lambda x: add_c(x), lig))
lib_dic = dict(zip(lib_bc_addC, lig))
all_seq_match = list(map(lambda x: find_lig_bc(find_bc(x, lib_bc_addC), lib_dic), all_seq_10bp))
all_seq_dic = dict(zip(all_seq_10bp, all_seq_match))
pickle_out = open(ligation_pickle_temporary,"wb")
pickle.dump(all_seq_dic, pickle_out)
pickle_out.close()

# generate the simplified dictionary
pickle_in = open(ligation_pickle_temporary, "rb")
lig_dic = pickle.load(pickle_in)
lig_filter = {k: v for k, v in lig_dic.items() if v != ""}
pickle_out = open(ligation_pickle_final, "wb")
pickle.dump(lig_filter, pickle_out, 2)
pickle_in.close()
pickle_out.close()
```
