## Supplementary file 6 for "A global view of aging and Alzheimer’s pathogenesis-associated cell population dynamics and molecular signatures in the human and mouse brains": sam_split_paired_end.py

```
'''
Modified by Andras Sziraki, 10/01/2021: to speed up the script, only works with the read name sorted sam file implemented in the paired end pipeline
in this script, I will read into the sam file and the barcode file, and then count the reads number per
barcode
'''
import sys
import matplotlib as mpl
mpl.use('Agg')
import matplotlib.pyplot as plt
from matplotlib.backends.backend_pdf import PdfPages

def split_samfile(sam_file, output_folder, cutoff):
     
    sam = open(sam_file)
    pre_barcode = 'EMPTY'
    plot_name = (sam_file.split('/')[-1]).split('.')[0]
    sample_list_file = open(output_folder + '/' + plot_name + '.' + 'sample_list.txt', 'w')
    header = []
    alignments = []
    for line in sam:
        if (line[0] == '@'):
            header.append(line)
        else:
            barcode = (((line.split('\t'))[0]).split(','))[0]
            if barcode == pre_barcode:
                alignments.append(line)
            else:
                if ( len(alignments) >= cutoff):
                    output_file = output_folder + '/' + plot_name + '.' + pre_barcode + '.sam'
                    file = open(output_file, 'a')
                    for element in header:
                        file.write(element)
                    for alignment in alignments:
                        file.write(alignment)
                    file.close()
                    sample_list_file.write(plot_name + '.' + pre_barcode + '\n')
                alignments = []
                alignments.append(line)
                pre_barcode = barcode
    
    ## write out last cell
    if ( len(alignments) >= cutoff):
        output_file = output_folder + '/' + plot_name + '.' + pre_barcode + '.sam'
        file = open(output_file, 'a')
        for element in header:
            file.write(element)
        for alignment in alignments:
            file.write(alignment)
        file.close()
        sample_list_file.write(plot_name + '.' + pre_barcode + '\n')
    
    ## close files
    sample_list_file.close()
    sam.close()

if __name__ == '__main__':
    sam_file = sys.argv[1]
    output_folder = sys.argv[2]
    cutoff = int(sys.argv[3])
    split_samfile(sam_file, output_folder, cutoff)
```
