## Supplementary file 6 for "A global view of aging and Alzheimer’s pathogenesis-associated cell population dynamics and molecular signatures in the human and mouse brains": sci3_split_paired_end.sh

```
input_folder=$1
sample=$2
output_folder=$3
cutoff=$4

python="/rugpfs/fs0/cao_lab/scratch/asziraki/anaconda3/envs/original_pipeline/bin/python2.7"
python_script="//rugpfs/fs0/cao_lab/store/zlu/ZL_20220212_Mouse_brain_aging_atlas/Scripts/EasySci_ATAC3_Scripts//sam_split_paired_end.py"

$python $python_script $input_folder/$sample.sam $output_folder $cutoff
echo splitting sample done: $sample
```
