## Supplementary file 6 for "A global view of aging and Alzheimer’s pathogenesis-associated cell population dynamics and molecular signatures in the human and mouse brains": snapATAC2_make_h5ad.py

```
import sys
working_folder = sys.argv[1]
fragment_file = sys.argv[2]
gff_file=sys.argv[3]
min_tsse=int(sys.argv[4])
min_nfragments=int(sys.argv[5])
chr_name_file=str(sys.argv[6])

print("working_folder",working_folder)
print("fragment_file",fragment_file)
print("gff_file",gff_file)
print("min_tsse",str(min_tsse))
print("min_nfragments",str(min_nfragments))

import snapatac2 as snap
import pandas as pd
import os 
import numpy as np
import seaborn as sns
import matplotlib.pyplot as plt

os.chdir(working_folder)
gene_annotation = gff_file
# gene_annotation = "/rugpfs/fs0/cao_lab/scratch/zlu/ZL_20220212_Mouse_brain_aging_atlas/Intermediate_data/220324_Novaseq_snapATAC2/fragmentfile/gencode.vM28.annotation.gff3.gz"
chr_name = pd.read_csv(chr_name_file,sep='\t',header=None)
mm39_chr_length = dict(zip(chr_name.iloc[:22,0],chr_name.iloc[:22,1]))
adata = snap.pp.import_data(
    fragment_file,
    gene_annotation,
    mm39_chr_length,
    sorted_by_barcode=False,
    n_jobs=32,
    min_num_fragments=min_nfragments,
    min_tsse=min_tsse,
    output= 'adata.h5ad',   
)
snap.pp.make_tile_matrix(adata)

adata_r = snap.read_h5ad("adata.h5ad",backed='r+')
gene_matrix = snap.pp.make_gene_matrix(adata_r, gff_file=gene_annotation,backed='r+')
```
