## Supplementary file 6 for "A global view of aging and Alzheimer’s pathogenesis-associated cell population dynamics and molecular signatures in the human and mouse brains": UMI_attach_EasySci_ATAC3.py

```
import subprocess
import sys
from Levenshtein import distance
import gzip
from multiprocessing import Pool
from functools import partial
import pickle


def UMI_attach_read2_barcode_list(sample, input_folder, output_folder, RT_barcode_list_N5, RT_barcode_list_N7,ligation_barcode_list, mismatch_rate = 1):

    Read1 = input_folder + "/" + sample + ".R1.fastq.gz"
    Read2 = input_folder + "/" + sample + ".R3.fastq.gz"
    Read3 = input_folder + "/" + sample + ".R2.fastq.gz"

    output_file1 = output_folder + "/" + sample + ".R1.fastq.gz"
    output_file2 = output_folder + "/" + sample + ".R2.fastq.gz"

    f1 = gzip.open(Read1)
    f2 = gzip.open(Read2)
    I5_reads = gzip.open(Read3)

    f3 = gzip.open(output_file1, 'wb')
    f4 = gzip.open(output_file2, 'wb')

    line1 = f1.readline()
    line2 = f2.readline()
    line3 = I5_reads.readline()
    total_line = 0
    filtered_line = 0
    ligation_matched_line  = 0 
    while (line1):
        total_line += 1
        line1_header = line1
        line2_header = line2
        line1 = f1.readline()
        line2 = f2.readline()
        line3 = I5_reads.readline()
            # first check if the ligation barcode match with the barcode
        tmp_lig = line3[0:10]
        if tmp_lig.decode() in ligation_barcode_list:
            ligation_matched_line += 1
            ligation_bc_match = ligation_barcode_list[tmp_lig.decode()]
            BC1 = line1[0:6].decode()
            if BC1 in RT_barcode_list_N5:
                BC1_match = RT_barcode_list_N5[BC1]
                # check Tagmentation Read2 barcode
                BC2 = line2[0:6].decode()

                if BC2 in RT_barcode_list_N7:
                    BC2_match = RT_barcode_list_N7[BC2]
                    filtered_line += 1
                    first_line = '@' + ligation_bc_match + BC1_match + BC2_match + ',' +line1_header[1:].decode()
                    f3.write(first_line.encode())
                    first_line = '@' + ligation_bc_match + BC1_match + BC2_match + ',' +line2_header[1:].decode()
                    f4.write(first_line.encode())
                    
                    second_line = line1[25:]
                    f3.write(second_line)
                    second_line = line2[25:]
                    f4.write(second_line)
                    
                    third_line = f1.readline()
                    f3.write(third_line)
                    third_line = f2.readline()
                    f4.write(third_line)
                    
                    four_line = f1.readline()
                    four_line2 = four_line[25:]
                    f3.write(four_line2)
                    four_line = f2.readline()
                    four_line2 = four_line[25:]
                    f4.write(four_line2)
                    
                    line1 = f1.readline()
                    line2 = f2.readline()
                else:
                    line1 = f1.readline()
                    line1 = f1.readline()
                    line1 = f1.readline()
                    line2 = f2.readline()
                    line2 = f2.readline()
                    line2 = f2.readline()
                    
            else:
                line1 = f1.readline()
                line1 = f1.readline()
                line1 = f1.readline()
                line2 = f2.readline()
                line2 = f2.readline()
                line2 = f2.readline()
        else:      
            line1 = f1.readline()
            line1 = f1.readline()
            line1 = f1.readline()
            line2 = f2.readline()
            line2 = f2.readline()
            line2 = f2.readline()
        line3 = I5_reads.readline() 
        line3 = I5_reads.readline()
        line3 = I5_reads.readline()
    f1.close()
    f2.close()
    f3.close()
    f4.close()
    print("sample name: %s, total line: %f, ligation matched line: %f, \
          filtered line: %f, filter rate: %f" 
            %(sample, total_line,ligation_matched_line, filtered_line, float(filtered_line) / float(total_line)))


def attach_UMI_files(input_folder, sampleID, output_folder, RT_barcode_file_N5, RT_barcode_file_N7, ligation_barcode_file,core):
    
    init_message = '''
    --------------------------start attaching UMI-----------------------------
    input folder: %s
    sample ID: %s
    output_folder: %s
    RT barcode file N5: %s
    RT barcode file N7: %s
    ligation barcode file: %s
    ___________________________________________________________________________
    ''' %(input_folder, sampleID, output_folder, RT_barcode_file_N5, RT_barcode_file_N7,ligation_barcode_file)
    
    print(init_message)
    
    
    print("Load N5/N7 barcode dictionary...")
    
    barcodes = open(RT_barcode_file_N5, "rb")
    RT_barcode_list_N5 = pickle.load(barcodes)
    barcodes.close()

    barcodes = open(RT_barcode_file_N7, "rb")
    RT_barcode_list_N7 = pickle.load(barcodes)
    barcodes.close()

    print("Load ligation barcode dictionary...")
    
    # generate the ligation barcode list
    barcodes = open(ligation_barcode_file, "rb")
    ligation_barcode_list = pickle.load(barcodes)
    barcodes.close()

    #for each sample in the sample list, use the read1 file, read2 file, output file
    # and barcode_list to run UMI_attach_read2_barcode_list
    sample_file = open(sampleID)
    sample_list = []
    for line in sample_file:
        sample = line.strip()
        sample_list.append(sample)
    sample_file.close()
    
    # parallele for the functions
    p = Pool(processes = int(core))
    #print("Processing core number: ", core_number)
    func = partial(UMI_attach_read2_barcode_list, input_folder = input_folder, output_folder=output_folder, RT_barcode_list_N5=RT_barcode_list_N5, RT_barcode_list_N7=RT_barcode_list_N7, 
        ligation_barcode_list=ligation_barcode_list, mismatch_rate = 1)
    result = p.map(func, sample_list)
    p.close()
    p.join()
    
    #print the completion message
    com_message = '''~~~~~~~~~~~~~~~UMI attachment done~~~~~~~~~~~~~~~~~~'''
    print(com_message)
    
if __name__ == "__main__":
    input_folder = sys.argv[1]
    sampleID = sys.argv[2]
    output_folder = sys.argv[3]
    RT_barcode_file_N5 = sys.argv[4]
    RT_barcode_file_N7 = sys.argv[5]
    ligation_barcode_file = sys.argv[6]
    core=sys.argv[7]
    attach_UMI_files(input_folder, sampleID, output_folder, RT_barcode_file_N5, RT_barcode_file_N7,ligation_barcode_file, core)
```
