## Supplementary file 3 for "A global view of aging and Alzheimer’s pathogenesis-associated cell population dynamics and molecular signatures in the human and mouse brains": EasySci_main.sh

```
#! /bin/bash
#SBATCH -N 1
## number of cores 
#SBATCH -n 24
#SBATCH --partition=hpc
#SBATCH -o ./log_file.out

## Location of the bashrc file to activate the conda environments.
bashrc_location=
source $bashrc_location
conda activate original_pipeline
# 'original_pipeline' conda environment is necessarily to run the pipeline
# the EasySci-RNA pipeline accept a input folder, and then use the default parameter for sequencing processing and generating the gene count matrix for downstream analysis

# The script is for processing EasySci-RNA sequencing reads. For running in other environment, some modules, R and python packages are needed to be installed.

# Define the location of the sub script folder
script_path=./script_folder/


###################### INPUT PARAMETERS START

# define the fastq folder including all fastq files
fastq_folder=""

# define the PCR group sample id for each fastq file
sample_ID=""

# define the output folder
all_output_folder=""

# define the core number for parallele processing
core=24
# define the core number for reads filtering and sorting 
samtools_core=4

# define the number of UMI cutoff for splitting single cell; cells with UMIs less than this number will be discarded, default = 200
cutoff=200

# define the location of index files for reads alignment with STAR
# Mouse genome version M27
index=

# define the gtf file for gene counting
# Mouse genome version M27
gtf_file=

# define the gtf file for exon counting
# Mouse genome version M27
gtf_file_exon=

# define the location of the ligation barcodes
ligation_barcode=$script_path/Ligation_barcodes.pickle2
# define the location of the RT barcodes
RT_barcode=$script_path/RT_barcodes.pickle2
# define the location of the randomN RT barcodes for the gene counting step
randomN_barcode_file=$script_path/RandomN_RT_barcodes.txt

################################## INPUT PARAMETERS END

# define the location of the R script for multi-core processing
R_script=$script_path/paralell.R


now=$(date)
echo "Current time : $now"

############ UMI attach
# the script take an input folder, a sample ID list, an output folder, the RT barcode list, the ligation barcode list and core number and a rendom hexamer barcode list. Then it extract the RT barcode from read1, the ligation barocde from read2, correct them to the nearest RT and ligation barcode (with edit distance <= 1), and attach the RT and ligation barcode and UMI sequence to the read name of read1 and read3. Reads with unmatched RT or ligation barcodes are discarded.

input_folder=$fastq_folder
output_folder=$all_output_folder/UMI_attach
script=$script_path/barcode_extraction.py
echo "Changing the name of the fastq files..."
for sample in $(cat $sample_ID); do echo changing name $sample; mv $input_folder/*$sample*R1*.fastq.gz $input_folder/$sample.R1.fastq.gz; mv $input_folder/*$sample*R2*.fastq.gz $input_folder/$sample.R2.fastq.gz; mv $input_folder/*$sample*R3*.fastq.gz $input_folder/$sample.R3.fastq.gz; done

echo "Attaching barcode and UMI...."
mkdir -p $output_folder
python $script $input_folder $sample_ID $output_folder $ligation_barcode $RT_barcode $core $randomN_barcode_file
echo "Barcode transformed and UMI attached."


################# Trimming the read2
echo
echo "Start trimming the read2 file..."
echo $(date)

trimmed_fastq=$all_output_folder/trimmed_fastq
UMI_attached=$all_output_folder/UMI_attach
bash_script=$script_path/trimming.sh

Rscript $R_script $bash_script $UMI_attached $sample_ID $trimmed_fastq $core


############ Align the reads with STAR, filter the reads based on q > 30, and remove duplicates based on UMI sequence and tagmentation site

#define the output folder for mapping
input_folder=$trimmed_fastq
STAR_output_folder=$all_output_folder/STAR_alignment
filtered_sam_folder=$all_output_folder/filtered_sam
rmdup_sam_folder=$all_output_folder/rmdup_sam

#align read2 to the index file using STAR
echo "Start alignment using STAR..."
echo input folder: $input_folder
echo sample ID file: $sample_ID
echo index file: $index
echo output_folder: $STAR_output_folder
#make the output folder
mkdir -p $STAR_output_folder
#remove the index from the memory
STAR --genomeDir $index --genomeLoad Remove
#start the alignment
for sample in $(cat $sample_ID); do echo Aligning $sample;STAR --runThreadN $core --outSAMstrandField intronMotif --genomeDir $index --readFilesCommand zcat --readFilesIn $input_folder/$sample*R1*gz $input_folder/$sample*R2*gz --outFileNamePrefix $STAR_output_folder/$sample --genomeLoad LoadAndKeep; done
#remove the index from the memory
STAR --genomeDir $index --genomeLoad Remove
echo "All alignment done."


#Filter and sort the sam file
echo
echo "Start filter and sort the sam files..."
echo input folder: $STAR_output_folder
echo output folder: $filtered_sam_folder
bash_script=$script_path/filtering.sh
Rscript $R_script $bash_script $STAR_output_folder $sample_ID $filtered_sam_folder $samtools_core


# Remove the duplicates based on UMI and barcode, chromosome and positiion information.
echo
echo "Start removing duplicates..."
echo input folder: $filtered_sam_folder
echo output folder: $rmdup_sam_folder
mkdir -p $rmdup_sam_folder
bash_script=$script_path/duplicate_removal.sh
Rscript $R_script $bash_script $filtered_sam_folder $sample_ID $rmdup_sam_folder $core $script_path


################# Split the sam file based on the barcode
sam_folder=$all_output_folder/rmdup_sam
output_folder=$all_output_folder/sam_splitted

echo
echo "Start splitting the sam file..."
echo samfile folder: $sam_folder
echo sample list: $sample_ID
echo ouput folder: $output_folder
echo cutoff value: $cutoff

bash_script=$script_path/sam_split.sh
Rscript $R_script $bash_script $sam_folder $sample_ID $output_folder $core $cutoff $script_path


cat $output_folder/*sample_list.txt>$output_folder/All_samples.txt
cp $output_folder/All_samples.txt $output_folder/../barcode_samples.txt
# output the report the report/barcode_read_distribution folder
mkdir -p $output_folder/../report/barcode_read_distribution
mv $output_folder/*.txt $output_folder/../report/barcode_read_distribution/
echo
echo "All sam file splitted."


################### Calculate the reads numbers along the computational pipeline
fastq_folder=$fastq_folder
trimmed_folder=$trimmed_fastq
UMI_attach=$all_output_folder/UMI_attach
alignment=$STAR_output_folder
filtered_sam=$filtered_sam_folder
rm_dup_sam=$rmdup_sam_folder
#split_sam=$parental_folder/splited_sam
report_folder=$all_output_folder/report/read_num
echo
echo "Start calculating the reads number..."
#make the report folder
mkdir -p $report_folder
#calculate the read number and output the read number into the report folder
echo sample,total reads,after filtering barcode,after trimming,uniquely aligned reads,After remove duplicates>$report_folder/read_number.csv
for sample in $(cat $sample_ID); do echo calculating $sample; echo $sample,$(expr $(zcat $fastq_folder/$sample*R2*.gz|wc -l) / 4),$(expr $(zcat $UMI_attach/$sample*R2*.gz|wc -l) / 4),$(expr $(zcat $trimmed_folder/$sample*R2*.gz|wc -l) / 4) ,$(expr $(samtools view $filtered_sam/$sample.sam|wc -l) / 2),$(expr $(samtools view $rm_dup_sam/$sample.sam|wc -l) / 2)>>$report_folder/read_number.csv; done
echo "Read number calculation is done."


##################### Count reads mapping to genes
output_folder=$all_output_folder/report/human_mouse_gene_count/
input_folder=$all_output_folder/sam_splitted
script=$script_path/gene_counting.py
sample_ID=$all_output_folder/barcode_samples.txt
echo "Start the gene count...."
python $script $gtf_file $input_folder $sample_ID $core $randomN_barcode_file

echo "Make the output folder and transfer the files..."
mkdir -p $output_folder
find $input_folder/ -name "*.count" -print0 | sort -z | xargs -r0 cat > $output_folder/count.MM
find $input_folder/ -name "*.count" -delete
find $input_folder/ -name "*.report" -print0 | sort -z | xargs -r0 cat > $output_folder/report.MM
find $input_folder/ -name "*.report" -delete
mv $input_folder/*_annotate.txt $output_folder/
echo "All output files are transferred~"

################# Transform output files to an easy to read in R file
conda deactivate
conda activate original_pipeline_final_step
R_script=$script_path/formatting.R
Rscript $R_script $all_output_folder/report
conda deactivate

mkdir $all_output_folder/report/Log_files
mv Log.out $all_output_folder/report/Log_files/
mv Log.progress.out $all_output_folder/report/Log_files/
mv Aligned.out.sam $all_output_folder/report/Log_files/
mv _STARtmp $all_output_folder/report/Log_files/


################# exon counting
conda activate original_pipeline
output_folder=$all_output_folder/report/exon-level/
input_folder=$all_output_folder/sam_splitted/
script=$script_path/exon_counting.py
sample_ID=$all_output_folder/barcode_samples.txt
echo "Start the exon count...."
python $script $gtf_file_exon $input_folder $sample_ID $core

echo "Make the output folder and transfer the files..."
mkdir -p $output_folder
find $input_folder/ -name "*.count" -print0 | sort -z | xargs -r0 cat > $output_folder/count.MM
find $input_folder/ -name "*.count" -delete
find $input_folder/ -name "*.report" -print0 | sort -z | xargs -r0 cat > $output_folder/report.MM
find $input_folder/ -name "*.report" -delete
mv $input_folder/*_annotate.txt $output_folder/
echo "All output files are transferred after exon counting~"

################# transform output files to an easy to read in R file
conda deactivate
conda activate original_pipeline_final_step
R_script=$script_path/formatting_exon.R
Rscript $R_script $all_output_folder/report/exon-level/
conda deactivate


now=$(date)
echo "Current time : $now"
```
