## Supplementary file 3 for "A global view of aging and Alzheimer’s pathogenesis-associated cell population dynamics and molecular signatures in the human and mouse brains": README.txt

EasySci-RNA computational processing pipeline:

This pipeline takes the raw sequencer generated files as input and outputs the cell/gene matrix. The pipeline consist of the following steps: demultiplexing based on the P7 barcodes; barcode extraction and matching; trimming the adaptor and polyA sequences, alignment; filtyering of the low-quality alignments; PCR duplicate removal; splitting of the reads to single-cell SAM files and removal of low signal cells; gene and exon expression counting.


Demultiplexing:

Demultiplexing for EasySci-RNA uses only the P7 barcodes, because in EasySci-RNA the P5 barcode is added by ligation and every possible P5-P7 combination is present in the final data. This would result in a very high file number if both barcodes would be used to demultiplex. The demultiplexing is done with Illumina’s bcl2fastq software with the following settings:

bcl2fastq --runfolder-dir INPUT_FOLDER(sequencer generated files) -o OUTPUT_FOLDER --sample-sheet SAMPLE_SHEET(example can be found in the “example_files” folder) --reports-dir OUTPUT_FOLDER/report --barcode-mismatches 1 --create-fastq-for-index-reads --no-lane-splitting --use-bases-mask Y*,I*,Y*,Y* --minimum-trimmed-read-length 0 --mask-short-adapter-reads 0

After demultiplexing there should be 4 files:
•	R1: Read1
•	R2: P5 barcode
•	R3: read2
•	I1: P7 barcode


Main computational pipeline:

Run the EasySci_main.sh file to use the computational pipeline:
The following input parameters need to be set:

•	bashrc_location: Location of the bashrc file to activate the conda environments.

•	fastq_folder: Input FASTQ files folder.

•	sample_ID: Sample ID file, name of the demultiplexed files, one sample name per row without the R1/R1/R3 or fastq ending. An example file can be found in the “example_files” folder.

•	all_output_folder: Output folder for the intermediary and final files.

•	core: Number of cores used during the computational pipeline.

•	samtools_core: Number of cores used during the filtering and sorting step.

•	cutoff: Number of read cutoff for splitting single cell; cells with reads less than this number will be discarded. Since this pipeline uses both read pairs, the actual number of read pairs kept will be half this number.

•	index: STAR index file for alignment. This file was generated by the following STAR command:

	STAR --runThreadN 15 --runMode genomeGenerate --genomeDir OUTPUT_FOLDER --genomeFastaFiles INPUT_FASTA_FILE --sjdbGTFfile INPUT_GTF_FILE

	INPUT_FASTA_FILE was downloaded from this location:
	http://ftp.ebi.ac.uk/pub/databases/gencode/Gencode_mouse/release_M27/GRCm39.primary_assembly.genome.fa.gz
	INPUT_GTF_FILE was downloaded from this location: http://ftp.ebi.ac.uk/pub/databases/gencode/Gencode_mouse/release_M27/gencode.vM27.primary_assembly.annotation.gtf.gz

•	gtf_file: GTF file for the gene counting step, this file can be downloaded from the following location: http://ftp.ebi.ac.uk/pub/databases/gencode/Gencode_mouse/release_M27/gencode.vM27.primary_assembly.annotation.gtf.gz

•	gtf_file_exon: GTF file for the exon counting step, this file can be created from the above GTF file by subsetting for only the exonic regions.

•	ligation_barcode: Ligation barcodes as a dictionary, where barcodes one edit distance away are included. This file can be created using the generate_barcode_dictionary.sh script in the script_folder, but the file with the barcode list used in our experiments are included for easier usage.

•	RT_barcode: Reverse transcription barcodes as a dictionary, where barcodes one edit distance away are included. This file can be created using the generate_barcode_dictionary.sh script in the script_folder, but the file with the barcode list used in our experiments are included for easier usage.

•	randomN_barcode_file: Random hexamer barcode list as a text file.


Results:

The final output file can be found here: output_folder/report/Summary.RData . This file can be loaded in R using the “load()” command and will contain 5 data frames:

•	df_cell: cell annotation
•	df_gene: gene annotation
•	gene_count_all: gene x cell expression matrix for reads overlapping with exons+introns (this file is used usually for downstream analysis)
•	gene_count_exon: gene x cell expression matrix for only reads mapping to exons
•	gene_count_intron: gene x cell expression matrix for only reads mapping to introns

The exon-level count matrix can be found here: output_folder/report/exon-level/Summary.RData. This file contains the following data frames:

•	df_cell: cell annotation
•	df_gene: exon annotation
•	gene_count_all: exon x cell expression matrix

During the computational pipeline the reads originating from the shortdT and random hexamer RT primers are kept as separate pseudo cells. During the downstream processing the counts from the same cells but from the different RT primers need to be merged, based on matching ligation, PCR barcodes and the paired RT primers per well.


Environments:

During the computational pipeline 3 separate conda environments were used. One during the demultiplexing step and two during the main computational pipeline (environments original_pipeline and original_pipeline_final_step). Below are the detailed lists of the softwares installed in these environments.


Demultiplexing environment:

# Name                    Version                   Build  Channel
_libgcc_mutex             0.1                        main  
bcl2fastq                 2.19.0                        1    dranew
boost_lib                 1.54.0                        1    dranew
libgcc                    7.2.0                h69d50b8_2  
libgcc-ng                 9.1.0                hdf63c60_0  
libstdcxx-ng              9.1.0                hdf63c60_0  
zlib                      1.2.11               h7b6447c_3  


Environment original_pipeline:

# Name                    Version                   Build  Channel
_r-mutex                  1.0.0               anacondar_1  
backports                 1.0                        py_2    anaconda
backports.functools_lru_cache 1.6.1                      py_0    anaconda
backports.shutil_get_terminal_size 1.0.0                    py27_2    anaconda
backports_abc             0.5                        py_1    anaconda
bioconductor-biocparallel 1.4.3                  r3.2.2_0    bioconda
biopython                 1.74             py27h7b6447c_0    anaconda
blas                      1.0                         mkl    anaconda
bzip2                     1.0.8                h7b6447c_0  
ca-certificates           2020.10.14                    0    anaconda
cairo                     1.14.12              h8948797_3  
certifi                   2019.11.28               py27_0    anaconda
curl                      7.55.1               hcb0b314_2    anaconda
cutadapt                  1.8.3                    py27_0    bioconda
cycler                    0.10.0                   py27_0    anaconda
dbus                      1.13.16              hb2f20db_0    anaconda
decorator                 4.4.2                      py_0    anaconda
enum34                    1.1.6                    py27_1    anaconda
expat                     2.2.9                he6710b0_2    anaconda
fastqc                    0.11.9                        0    bioconda
font-ttf-dejavu-sans-mono 2.37                 h6964260_0  
fontconfig                2.13.0               h9420a91_0  
freetype                  2.10.2               h5ab3b9f_0  
fribidi                   1.0.10               h7b6447c_0  
functools32               3.2.3.2                  py27_1    anaconda
futures                   3.3.0                    py27_0    anaconda
glib                      2.65.0               h3eb4bd4_0  
graphite2                 1.3.14               h23475e2_0  
gst-plugins-base          1.14.0               hbbd80ab_1  
gstreamer                 1.14.0               hb31296c_0  
harfbuzz                  2.4.0                hca77d97_1  
htseq                     0.11.3           py27hb3f55d8_0    bioconda
icu                       58.2                 he6710b0_3  
intel-openmp              2020.2                      254    anaconda
ipykernel                 4.10.0                   py27_0    anaconda
ipython                   5.8.0                    py27_0    anaconda
ipython_genutils          0.2.0                    py27_0    anaconda
jpeg                      9b                   h024ee3a_2  
jupyter_client            5.3.4                    py27_0    anaconda
jupyter_core              4.6.1                    py27_0    anaconda
kiwisolver                1.1.0            py27he6710b0_0    anaconda
libdeflate                1.0                  h14c3975_1    bioconda
libedit                   3.1.20191231         h14c3975_1    anaconda
libffi                    3.3                  he6710b0_2    anaconda
libgcc                    7.2.0                h69d50b8_2  
libgcc-ng                 9.1.0                hdf63c60_0    anaconda
libgfortran-ng            7.3.0                hdf63c60_0    anaconda
libpng                    1.6.37               hbc83047_0  
libsodium                 1.0.18               h7b6447c_0    anaconda
libssh2                   1.8.0                h9cfc8f7_4    anaconda
libstdcxx-ng              9.1.0                hdf63c60_0    anaconda
libtiff                   4.1.0                h2733197_1  
libuuid                   1.0.3                h1bed415_2  
libxcb                    1.14                 h7b6447c_0  
libxml2                   2.9.10               he19cac6_1  
lz4-c                     1.9.2                he6710b0_1  
matplotlib                2.2.3            py27hb69df0a_0    anaconda
mkl                       2019.4                      243    anaconda
mkl-service               2.3.0            py27he904b0f_0    anaconda
mkl_fft                   1.0.15           py27ha843d7b_0    anaconda
mkl_random                1.1.0            py27hd6b4f25_0    anaconda
ncurses                   6.2                  he6710b0_1    anaconda
numpy                     1.16.6           py27hbc911f0_0    anaconda
numpy-base                1.16.6           py27hde5b4d6_0    anaconda
openjdk                   8.0.152              h7b6447c_3  
openssl                   1.0.2u               h7b6447c_0    anaconda
pandas                    0.22.0           py27hf484d3e_0    anaconda
pango                     1.45.3               hd140c19_0  
pathlib2                  2.3.5                    py27_0    anaconda
pcre                      8.44                 he6710b0_0  
perl                      5.26.2               h14c3975_0  
pexpect                   4.7.0                    py27_0    anaconda
pickleshare               0.7.5                    py27_0    anaconda
pip                       19.3.1                   py27_0    anaconda
pixman                    0.40.0               h7b6447c_0  
prompt_toolkit            1.0.15                   py27_0    anaconda
ptyprocess                0.6.0                    py27_0    anaconda
pygments                  2.5.2                      py_0    anaconda
pyparsing                 2.4.7                      py_0    anaconda
pyqt                      5.9.2            py27h22d08a2_1    anaconda
pysam                     0.15.3           py27hda2845c_1    bioconda
python                    2.7.18               h15b4118_1    anaconda
python-dateutil           2.8.1                      py_0    anaconda
python-levenshtein        0.12.0          py27h516909a_1001    conda-forge
pytz                      2020.1                     py_0    anaconda
pyzmq                     18.1.0           py27he6710b0_0    anaconda
qt                        5.9.6                h8703b6f_2  
r                         3.2.2                         0  
r-base                    3.2.2                         0  
r-boot                    1.3_17                r3.2.2_0a  
r-class                   7.3_14                r3.2.2_0a  
r-cluster                 2.0.3                 r3.2.2_0a  
r-codetools               0.2_14                r3.2.2_0a  
r-foreign                 0.8_66                r3.2.2_0a  
r-futile.logger           1.4.1                  r3.2.2_0    bioconda
r-futile.options          1.0.0                  r3.2.2_0    bioconda
r-kernsmooth              2.23_15               r3.2.2_0a  
r-lambda.r                1.1.7                  r3.2.2_0    bioconda
r-lattice                 0.20_33               r3.2.2_0a  
r-mass                    7.3_45                r3.2.2_0a  
r-matrix                  1.2_2                 r3.2.2_0a  
r-mgcv                    1.8_9                 r3.2.2_0a  
r-nlme                    3.1_122               r3.2.2_0a  
r-nnet                    7.3_11                r3.2.2_0a  
r-recommended             3.2.2                  r3.2.2_0  
r-rpart                   4.1_10                r3.2.2_0a  
r-snow                    0.4_1                  r3.2.2_0    bioconda
r-spatial                 7.3_11                r3.2.2_0a  
r-survival                2.38_3                r3.2.2_0a  
readline                  8.0                  h7b6447c_0    anaconda
samtools                  1.4.1                         0    bioconda
scandir                   1.10.0           py27h7b6447c_0    anaconda
setuptools                44.0.0                   py27_0    anaconda
simplegeneric             0.8.1                    py27_2    anaconda
singledispatch            3.4.0.3                  py27_0    anaconda
sip                       4.19.13          py27he6710b0_0    anaconda
six                       1.15.0                     py_0    anaconda
sqlite                    3.33.0               h62c20be_0    anaconda
star                      2.5.2b                        0    bioconda
subprocess32              3.5.4            py27h7b6447c_0    anaconda
tk                        8.6.10               hbc83047_0    anaconda
tornado                   5.1.1            py27h7b6447c_0    anaconda
traitlets                 4.3.3                    py27_0    anaconda
trim-galore               0.4.1                         1    bioconda
wcwidth                   0.2.5                      py_0    anaconda
wheel                     0.35.1                     py_0    anaconda
xz                        5.2.5                h7b6447c_0  
zeromq                    4.3.3                he6710b0_3    anaconda
zlib                      1.2.11               h7b6447c_3    anaconda
zstd                      1.4.5                h9ceee32_0  


Environment original_pipeline_final_step:

# Name                    Version                   Build  Channel
_libgcc_mutex             0.1                        main  
_r-mutex                  1.0.0                     mro_2    r
binutils_impl_linux-64    2.33.1               he6710b0_7  
binutils_linux-64         2.33.1              h9595d00_15  
ca-certificates           2020.7.22                     0  
cairo                     1.14.12              h8948797_3  
curl                      7.71.1               hbc83047_1  
fontconfig                2.13.0               h9420a91_0  
freetype                  2.10.2               h5ab3b9f_0  
fribidi                   1.0.10               h7b6447c_0  
gcc_impl_linux-64         7.3.0                habb00fd_1  
gcc_linux-64              7.3.0               h553295d_15  
gfortran_impl_linux-64    7.3.0                hdf63c60_1  
gfortran_linux-64         7.3.0               h553295d_15  
glib                      2.56.2               hd408876_0  
graphite2                 1.3.14               h23475e2_0  
gxx_impl_linux-64         7.3.0                hdf63c60_1  
gxx_linux-64              7.3.0               h553295d_15  
harfbuzz                  1.8.8                hffaf4a1_0  
icu                       58.2                 he6710b0_3  
krb5                      1.18.2               h173b8e3_0  
ld_impl_linux-64          2.33.1               h53a641e_7  
libcurl                   7.71.1               h20c2e04_1  
libedit                   3.1.20191231         h14c3975_1  
libffi                    3.2.1             hf484d3e_1007  
libgcc-ng                 9.1.0                hdf63c60_0  
libgfortran-ng            7.3.0                hdf63c60_0  
libpng                    1.6.37               hbc83047_0  
libssh2                   1.9.0                h1ba5d50_1  
libstdcxx-ng              9.1.0                hdf63c60_0  
libuuid                   1.0.3                h1bed415_2  
libxcb                    1.14                 h7b6447c_0  
libxml2                   2.9.10               he19cac6_1  
make                      4.2.1                h1bed415_1  
mro-base                  3.5.1                         3    r
mro-base_impl             3.5.1                h9a62091_0    r
ncurses                   6.2                  he6710b0_1  
openssl                   1.1.1h               h7b6447c_0  
pango                     1.42.4               h049681c_0  
pcre                      8.44                 he6710b0_0  
pixman                    0.40.0               h7b6447c_0  
r-assertthat              0.2.0           mro351hf348343_0    r
r-backports               1.1.2           mro351hd10c6a6_0    r
r-base64enc               0.1_3           mro351hd10c6a6_0    r
r-bh                      1.66.0_1        mro351hf348343_0    r
r-bindr                   0.1.1           mro351hf348343_0    r
r-bindrcpp                0.2.2           mro351hebc1506_0    r
r-broom                   0.5.0           mro351hf348343_0    r
r-callr                   2.0.4           mro351hf348343_0    r
r-cellranger              1.1.0           mro351hf348343_0    r
r-cli                     1.0.0           mro351hf348343_0    r
r-clipr                   0.4.1           mro351hf348343_0    r
r-colorspace              1.3_2           mro351hd10c6a6_0    r
r-crayon                  1.3.4           mro351hf348343_0    r
r-curl                    3.2             mro351hd10c6a6_1    r
r-data.table              1.11.4          mro351hd10c6a6_0    r
r-dbi                     1.0.0           mro351hf348343_0    r
r-dbplyr                  1.2.2           mro351hf348343_0    r
r-dichromat               2.0_0           mro351hf348343_0    r
r-digest                  0.6.15          mro351hd10c6a6_0    r
r-dplyr                   0.7.6           mro351hebc1506_0    r
r-evaluate                0.11            mro351hf348343_0    r
r-fansi                   0.2.3           mro351hd10c6a6_0    r
r-forcats                 0.3.0           mro351hf348343_0    r
r-ggplot2                 2.2.1           mro343h889e2dd_0    r
r-glue                    1.3.0           mro351hd10c6a6_0    r
r-gtable                  0.2.0           mro351hf348343_0    r
r-haven                   1.1.2           mro351hebc1506_0    r
r-highr                   0.7             mro351hf348343_0    r
r-hms                     0.4.2           mro351hf348343_0    r
r-htmltools               0.3.6           mro351hebc1506_0    r
r-httr                    1.3.1           mro351hf348343_1    r
r-jsonlite                1.5             mro351hd10c6a6_0    r
r-knitr                   1.20            mro351hf348343_0    r
r-labeling                0.3             mro351hf348343_0    r
r-lattice                 0.20_35         mro351hd10c6a6_0    r
r-lazyeval                0.2.1           mro351hd10c6a6_0    r
r-lubridate               1.7.4           mro351hebc1506_0    r
r-magrittr                1.5             mro351hf348343_0    r
r-markdown                0.8             mro351hd10c6a6_0    r
r-mass                    7.3_50          mro351hd10c6a6_0    r
r-matrix                  1.2_14          mro351hac1494b_0    r
r-mime                    0.5             mro351hd10c6a6_0    r
r-modelr                  0.1.2           mro351hf348343_0    r
r-munsell                 0.5.0           mro351hf348343_0    r
r-nlme                    3.1_137         mro351hac1494b_0    r
r-openssl                 1.0.2           mro351hd10c6a6_1    r
r-pillar                  1.3.0           mro351hf348343_0    r
r-pkgconfig               2.0.1           mro351hf348343_0    r
r-plogr                   0.2.0           mro351hf348343_0    r
r-plyr                    1.8.4           mro351hebc1506_0    r
r-praise                  1.0.0           mro351hf348343_0    r
r-processx                3.1.0           mro351hebc1506_0    r
r-purrr                   0.2.5           mro351hd10c6a6_0    r
r-r6                      2.2.2           mro351hf348343_0    r
r-rcolorbrewer            1.1_2           mro351hf348343_0    r
r-rcpp                    0.12.18         mro351hebc1506_0    r
r-readr                   1.1.1           mro351hebc1506_0    r
r-readxl                  1.1.0           mro351hebc1506_0    r
r-rematch                 1.0.1           mro351hf348343_0    r
r-reprex                  0.2.0           mro351hf348343_0    r
r-reshape2                1.4.3           mro351hebc1506_0    r
r-revoutils               11.0.0                 mro351_0    r
r-revoutilsmath           11.0.0                 mro351_0    r
r-rlang                   0.2.1           mro351hd10c6a6_0    r
r-rmarkdown               1.10            mro351hf348343_0    r
r-rprojroot               1.3_2           mro351hf348343_0    r
r-rstudioapi              0.7             mro351hf348343_0    r
r-rvest                   0.3.2           mro351hf348343_0    r
r-scales                  0.5.0           mro351hebc1506_0    r
r-selectr                 0.4_1           mro351hf348343_0    r
r-stringi                 1.2.4           mro351hebc1506_0    r
r-stringr                 1.3.1           mro351hf348343_0    r
r-testthat                2.0.0           mro351hebc1506_0    r
r-tibble                  1.4.2           mro351hd10c6a6_0    r
r-tidyr                   0.8.1           mro351hebc1506_0    r
r-tidyselect              0.2.4           mro351hebc1506_0    r
r-tidyverse               1.2.1           mro351hf348343_0    r
r-tinytex                 0.6             mro351hf348343_0    r
r-utf8                    1.1.4           mro351hd10c6a6_0    r
r-viridislite             0.3.0           mro351hf348343_0    r
r-whisker                 0.3_2           mro351hf348343_0    r
r-withr                   2.1.2           mro351hf348343_0    r
r-xfun                    0.3             mro351hf348343_0    r
r-xml2                    1.2.0           mro351hebc1506_0    r
r-yaml                    2.2.0           mro351hd10c6a6_0    r
xz                        5.2.5                h7b6447c_0  
zlib                      1.2.11               h7b6447c_3  
