## Supplementary file 3 for "A global view of aging and Alzheimer’s pathogenesis-associated cell population dynamics and molecular signatures in the human and mouse brains": barcode_extraction.py

```
import subprocess
import sys
from Levenshtein import distance
import gzip
from multiprocessing import Pool
from functools import partial
import pickle
import re
from Bio.Seq import Seq

    
def UMI_attach_read2_barcode_list(sample, input_folder, output_folder, ligation_barcode_list, RT_barcode_list, randomN_barcodes):
    #open the read1, read2, and output file
    Read1 = input_folder + "/" + sample + ".R1.fastq.gz"
    Read2 = input_folder + "/" + sample + ".R3.fastq.gz"
    Read3 = input_folder + "/" + sample + ".R2.fastq.gz"
    output_file1 = output_folder + "/" + sample + ".R2.fastq.gz"
    output_file2 = output_folder + "/" + sample + ".R1.fastq.gz"
    f1 = gzip.open(Read1)
    f2 = gzip.open(Read2)
    f3 = gzip.open(output_file1, 'wb')
    f4 = gzip.open(Read3)
    f5 = gzip.open(output_file2, 'wb')
    
    line1 = f1.readline()
    line2 = f2.readline()
    line3 = f4.readline()
    total_line = 0
    filtered_line = 0
    
    while (line1):
        total_line += 1
        line1 = f1.readline()
        line3 = f4.readline()
        # first check if the ligation barcode match with the barcode
        tmp_lig = line3[0:10]

        #print("ligation barcode: ", ligation_bc_match)
        if tmp_lig in ligation_barcode_list:
            
            ligation_bc_match = ligation_barcode_list[tmp_lig]
            # check RT barcode
            target_RT = line1[8:18]

            if target_RT in RT_barcode_list:
                barcode = RT_barcode_list[target_RT]
                UMI = line1[:8]


                if barcode in randomN_barcodes:
                    first_line_3 = '@' + ligation_bc_match + barcode + ',' + UMI + ',' + re.sub("3:N:","2:N:",line2[1:])
                    first_line_5 = '@' + ligation_bc_match + barcode + ',' + UMI + ',' + re.sub("3:N:","1:N:",line2[1:])

                    second_line_5 = line1[18:]                
                    result_adaptor = re.search("CTGTCTCTTATACACAT", second_line_5)
                    if result_adaptor == None:
                        second_line_5 = second_line_5
                    else:
                        second_line_5 = second_line_5[:result_adaptor.start()] + "\n"

                    seq = Seq(target_RT)
                    seqRC = str(seq.reverse_complement())
                    second_line_3 = f2.readline()
                    result_barcode = re.search(seqRC, second_line_3)
                    if result_barcode == None:
                        second_line_3 = second_line_3
                    else:
                        second_line_3 = second_line_3[:result_barcode.start()] + "\n"

                    third_line_3 = f2.readline()
                    third_line_5 = f1.readline()

                    four_line_5 = f1.readline()
                    four_line_5 = four_line_5[18:]
                    if result_adaptor == None:
                        four_line_5 = four_line_5
                    else:
                        four_line_5 = four_line_5[:result_adaptor.start()] + "\n"

                    four_line_3 = f2.readline()
                    if result_barcode == None:
                        four_line_3 = four_line_3
                    else:
       	       	       	four_line_3 = four_line_3[:result_barcode.start()] + "\n"

                    line1 = f1.readline()
                    line2 = f2.readline()

                    if len(second_line_3) > 20 and len(second_line_5) > 20 :
                        filtered_line += 1
                        f3.write(first_line_3)
                        f5.write(first_line_5)
       	       	       	f3.write(second_line_3)
       	       	       	f5.write(second_line_5)
       	       	       	f3.write(third_line_3)
       	       	       	f5.write(third_line_5)
       	       	       	f3.write(four_line_3)
       	       	       	f5.write(four_line_5) 

                else:            

                    first_line_3 = '@' + ligation_bc_match + barcode + ',' + UMI + ',' + re.sub("3:N:","2:N:",line2[1:])
                    first_line_5 = '@' + ligation_bc_match + barcode + ',' + UMI + ',' + re.sub("3:N:","1:N:",line2[1:])

                    second_line_5 = line1[33:]
                    result_adaptor = re.search("CTGTCTCTTATACACAT", second_line_5)
                    if result_adaptor == None:
                        second_line_5 = second_line_5
                    else:
                        second_line_5 = second_line_5[:result_adaptor.start()] + "\n"

                    seq = Seq(target_RT)
                    seqRC = str(seq.reverse_complement())
                    second_line_3 = f2.readline()
                    result_barcode = re.search(seqRC, second_line_3)
                    if result_barcode == None:
                        second_line_3 = second_line_3
                    else:
                        second_line_3 = second_line_3[:(result_barcode.start() - 15)] + "\n"

                    third_line_3 = f2.readline()
                    third_line_5 = f1.readline()

                    four_line_5 = f1.readline()
                    four_line_5 = four_line_5[33:]
                    if result_adaptor == None:
                        four_line_5 = four_line_5
                    else:
                        four_line_5 = four_line_5[:result_adaptor.start()] + "\n"

                    four_line_3 = f2.readline()
                    if result_barcode == None:
                        four_line_3 = four_line_3
                    else:
                        four_line_3 = four_line_3[:(result_barcode.start() - 15)] + "\n"

                    line1 = f1.readline()
                    line2 = f2.readline()

                    if len(second_line_3) > 20 and len(second_line_5) > 20 :
                        filtered_line += 1
                        f3.write(first_line_3)
                        f5.write(first_line_5)
                        f3.write(second_line_3)
                        f5.write(second_line_5)
                        f3.write(third_line_3)
                        f5.write(third_line_5)
                        f3.write(four_line_3)
                        f5.write(four_line_5)


            else:
                line2 = f2.readline()
                line2 = f2.readline()
                line2 = f2.readline()
                line2 = f2.readline()
                line1 = f1.readline()
                line1 = f1.readline()
                line1 = f1.readline()

                
        else:
            line2 = f2.readline()
            line2 = f2.readline()
            line2 = f2.readline()
            line2 = f2.readline()
       	    line1 = f1.readline()
            line1 = f1.readline()
            line1 = f1.readline()


        line3 = f4.readline() 
        line3 = f4.readline()
        line3 = f4.readline()

    f1.close()
    f2.close()
    f3.close()
    f4.close()
    f5.close()
    print("sample name: %s, total line: %f, filtered line: %f, filter rate: %f" 
          %(sample, total_line, filtered_line, float(filtered_line) / float(total_line)))

# this function accept an input folder and a output folder and then generate the output file with the index
def attach_UMI_files(input_folder, sampleID, output_folder, ligation_barcode_file, RT_barcode_file, core, randomN_barcode_file):
    
    init_message = '''
    --------------------------start attaching UMI-----------------------------
    input folder: %s
    sample ID: %s
    output_folder: %s
    ligation barcode file: %s
    RT barcode file: %s
    ___________________________________________________________________________
    ''' %(input_folder, sampleID, output_folder, ligation_barcode_file, RT_barcode_file)
    
    print(init_message)
    
    print("Load ligation barcode dictionary...")
    
    # generate the ligation barcode list
    barcodes = open(ligation_barcode_file, "rb")
    ligation_barcode_list = pickle.load(barcodes)
    barcodes.close()
    
    print("Load RT barcode dictionary...")
    
    # generate the RT barcode list:
    barcodes = open(RT_barcode_file, "rb")
    RT_barcode_list = pickle.load(barcodes)
    barcodes.close()

    # load randomN barcodes
    barcodes = open(randomN_barcode_file, "rb")
    randomN_barcodes = []
    for line in barcodes:
        barcode = line.strip()
        randomN_barcodes.append(barcode)
    barcodes.close()   
    
    #for each sample in the sample list, use the read1 file, read2 file, output file
    # and barcode_list to run UMI_attach_read2_barcode_list
    sample_file = open(sampleID)
    sample_list = []
    for line in sample_file:
        sample = line.strip()
        sample_list.append(sample)
    sample_file.close()
    
    # parallele for the functions
    p = Pool(processes = int(core))
    func = partial(UMI_attach_read2_barcode_list, input_folder = input_folder, output_folder=output_folder, ligation_barcode_list = ligation_barcode_list, RT_barcode_list=RT_barcode_list, randomN_barcodes = randomN_barcodes)
    result = p.map(func, sample_list)
    p.close()
    p.join()
    
    #print the completion message
    com_message = '''~~~~~~~~~~~~~~~UMI attachment done~~~~~~~~~~~~~~~~~~'''
    print(com_message)
    
if __name__ == "__main__":
    input_folder = sys.argv[1]
    sampleID = sys.argv[2]
    output_folder = sys.argv[3]
    ligation_barcode_file = sys.argv[4]
    RT_barcode_file = sys.argv[5]
    core=sys.argv[6]
    randomN_barcode_file=sys.argv[7]
    attach_UMI_files(input_folder, sampleID, output_folder, ligation_barcode_file, RT_barcode_file, core, randomN_barcode_file)
```
