## Supplementary file 3 for "A global view of aging and Alzheimer’s pathogenesis-associated cell population dynamics and molecular signatures in the human and mouse brains": duplicate_removal.py

```
from Levenshtein import distance
import sys
import matplotlib as mpl
import numpy as np
mpl.use('Agg')
import matplotlib.pyplot as plt
import pandas as pd
sys.setrecursionlimit(1000000)
def rm_dup_samfile(samfile, output_file):
    f1 = open(samfile)
    f2 = open(output_file, 'w')
    f3 = open(output_file+'.csv', 'w')
    pre_location = []
    pre_barcode_UMI = [] 

    dup = False
    
    pre_dup_num = 0
    cur_dup_num = 0

    line1 = f1.readline()
    
    while(line1):
        
        if (line1[0] == '@'):
            f2.write(line1)
            line1 = f1.readline()
        else:
            name1 = (((line1.split('\t'))[0]).split(','))
            barcode_UMI1 = name1[0] + name1[1]
            chrom_num1 = (line1.split('\t'))[2]
            start_site1 = (line1.split('\t'))[3]
            
            line2 = f1.readline()
            chrom_num2 = (line2.split('\t'))[2]
            start_site2 = (line2.split('\t'))[3]
            current_location = chrom_num1 + "-" + start_site1 + "-" + chrom_num2 + "-" + start_site2

            if((line1.split('\t'))[0] != (line2.split('\t'))[0]):
                print "WARNING: not proper read pairing during duplicate removal"

            if ((barcode_UMI1 == pre_barcode_UMI)):    
                dup = False
                for each_location in pre_location:
                    if each_location == current_location:
                        dup = True
                        break
                if dup == False:
                    pre_dup_num = cur_dup_num
                    cur_dup_num = 1
                    f2.write(line1)
                    f2.write(line2)
                    pre_location.add(current_location)
                    f3.write('%d' %(pre_dup_num))
                    f3.write('\n')
                    line1 = f1.readline()
                else:
                    cur_dup_num += 1
                    line1 = f1.readline()              
            else:
                pre_dup_num = cur_dup_num
                cur_dup_num = 1
                f2.write(line1)
                f2.write(line2)
                pre_barcode_UMI = barcode_UMI1
                pre_location = set()
                pre_location.add(current_location)
                line1 = f1.readline()
                if (pre_dup_num != 0):
                    f3.write("%d" % (pre_dup_num))
                    f3.write('\n')
    
    f1.close()
    f2.close()
    f3.close()
    
    '''
    #plot the histogram for the read duplication number
    dups = (pd.read_csv(output_file+'.csv', header=None))[0]
    fig = plt.figure()
    plt.hist(dups, bins=100)
    plt.xlabel("Duplication number")
    plt.ylabel("Read number")
    fig.savefig(output_file + '.png')
    '''

if __name__ == "__main__":
    samfile = sys.argv[1]
    output_file = sys.argv[2]
    rm_dup_samfile(samfile, output_file)
```
