## Supplementary file 3 for "A global view of aging and Alzheimer’s pathogenesis-associated cell population dynamics and molecular signatures in the human and mouse brains": duplicate_removal.sh

```
input_folder=$1
sample=$2
output_folder=$3
script_path_folder=$4

python_script=$script_path_folder/duplicate_removal.py
echo Filtering sample: $sample
python $python_script $input_folder/$sample.sam $output_folder/$sample.sam
echo Filtering $sample done.
```
