## Supplementary file 3 for "A global view of aging and Alzheimer’s pathogenesis-associated cell population dynamics and molecular signatures in the human and mouse brains": exon_counting.py

```
from __future__ import division
import itertools
import collections
import numpy as np
import pandas as pd
from multiprocessing import Pool
from multiprocessing import *
import HTSeq
import sys
from functools import partial
import logging

def sciRNAseq_count(sample, input_folder, genes, gene_annotat, sample_ID):
    input_sam = input_folder + "/" + sample + ".sam"
    report = input_folder + "/" + sample + ".report"
    count_output = input_folder + "/" + sample + ".count"
    
    counts = collections.Counter()
    sam_file = input_sam
    almnt_file = HTSeq.SAM_Reader(sam_file)
    sam_name = sample
    cell_ID = sample_ID.index(sample) + 1

    perfect_inter_exon = 0
    
    print("Start read the input file: " + sam_file + "....")

    for bundle in HTSeq.pair_SAM_alignments( almnt_file, bundle=True ):

        if len(bundle) != 1:
            continue  # Skip multiple alignments
        first_almnt, second_almnt = bundle[0]  # extract pair
        if (not first_almnt.aligned) or (not second_almnt.aligned):
            count[ "_unmapped" ] += 1
            continue
        if (first_almnt.iv.chrom not in genes.chrom_vectors) or (second_almnt.iv.chrom not in genes.chrom_vectors):
            counts["_unmapped"] += 1
            continue

        if first_almnt.iv.strand == "+":
            first_almnt.iv.strand = "-"
        else:
            first_almnt.iv.strand = "+"        

        gene_id_intersect_1 = set()
        inter_count_1 = 0
        for cigop in first_almnt.cigar:
            if cigop.type != "M":
                continue
            if cigop.ref_iv.strand == "+":
                cigop.ref_iv.strand = "-"
            else:
                cigop.ref_iv.strand = "+"
            for iv,val in genes[cigop.ref_iv].steps():
                if inter_count_1 == 0:
                    gene_id_intersect_1 |= val
                    inter_count_1 += 1
                else:
                    gene_id_intersect_1 &= val

        gene_id_intersect_2 = set()
        inter_count_2 = 0
        for cigop in second_almnt.cigar:
            if cigop.type != "M":
                continue
            for iv,val in genes[cigop.ref_iv].steps():
                if inter_count_2 == 0:
                    gene_id_intersect_2 |= val
                    inter_count_2 += 1
                else:
                    gene_id_intersect_2 &= val


        
        
        
        if len(gene_id_intersect_1) > 0 or len(gene_id_intersect_2) > 0:
            if len(gene_id_intersect_1) == 0:
                counts["_no_feature"] += 1
                genes_2 = [l.split('-')[0] for l in ','.join(list(gene_id_intersect_2)).split(",")]
                if len(np.unique(np.array(genes_2))) == 1:
                    perfect_inter_exon += 1
                    for i in range(0,len(gene_id_intersect_2)):
                        gene_id = list(gene_id_intersect_2)[i]
                        counts[gene_id] += 1/len(gene_id_intersect_2)
                else:
                    counts["_ambiguous"] += 1
            elif len(gene_id_intersect_2) == 0:
                counts["_no_feature"] += 1
                genes_1 = [l.split('-')[0] for l in ','.join(list(gene_id_intersect_1)).split(",")]
                if len(np.unique(np.array(genes_1))) == 1:
                    perfect_inter_exon += 1
                    for i in range(0,len(gene_id_intersect_1)):
                        gene_id = list(gene_id_intersect_1)[i]
                        counts[gene_id] += 1/len(gene_id_intersect_1)
                else:
                    counts["_ambiguous"] += 1
            elif len(gene_id_intersect_1) == 1 and len(gene_id_intersect_2) == 1:
                genes_1 = [l.split('-')[0] for l in ','.join(list(gene_id_intersect_1)).split(",")]
                genes_2 = [l.split('-')[0] for l in ','.join(list(gene_id_intersect_2)).split(",")]
                if genes_1 == genes_2:
                        gene_id = list(gene_id_intersect_1)[0]
                        counts[gene_id] += 1
                        perfect_inter_exon += 1
                        gene_id = list(gene_id_intersect_2)[0]
                        counts[gene_id] += 1
                        perfect_inter_exon += 1
                else:
                    counts["_ambiguous"] += 2
            elif len(gene_id_intersect_1) > 1 and len(gene_id_intersect_2) > 1:
                genes_1 = [l.split('-')[0] for l in ','.join(list(gene_id_intersect_1)).split(",")]
                genes_2 = [l.split('-')[0] for l in ','.join(list(gene_id_intersect_2)).split(",")]               
                intersection = list(set(genes_1) & set(genes_2))
                if len(intersection) == 1:
                    gene_id_1 = list(np.array(list(gene_id_intersect_1))[list(np.where(np.array(genes_1) == intersection[0])[0])])
                    gene_id_2 = list(np.array(list(gene_id_intersect_2))[list(np.where(np.array(genes_2) == intersection[0])[0])])
                    perfect_inter_exon += 2
                    for i in range(0,len(gene_id_1)):
                        gene_id = list(gene_id_1)[i]
                        counts[gene_id] += 1/len(gene_id_1)
                    for i in range(0,len(gene_id_2)):
                        gene_id = list(gene_id_2)[i]
                        counts[gene_id] += 1/len(gene_id_2)                
                else:
                    counts["_ambiguous"] += 2
            elif len(gene_id_intersect_1) > 1:
                genes_1 = [l.split('-')[0] for l in ','.join(list(gene_id_intersect_1)).split(",")]
                genes_2 = [l.split('-')[0] for l in ','.join(list(gene_id_intersect_2)).split(",")]
                if(genes_2[0] in genes_1):
                    gene_id = list(gene_id_intersect_2)[0]
                    counts[gene_id] += 1
                    perfect_inter_exon += 2
                    gene_id_1 = list(np.array(list(gene_id_intersect_1))[list(np.where(np.array(genes_1) == genes_2[0])[0])])
                    for i in range(0,len(gene_id_1)):
                        gene_id = list(gene_id_1)[i]
                        counts[gene_id] += 1/len(gene_id_1)
                else:
                    gene_id = list(gene_id_intersect_2)[0]
                    counts[gene_id] += 1
                    perfect_inter_exon += 1
                    counts["_ambiguous"] += 1               
            elif len(gene_id_intersect_2) > 1:
                genes_1 = [l.split('-')[0] for l in ','.join(list(gene_id_intersect_1)).split(",")]
                genes_2 = [l.split('-')[0] for l in ','.join(list(gene_id_intersect_2)).split(",")]
                if(genes_1[0] in genes_2):
                    gene_id = list(gene_id_intersect_1)[0]
                    counts[gene_id] += 1
                    perfect_inter_exon += 2
                    gene_id_2 = list(np.array(list(gene_id_intersect_2))[list(np.where(np.array(genes_2) == genes_1[0])[0])])
                    for i in range(0,len(gene_id_2)):
                        gene_id = list(gene_id_2)[i]
                        counts[gene_id] += 1/len(gene_id_2)                 
                else:
                    gene_id = list(gene_id_intersect_1)[0]
                    counts[gene_id] += 1
                    perfect_inter_exon += 1
                    counts["_ambiguous"] += 1         
        else:
            gene_id_intersect_1, gene_id_intersect_2 = test_opposite_strand(first_almnt, second_almnt, genes, 1)    
            if len(gene_id_intersect_1) > 0 or len(gene_id_intersect_2) > 0:
                if len(gene_id_intersect_1) == 0:
                    counts["_no_feature"] += 1
                    genes_2 = [l.split('-')[0] for l in ','.join(list(gene_id_intersect_2)).split(",")]
                    if len(np.unique(np.array(genes_2))) == 1:
                        perfect_inter_exon += 1
                        for i in range(0,len(gene_id_intersect_2)):
                            gene_id = list(gene_id_intersect_2)[i]
                            counts[gene_id] += 1/len(gene_id_intersect_2)
                    else:
                        counts["_ambiguous"] += 1
                elif len(gene_id_intersect_2) == 0:
                    counts["_no_feature"] += 1
                    genes_1 = [l.split('-')[0] for l in ','.join(list(gene_id_intersect_1)).split(",")]
                    if len(np.unique(np.array(genes_1))) == 1:
                        perfect_inter_exon += 1
                        for i in range(0,len(gene_id_intersect_1)):
                            gene_id = list(gene_id_intersect_1)[i]
                            counts[gene_id] += 1/len(gene_id_intersect_1)
                    else:
                        counts["_ambiguous"] += 1
                elif len(gene_id_intersect_1) == 1 and len(gene_id_intersect_2) == 1:
                    genes_1 = [l.split('-')[0] for l in ','.join(list(gene_id_intersect_1)).split(",")]
                    genes_2 = [l.split('-')[0] for l in ','.join(list(gene_id_intersect_2)).split(",")]
                    if genes_1 == genes_2:
                            gene_id = list(gene_id_intersect_1)[0]
                            counts[gene_id] += 1
                            perfect_inter_exon += 1
                            gene_id = list(gene_id_intersect_2)[0]
                            counts[gene_id] += 1
                            perfect_inter_exon += 1
                    else:
                        counts["_ambiguous"] += 2
                elif len(gene_id_intersect_1) > 1 and len(gene_id_intersect_2) > 1:
                    genes_1 = [l.split('-')[0] for l in ','.join(list(gene_id_intersect_1)).split(",")]
                    genes_2 = [l.split('-')[0] for l in ','.join(list(gene_id_intersect_2)).split(",")]               
                    intersection = list(set(genes_1) & set(genes_2))
                    if len(intersection) == 1:
                        gene_id_1 = list(np.array(list(gene_id_intersect_1))[list(np.where(np.array(genes_1) == intersection[0])[0])])
                        gene_id_2 = list(np.array(list(gene_id_intersect_2))[list(np.where(np.array(genes_2) == intersection[0])[0])])
                        perfect_inter_exon += 2
                        for i in range(0,len(gene_id_1)):
                            gene_id = list(gene_id_1)[i]
                            counts[gene_id] += 1/len(gene_id_1)
                        for i in range(0,len(gene_id_2)):
                            gene_id = list(gene_id_2)[i]
                            counts[gene_id] += 1/len(gene_id_2)                
                    else:
                        counts["_ambiguous"] += 2
                elif len(gene_id_intersect_1) > 1:
                    genes_1 = [l.split('-')[0] for l in ','.join(list(gene_id_intersect_1)).split(",")]
                    genes_2 = [l.split('-')[0] for l in ','.join(list(gene_id_intersect_2)).split(",")]
                    if(genes_2[0] in genes_1):
                        gene_id = list(gene_id_intersect_2)[0]
                        counts[gene_id] += 1
                        perfect_inter_exon += 2
                        gene_id_1 = list(np.array(list(gene_id_intersect_1))[list(np.where(np.array(genes_1) == genes_2[0])[0])])
                        for i in range(0,len(gene_id_1)):
                            gene_id = list(gene_id_1)[i]
                            counts[gene_id] += 1/len(gene_id_1)
                    else:
                        gene_id = list(gene_id_intersect_2)[0]
                        counts[gene_id] += 1
                        perfect_inter_exon += 1
                        counts["_ambiguous"] += 1               
                elif len(gene_id_intersect_2) > 1:
                    genes_1 = [l.split('-')[0] for l in ','.join(list(gene_id_intersect_1)).split(",")]
                    genes_2 = [l.split('-')[0] for l in ','.join(list(gene_id_intersect_2)).split(",")]
                    if(genes_1[0] in genes_2):
                        gene_id = list(gene_id_intersect_1)[0]
                        counts[gene_id] += 1
                        perfect_inter_exon += 2
                        gene_id_2 = list(np.array(list(gene_id_intersect_2))[list(np.where(np.array(genes_2) == genes_1[0])[0])])
                        for i in range(0,len(gene_id_2)):
                            gene_id = list(gene_id_2)[i]
                            counts[gene_id] += 1/len(gene_id_2)                 
                    else:
                        gene_id = list(gene_id_intersect_1)[0]
                        counts[gene_id] += 1
                        perfect_inter_exon += 1
                        counts["_ambiguous"] += 1 
            else:
                counts["_no_feature"] += 1
                    
    print("File name: ", sam_file)
    print("1: Intersect exon match: ", perfect_inter_exon)
    print("2: Ambiguous match for exons: ", counts["_ambiguous"])
    print("3: No match: ",  counts["_no_feature"])
    print("Sam file analysis finished~")
    
    with open(report, 'w') as report:
        report.write("1" + "," + str(cell_ID) + "," + str(perfect_inter_exon) + "\n")
        report.write("2" + "," + str(cell_ID) + "," + str(counts["_ambiguous"]) + "\n")
        report.write("3" + "," + str(cell_ID) + "," + str(counts["_no_feature"]) + "\n")
        
    with open(count_output, 'w') as count_output:
        for gene in counts:
            if (gene in ["_unmapped", "_ambiguous", "_no_feature"]):
                continue
            else:
                line = str(gene_annotat.loc[gene,3]) + "," + str(cell_ID) + "," + str(counts[gene]) + "\n"
                count_output.write(line)
    return 0


def test_opposite_strand(first_almnt_tmp, second_almnt_tmp, genes_tmp, exongene):
    gene_id_intersect_tmp_1 = set()
    inter_count_tmp_1 = 0
    
    for cigop in first_almnt_tmp.cigar:
        if cigop.type != "M":
            continue
        if exongene == 1:
            if cigop.ref_iv.strand == "+":
                cigop.ref_iv.strand = "-"
            else:
                cigop.ref_iv.strand = "+"                
        for iv,val in genes_tmp[cigop.ref_iv].steps():
            if inter_count_tmp_1 == 0:
                gene_id_intersect_tmp_1 |= val
                inter_count_tmp_1 += 1
            else:
                gene_id_intersect_tmp_1 &= val

    gene_id_intersect_tmp_2 = set()
    inter_count_tmp_2 = 0
    for cigop in second_almnt_tmp.cigar:
        if cigop.type != "M":
            continue
        if exongene == 1:
            if cigop.ref_iv.strand == "+":
                cigop.ref_iv.strand = "-"
            else:
                cigop.ref_iv.strand = "+"
        for iv,val in genes_tmp[cigop.ref_iv].steps():
            if inter_count_tmp_2 == 0:
                gene_id_intersect_tmp_2 |= val
                inter_count_tmp_2 += 1
            else:
                gene_id_intersect_tmp_2 &= val
                
    return(gene_id_intersect_tmp_1, gene_id_intersect_tmp_2)


def sciRNA_count_parallel(gtf_file, input_folder, sample_ID, core_number):
    # read in the gtf file, and then construct the genome interval for exons, genes, and gene end dictionary
    gtf_file = HTSeq.GFF_Reader(gtf_file, end_included=True)
    gene_annotat_file = input_folder + "/gene_name_annotate.txt"
    cell_annotat_file = input_folder + "/cell_annotate.txt"
    report_annotate_file = input_folder + "/report_annotate.txt"
    
    gene_annotat = open(gene_annotat_file, "w")
    cell_annotat = open(cell_annotat_file, "w")
    report_annotate = open(report_annotate_file, "w")
    
    genes = HTSeq.GenomicArrayOfSets( "auto", stranded=True )
    gene_n = 0
    gene_count = 0
    print("Start generating exon genomic arrays....")

    for feature in gtf_file:
        if feature.type == "gene":
            gene_n +=1
            genes[ feature.iv ] += feature.attr["gene_id"]
            gene_count += 1
            
            # for human and mouse gtf file
            message = (feature.attr["gene_id"] + "," + feature.attr["gene_type"] + "," 
                        + feature.attr["gene_name"] + "," + str(gene_count) + "\n")
            
            gene_annotat.write(message)
            

    print("Detected exon number: ", gene_n)
    
    gene_annotat.close()
    
    gene_annotat = pd.read_csv(gene_annotat_file, header=None)
    gene_annotat.index =  gene_annotat[0]
    
    sample_ID = list(pd.read_csv(sample_ID, header=None)[0])
    
    # generate the cell ID annotate file
    cell_count = 0
    for i in sample_ID:
        cell_count += 1
        message = i + "," + str(cell_count) + "\n"
        cell_annotat.write(message)
    cell_annotat.close()
    
    # Generate the report annotate file
    report_annotate.write("1, Intersect exon match\n")
    report_annotate.write("2, Ambiguous match for exons\n")
    report_annotate.write("3, No match\n")
    report_annotate.close()


    # parallele for the functions
    p = Pool(processes = int(core_number))
    #print("Processing core number: ", core_number)
    func = partial(sciRNAseq_count, input_folder = input_folder, genes=genes, gene_annotat = gene_annotat, sample_ID = sample_ID)
    #sciRNAseq_count(sample, input_folder, exons, genes, gene_end)
    result = p.map(func, sample_ID)
    p.close()
    p.join()

    print("All analysis done~")

    
if __name__ == "__main__":
    gtf_file = sys.argv[1]
    input_folder = sys.argv[2]
    sample_ID = sys.argv[3]
    core_number = sys.argv[4]
    sciRNA_count_parallel(gtf_file, input_folder, sample_ID, core_number)
```
