## Supplementary file 3 for "A global view of aging and Alzheimer’s pathogenesis-associated cell population dynamics and molecular signatures in the human and mouse brains": filtering.sh

```
input_folder=$1
sample=$2
output_folder=$3

echo Filtering sample: $sample
samtools view -q 30 -f 2 -F 780 $input_folder/$sample*.sam | sort --parallel=10 -k1,1 -k3,3 -k4,4n > $output_folder/$sample.noheader.sam
grep "@" $input_folder/$sample*.sam > $output_folder/$sample.header.sam
cat $output_folder/$sample.header.sam  $output_folder/$sample.noheader.sam > $output_folder/$sample.sam
rm $output_folder/$sample.header.sam
rm $output_folder/$sample.noheader.sam
echo Filtering $sample done.
```
