## Supplementary file 3 for "A global view of aging and Alzheimer’s pathogenesis-associated cell population dynamics and molecular signatures in the human and mouse brains": gene_counting.py

```
import itertools
import collections
import numpy as np
import pandas as pd
from multiprocessing import Pool
from multiprocessing import *
import HTSeq
import sys
from functools import partial
import logging


def sciRNAseq_count(sample, input_folder, exons, genes, gene_end, gene_annotat, sample_ID, randomN_barcodes):
    input_sam = input_folder + "/" + sample + ".sam"
    report = input_folder + "/" + sample + ".report"
    count_output = input_folder + "/" + sample + ".count"

    sample_barcode = sample.split(".")[1][10:20]    
    RT_barcode = "shortDT"
    if sample_barcode in randomN_barcodes:
        RT_barcode = "randomN"
    
    counts = collections.Counter()
    sam_file = input_sam
    almnt_file = HTSeq.SAM_Reader(sam_file)
    sam_name = sample
    cell_ID = sample_ID.index(sample) + 1

    perfect_inter_exon = 0
    nearest_inter_exon = 0
    perfect_combine_exon = 0
    nearest_combine_exon = 0
    perfect_inter_gene = 0
    nearest_inter_gene = 0
    perfect_combine_gene = 0
    nearest_combine_gene = 0
    
    print("Start read the input file: " + sam_file + "....")

    for bundle in HTSeq.pair_SAM_alignments( almnt_file, bundle=True ):

        if len(bundle) != 1:
            continue  # Skip multiple alignments
        first_almnt, second_almnt = bundle[0]  # extract pair
        if (not first_almnt.aligned) or (not second_almnt.aligned):
            count[ "_unmapped" ] += 1
            continue
        if (first_almnt.iv.chrom not in genes.chrom_vectors) or (second_almnt.iv.chrom not in genes.chrom_vectors):
            counts["_unmapped"] += 1
            continue

        if first_almnt.iv.strand == "+":
            first_almnt.iv.strand = "-"
        else:
            first_almnt.iv.strand = "+"        


        # First check the intersectin with exons
        gene_id_intersect = set()
        gene_id_combine = set()
        inter_count = 0
        for cigop in first_almnt.cigar:
            if cigop.type != "M":
                continue
            if cigop.ref_iv.strand == "+":
                cigop.ref_iv.strand = "-"
            else:
                cigop.ref_iv.strand = "+"
            for iv,val in exons[cigop.ref_iv].steps():
                gene_id_combine |= val
                if inter_count == 0:
                    gene_id_intersect |= val
                    inter_count += 1
                else:
                    gene_id_intersect &= val
        
        for cigop in second_almnt.cigar:
            if cigop.type != "M":
                continue

            for iv,val in exons[cigop.ref_iv].steps():
                gene_id_combine |= val
                if inter_count == 0:
                    gene_id_intersect |= val
                    inter_count += 1
                else:
                    gene_id_intersect &= val


        # first check the intersection set
        if len(gene_id_intersect) == 1:
            gene_id = list(gene_id_intersect)[0]
            counts[gene_id] += 1
            perfect_inter_exon += 1
        elif len(gene_id_intersect) > 1:
            gene_id = find_nearest_gene(first_almnt.iv.end_d, gene_id_intersect, gene_end, RT_barcode)
            counts[gene_id] += 1
            nearest_inter_exon += 1
        else:
            # if there no intersection match, then find the union sets
            if len(gene_id_combine) == 1:
                gene_id = list(gene_id_combine)[0]
                counts[gene_id] += 1
                perfect_combine_exon += 1
            elif len(gene_id_combine) > 1:
                gene_id = find_nearest_gene(first_almnt.iv.end_d, gene_id_combine, gene_end, RT_barcode)
                counts[gene_id] += 1
                nearest_combine_exon += 1
            else:
                # if there is no intersection match or union match, then search for genes to find the intronic match
                gene_id_intersect = set()
                gene_id_combine = set()
                inter_count = 0
                for cigop in first_almnt.cigar:
                    if cigop.type != "M":
                        continue
                    for iv,val in genes[cigop.ref_iv].steps():
                        gene_id_combine |= val
                        if inter_count == 0:
                            gene_id_intersect |= val
                            inter_count += 1
                        else:
                            gene_id_intersect &= val

                for cigop in second_almnt.cigar:
                    if cigop.type != "M":
                        continue
                    for iv,val in genes[cigop.ref_iv].steps():
                        gene_id_combine |= val
                        if inter_count == 0:
                            gene_id_intersect |= val
                            inter_count += 1
                        else:
                            gene_id_intersect &= val


                exonintron = "_intron"
                if len(gene_id_intersect) == 0 and len(gene_id_combine) == 0:
                    gene_id_combine, gene_id_intersect = test_flanking_region(first_almnt, second_almnt, genes)
                if len(gene_id_intersect) == 0 and len(gene_id_combine) == 0:
                    gene_id_combine, gene_id_intersect = test_opposite_strand(first_almnt, second_almnt, exons, 1)
                    if len(gene_id_intersect) > 0 or len(gene_id_combine) > 0:
                        exonintron = ""
                if len(gene_id_intersect) == 0 and len(gene_id_combine) == 0:
                    gene_id_combine, gene_id_intersect = test_opposite_strand(first_almnt, second_almnt, genes, 2)
 


                if len(gene_id_intersect) == 1:
                    gene_id = list(gene_id_intersect)[0] + exonintron
                    counts[gene_id] += 1
                    perfect_inter_gene += 1

                elif len(gene_id_intersect) > 1:
                    gene_id = find_nearest_gene(first_almnt.iv.end_d, gene_id_intersect, gene_end, RT_barcode) + exonintron
                    counts[gene_id] += 1
                    nearest_inter_gene += 1

                else:
                    # if there no intersection match, then find the union sets
                    if len(gene_id_combine) == 1:
                        gene_id = list(gene_id_combine)[0] + exonintron
                        counts[gene_id] += 1
                        perfect_combine_gene += 1

                    elif len(gene_id_combine) > 1:
                        gene_id = find_nearest_gene(first_almnt.iv.end_d, gene_id_combine, gene_end, RT_barcode) + exonintron
                        counts[gene_id] += 1
                        nearest_combine_gene += 1

                    else:
                        counts["_no_feature"] += 1
                        
    print("File name: ", sam_file)
    print("1: Perfect intersect exon match: ", perfect_inter_exon)
    print("2: Nearest intersect exon match: ", nearest_inter_exon)
    print("3: Perfect combine exon match: ", perfect_combine_exon)
    print("4: Nearest combine exon match: ", nearest_combine_exon)
    print("5: Perfect intersect gene match: ", perfect_inter_gene)
    print("6: Nearest intersect gene match: ", nearest_inter_gene)
    print("7: Perfect combine gene match: ", perfect_combine_gene)
    print("8: Nearest combine gene match: ", nearest_combine_gene)
    print("9: ambiguous match for exons: ", counts["_ambiguous"])
    print("10: ambiguous match for genes: ", counts["_ambiguous_intron"])
    print("11: No match: ",  counts["_no_feature"])
    print("Sam file analysis finished~")
    
    with open(report, 'w') as report:
        report.write("1" + "," + str(cell_ID) + "," + str(perfect_inter_exon) + "\n")
        report.write("2" + "," + str(cell_ID) + "," + str(nearest_inter_exon) + "\n")
        report.write("3" + "," + str(cell_ID) + "," + str(perfect_combine_exon) + "\n")
        report.write("4" + "," + str(cell_ID) + "," + str(nearest_combine_exon) + "\n")
        report.write("5" + "," + str(cell_ID) + "," + str(perfect_inter_gene) + "\n")
        report.write("6" + "," + str(cell_ID) + "," + str(nearest_inter_gene) + "\n")
        report.write("7" + "," + str(cell_ID) + "," + str(perfect_combine_gene) + "\n")
        report.write("8" + "," + str(cell_ID) + "," + str(nearest_combine_gene) + "\n")
        report.write("9" + "," + str(cell_ID) + "," + str(counts["_ambiguous"]) + "\n")
        report.write("10" + "," + str(cell_ID) + "," + str(counts["_ambiguous_intron"]) + "\n")
        report.write("11" + "," + str(cell_ID) + "," + str(counts["_no_feature"]) + "\n")
        
    with open(count_output, 'w') as count_output:
        for gene in counts:
            if (gene in ["_unmapped", "_ambiguous", "_ambiguous_intron", "_no_feature"]):
                continue
            else:
                line = str(gene_annotat.loc[gene,4]) + "," + str(cell_ID) + "," + str(counts[gene]) + "\n"
                count_output.write(line)
    return 0

# Define a function that accept the end position of the alignments, and a set of genes, and the dictionary of the
# gene end, and then return the gene with the nearest end to the alignment

# for each gene element in the gene_id_intersect, I will find the absolute distance between the alignment end and
# the transcript end, and select the minimum distance for each gene in the list, and then compare the genes, and use
# the nearest gene as the gene id for output

def find_nearest_gene(al_end, gene_id_intersect, gene_end, RT_barcode):
    gene_id_end = {}
    for gene in gene_id_intersect:
        if gene in gene_end:
            gene_id_end[gene] = (abs(np.array(list(gene_end[gene])) - al_end)).min()
        else:
            print "****************Found one gene without transcript annotation*****************", "Gene name: ", gene
    # filter the gene with the least distance. If there are two genes with the least distance, then  "_ambiguous" 
    # would be returned
    gene_end_min = np.min(gene_id_end.values())
    count = 0
    #print ("gene end distance: ", gene_id_end.values())
    for gene in gene_id_end:
        if (gene_id_end[gene] < (gene_end_min + 100)):
            count += 1
            gene_id = gene
    if count > 1:
        gene_id = "_ambiguous"
    
    if RT_barcode == "randomN":
        gene_id = "_ambiguous"
    
    return(gene_id)


def test_opposite_strand(first_almnt_tmp, second_almnt_tmp, genes_tmp, exongene):
    gene_id_intersect_tmp = set()
    gene_id_combine_tmp = set()
    inter_count_tmp = 0
    
    for cigop in first_almnt_tmp.cigar:
        if cigop.type != "M":
            continue
        if exongene == 1:
            if cigop.ref_iv.strand == "+":
                cigop.ref_iv.strand = "-"
            else:
                cigop.ref_iv.strand = "+"                
        for iv,val in genes_tmp[cigop.ref_iv].steps():
            gene_id_combine_tmp |= val
            if inter_count_tmp == 0:
                gene_id_intersect_tmp |= val
                inter_count_tmp += 1
            else:
                gene_id_intersect_tmp &= val

    for cigop in second_almnt_tmp.cigar:
        if cigop.type != "M":
            continue
        if exongene == 1:
            if cigop.ref_iv.strand == "+":
                cigop.ref_iv.strand = "-"
            else:
                cigop.ref_iv.strand = "+"
        for iv,val in genes_tmp[cigop.ref_iv].steps():
            gene_id_combine_tmp |= val
            if inter_count_tmp == 0:
                gene_id_intersect_tmp |= val
                inter_count_tmp += 1
            else:
                gene_id_intersect_tmp &= val
                
    return(gene_id_combine_tmp, gene_id_intersect_tmp)


def test_flanking_region(first_almnt_tmp, second_almnt_tmp, genes_tmp):
    gene_id_intersect_tmp = set()
    gene_id_combine_tmp = set()
    inter_count_tmp = 0
    
    if second_almnt_tmp.iv.strand == "+":
        
        if (second_almnt_tmp.iv.start - 1000) >= 0:
            difference = 1000
        else:
            difference = second_almnt_tmp.iv.start
     
        for cigop in second_almnt_tmp.cigar:
            if cigop.type != "M":
                continue

            cigop.ref_iv.start =  cigop.ref_iv.start - difference
            cigop.ref_iv.end =  cigop.ref_iv.end - difference
    
            for iv,val in genes_tmp[cigop.ref_iv].steps():
                gene_id_combine_tmp |= val
                if inter_count_tmp == 0:
                    gene_id_intersect_tmp |= val
                    inter_count_tmp += 1
                else:
                    gene_id_intersect_tmp &= val
            cigop.ref_iv.start = cigop.ref_iv.start + difference
            cigop.ref_iv.end = cigop.ref_iv.end + difference


        if (first_almnt_tmp.iv.start - difference) < 0:
            difference = first_almnt_tmp.iv.start
            
        for cigop in first_almnt_tmp.cigar:
            if cigop.type != "M":
                continue

            cigop.ref_iv.start =  cigop.ref_iv.start - difference
            cigop.ref_iv.end =  cigop.ref_iv.end - difference

            for iv,val in genes_tmp[cigop.ref_iv].steps():
                gene_id_combine_tmp |= val
                if inter_count_tmp == 0:
                    gene_id_intersect_tmp |= val
                    inter_count_tmp += 1
                else:
                    gene_id_intersect_tmp &= val
            cigop.ref_iv.start = cigop.ref_iv.start + difference
            cigop.ref_iv.end = cigop.ref_iv.end + difference

    else:
    
        for cigop in first_almnt_tmp.cigar:
            if cigop.type != "M":
                continue
            cigop.ref_iv.start = cigop.ref_iv.start + 1000
            cigop.ref_iv.end = cigop.ref_iv.end + 1000
            for iv,val in genes_tmp[cigop.ref_iv].steps():
                gene_id_combine_tmp |= val
                if inter_count_tmp == 0:
                    gene_id_intersect_tmp |= val
                    inter_count_tmp += 1
                else:
                    gene_id_intersect_tmp &= val
            cigop.ref_iv.start = cigop.ref_iv.start - 1000
            cigop.ref_iv.end = cigop.ref_iv.end - 1000

        for cigop in second_almnt_tmp.cigar:
            if cigop.type != "M":
                continue
            cigop.ref_iv.start = cigop.ref_iv.start + 1000
            cigop.ref_iv.end = cigop.ref_iv.end + 1000
            for iv,val in genes_tmp[cigop.ref_iv].steps():
                gene_id_combine_tmp |= val
                if inter_count_tmp == 0:
                    gene_id_intersect_tmp |= val
                    inter_count_tmp += 1
                else:
                    gene_id_intersect_tmp &= val
            cigop.ref_iv.start = cigop.ref_iv.start - 1000
            cigop.ref_iv.end = cigop.ref_iv.end - 1000
    
    return(gene_id_combine_tmp, gene_id_intersect_tmp)


'''
def sciRNAseq_count_mf(sample, input_folder, exons, genes, gene_end):
    try:
        return sciRNAseq_count(sample, input_folder, exons, genes, gene_end)
    except Exception:
        #logging.exception("f(%r) failed" % (sample, input_folder, exons, genes, gene_end,))
        print Exception.args()
'''

def sciRNA_count_parallel(gtf_file, input_folder, sample_ID, core_number, randomN_barcode_file):
    # read in the gtf file, and then construct the genome interval for exons, genes, and gene end dictionary
    gtf_file = HTSeq.GFF_Reader(gtf_file, end_included=True)
    gene_annotat_file = input_folder + "/gene_name_annotate.txt"
    cell_annotat_file = input_folder + "/cell_annotate.txt"
    report_annotate_file = input_folder + "/report_annotate.txt"
    
    gene_annotat = open(gene_annotat_file, "w")
    cell_annotat = open(cell_annotat_file, "w")
    report_annotate = open(report_annotate_file, "w")
    
    exons = HTSeq.GenomicArrayOfSets( "auto", stranded=True )
    genes = HTSeq.GenomicArrayOfSets( "auto", stranded=True )
    gene_end = {}
    exon_n = 0
    gene_n = 0
    transcript_n = 0
    gene_count = 0
    print("Start generating exon genomic arrays....")
    print("Start generating gene genomic arrays....")
    print("Start calculating transcript end of genes....")

    for feature in gtf_file:
        if feature.type == "exon":
            exon_n += 1
            exons[ feature.iv ] += feature.attr["gene_id"]
        elif feature.type == "gene":
            gene_n +=1
            genes[ feature.iv ] += feature.attr["gene_id"]
            gene_count += 1
            
            # for human and mouse gtf file
            message = (feature.attr["gene_id"] + "," + feature.attr["gene_type"] + "," 
                       + "exon" + "," + feature.attr["gene_name"] + "," + str(gene_count) + "\n")
            
            gene_annotat.write(message)
            
            gene_count += 1
            
            # for human and mouse gtf file
            message = (feature.attr["gene_id"] + "_intron" + "," + feature.attr["gene_type"] + "," 
                       + "intron" + "," + feature.attr["gene_name"] + "_intron" + "," + str(gene_count) + "\n")
            gene_annotat.write(message)
            
        elif feature.type == "transcript":
            transcript_n += 1
            #print "feature gene name: ", feature.attr["gene_id"]
            if feature.attr["gene_id"] in gene_end.keys():
                gene_end[ feature.attr["gene_id"] ].add(feature.iv.end_d)
            else:
                gene_end[ feature.attr["gene_id"] ] = set()
                gene_end[ feature.attr["gene_id"] ].add(feature.iv.end_d)

    print("Detected gene number: ", gene_n)
    print("Detected transcript number: ", transcript_n)
    print("Detected exon number: ", exon_n)
    
    gene_annotat.close()
    
    gene_annotat = pd.read_csv(gene_annotat_file, header=None)
    gene_annotat.index =  gene_annotat[0]
    #print("print WBGENE id:", gene_annotat.loc["WBGene00004947", 4])
    #print("Print transcript end, ", len(gene_end))
    
    sample_ID = list(pd.read_csv(sample_ID, header=None)[0])
    
    # generate the cell ID annotate file
    cell_count = 0
    for i in sample_ID:
        cell_count += 1
        message = i + "," + str(cell_count) + "\n"
        cell_annotat.write(message)
    cell_annotat.close()
    
    # Generate the report annotate file
    report_annotate.write("1, Perfect intersect exon match\n")
    report_annotate.write("2, Nearest intersect exon match\n")
    report_annotate.write("3, Perfect combine exon match\n")
    report_annotate.write("4, Nearest combine exon match\n")
    report_annotate.write("5, Perfect intersect gene match\n")
    report_annotate.write("6, Nearest intersect gene match\n")
    report_annotate.write("7, Perfect combine gene match\n")
    report_annotate.write("8, Nearest combine gene match\n")
    report_annotate.write("9, ambiguous match for exons\n")
    report_annotate.write("10, ambiguous match for genes\n")
    report_annotate.write("11, No match\n")
    report_annotate.close()
    

    # generate the randomN barcode list:
    barcodes = open(randomN_barcode_file, "rb")
    with barcodes as f:
        randomN_barcodes = f.read().splitlines()
    barcodes.close()


    # parallele for the functions
    p = Pool(processes = int(core_number))
    #print("Processing core number: ", core_number)
    func = partial(sciRNAseq_count, input_folder = input_folder, exons=exons, genes=genes, gene_end = gene_end, gene_annotat = gene_annotat, sample_ID = sample_ID, randomN_barcodes = randomN_barcodes)
    #sciRNAseq_count(sample, input_folder, exons, genes, gene_end)
    result = p.map(func, sample_ID)
    p.close()
    p.join()

    print("All analysis done~")
'''
def p_sciRNAseq_count(sample):
    print("get sample: ", sample)
    input_sam = input_folder + "/" + sample + ".sam"
    report = input_folder + "/" + sample + ".report"
    count_output = input_folder + "/" + sample + ".count"
    sciRNAseq_count(sample, input_sam, report, count_output, exons, genes, gene_end)

'''
    
if __name__ == "__main__":
    gtf_file = sys.argv[1]
    input_folder = sys.argv[2]
    sample_ID = sys.argv[3]
    core_number = sys.argv[4]
    randomN_barcode_file = sys.argv[5]
    sciRNA_count_parallel(gtf_file, input_folder, sample_ID, core_number, randomN_barcode_file)
```
