## Supplementary file 3 for "A global view of aging and Alzheimer’s pathogenesis-associated cell population dynamics and molecular signatures in the human and mouse brains": generate_barcode_dictionary.py

```
RT_bc_file = ./RT_barcodes.txt
lig_bc_file = ./Ligation_barcodes.txt

import itertools
from Levenshtein import distance
# for accept a list of characters ["A", "C", "G", "T"] and a length generate all permutations 
def combination_seqs(input_char, output_length):
    all_combine = list(map(lambda x: list(map(lambda x: "".join(x), set(itertools.permutations("".join(x))))), 
                           list(itertools.combinations_with_replacement(input_char, output_length))))
    return(list(itertools.chain.from_iterable(all_combine)))

# Here I am going to accept the permuted sequence list and the target list and then generate
# a dictionary including the permuted sequence and the matching sequence (within 1 edit distance)
# if there is no sequence matching, then return ""

def find_bc(seq, bc_list, mismatch_rate = 1):
    # in this script, I am going to accept a sequence and a ligation barcode list, and then
    # find if there is a matching barcode in the bc_list matching the first several bases of
    # the seq
    result = ""
    for bc in bc_list:
        mismatch = distance(bc, seq)
        if mismatch <= mismatch_rate:
            result = bc
            break
    return(result)

    
# generate all possible sequences of 10bp
print("Generating all possible sequences...")
all_seq_10bp = combination_seqs(["A", "C", "G", "T", "N"], 10)
RT_bc = open(RT_bc_file)
RT = list(map(lambda x: x.strip(), RT_bc.readlines()))
RT_bc.close()
print("Generating the dictionary for RT barcodes...")
# generat the dictionary for RT barcodes
all_seq_match = list(map(lambda x: find_bc(x, RT), all_seq_10bp))
all_seq_dic = dict(zip(all_seq_10bp, all_seq_match))
# save the dictionary into a file
import pickle
pickle_out = open("./RT_barcodes.pickle","wb")
pickle.dump(all_seq_dic, pickle_out)
pickle_out.close()
print("Generating the dictionary for ligation barcodes...")
# also generate the dictionary for ligation barcodes
lig_bc = open(lig_bc_file)
lig = list(map(lambda x: x.strip(), lig_bc.readlines()))
lig_bc.close()
all_seq_match = list(map(lambda x: find_bc(x, lig), all_seq_10bp))
all_seq_dic = dict(zip(all_seq_10bp, all_seq_match))
pickle_out = open("./Ligation_barcodes.pickle","wb")
pickle.dump(all_seq_dic, pickle_out)
pickle_out.close()

# generate the simplified dictionary
pickle_in = open("./Ligation_barcodes.pickle", "rb")
lig_dic = pickle.load(pickle_in)
lig_filter = {k: v for k, v in lig_dic.items() if v != ""}
pickle_out = open("./Ligation_barcodes.pickle2", "wb")
pickle.dump(lig_filter, pickle_out, 2)
pickle_in.close()
pickle_out.close()

pickle_in = open("./RT_barcodes.pickle", "rb")
RT_dic = pickle.load(pickle_in)
RT_filter = {k: v for k, v in RT_dic.items() if v != ""}
pickle_out = open("./RT_barcodes.pickle2", "wb")
pickle.dump(RT_filter, pickle_out, 2)
pickle_in.close()
```
