## Supplementary file 3 for "A global view of aging and Alzheimer’s pathogenesis-associated cell population dynamics and molecular signatures in the human and mouse brains": generate_barcode_dictionary.sh

```
#! /bin/bash
#SBATCH -N 1
## number of cores 
#SBATCH -n 1
#SBATCH -o ./Log_file.out

bashrc_location=/ru-auth/local/home/asziraki/projects/AS_20200820_single_cell_pipeline/Scripts/original_pipeline/sci-RNA-seq3_pipeline/.bashrc
source $bashrc_location
conda activate original_pipeline

python generate_barcode_dictionary.py

conda deactivate
```
