## Supplementary file 3 for "A global view of aging and Alzheimer’s pathogenesis-associated cell population dynamics and molecular signatures in the human and mouse brains": sam_split.sh

```
input_folder=$1
sample=$2
output_folder=$3
cutoff=$4
script_path_folder=$5

python_script=$script_path_folder/sam_split.py
python $python_script $input_folder/$sample.sam $output_folder $cutoff
echo splitting sample done: $sample
```
