## Supplementary file 3 for "A global view of aging and Alzheimer’s pathogenesis-associated cell population dynamics and molecular signatures in the human and mouse brains": trimming.sh

```
input_folder=$1
sample=$2
output_folder=$3

echo Trimming sample: $sample
trim_galore --paired $input_folder/$sample*R1*.gz $input_folder/$sample*R2*.gz -a2 AAAAAAAA --stringency 3 -o $output_folder
echo Trimming $sample done.
```
