## Supplementary file 1 for "A global view of aging and Alzheimer’s pathogenesis-associated cell population dynamics and molecular signatures in the human and mouse brains"

**EasySci-RNA protocol**

**Abstract**

Single-cell combinatorial indexing ('sci-') is a methodological framework that employs split-pool barcoding to uniquely label the nucleic acid contents of large numbers of single cells or nuclei. Although much progress has been made in making combinatorial indexing methods more efficient, easier to perform, and less costly, there are still major shortcomings in these high-throughput RNA-sequencing techniques. To address this, we employ a new 3-level sci-RNA-seq method (EasySci-RNA) which includes optimizations that drastically improve efficiency, decrease cost per cell sequenced, and increase gene body coverage compared to the previous iteration of the method (sci-RNA-seq3). The new EasySci-RNA method allows for the preparation of 1,000,000 nuclei for under $1,000.

**Protocol workflow**

- Buffer Preparation (Steps 1-12)
- Ligation Primer Annealing (Steps 13-16)
- Tn5 loading (Step 17)
- Nuclei Extraction (Steps 18-26, ~2.5 hours)
- Nuclei Wash (Steps 27-28, ~15-30 minutes,)
- Nuclei Counting (Step 29)
- Reverse Transcription (Steps 30-33, ~1-2.5 hours)
- Pool/Centrifuge/Resuspend/Redistribute (Steps 34-35, 15 minutes)
- Ligation (Steps 36-40, ~2 hours)
- Pool/Centrifuge/Resuspend/Redistribute/Quantify (Steps 41-45, 30 minutes)
- Second-Strand Synthesis (Steps 46-48, ~1.25 hours)
- 0.8x Ampure Beads Purification (Steps 49-55, ~1 hour)
- Tagmentation (Steps 56-57, ~10 minutes)
- SDS Treatment (Steps 58-61, ~1.5 hours)
- PCR (Step 62, ~45 minutes)
- Library Purification (Steps 63-74, ~1 hour)

To validate the experimental setup, it is recommended to start with a species-mixing experiment. We normally start with a mixture of human (HEK 293T) and mouse (NIH/3T3) cells. A good run normally yields single-cell transcriptomes with over 5000 UMIs (with over 20,000 sequencing reads) per cell and >90% purity.

**Required Equipments**

- Bioruptor Sonication Device
- Hemocytometers (Neubauer Improved, Bulldog Bio VWR #102966-632)
- Centrifuge (Eppendorf 5702 RH)
- DynaMag-96 Side Skirted Magnet (Invitrogen, 12027) / DynaMag-96 Side Magnet (Invitrogen, 12331D)
- 12-tube Magnetic Separation Rack (NEB, S1509S)
- Eppendorf Mastercycler (4x)
- Freezer (-20C, -80C) and Refrigerator (4C)
- Gel Imager
- Ice Buckets
- Microscope
- Multi-channel Pipettes (2-20μL, 20-200μL) (Rainin Instruments)
- Pipettors
- 96 well Pipetting System
- Liquid nitrogen tank for sample storage
- FreezeCell Cell Freezing Container (GeneSeeSci, catalog number: 27-802) Eppendorf ThermoMixer C (5382000023) OR Fisherbrand Nutating Mixer (88861043)

**Primer Sequences used**

All primer sequences including RT/Ligation/PCR primers are attached as a separate excel file (EasySci-RNA_primer_sequences.xlsx). All primers are ordered from IDT with standard desalting.

**Materials used**

- Nuclease free water (Ambion, AM 9937)
- 10cm cell culture dish (Genesee, 25-202)
- 6cm cell culture dish (Genesee, 25-260)
- OEMTOOLS 25181 Razor Blades, 100 Pack (VWR, 55411-0055)
- Ward's 40um Sterile Cell Strainer (VWR, 470236-276)
- PluriStrainer Mini 40um (PluriSelect 43-10040-70)
- PluriStrainer Mini 20um (PluriSelect 43-10020-70)
- PluriStrainer Mini 5um (PluriSelect 43-10005-70)
- BD New STERILE , Sealed , 5 ML Syringes Only LUER Lock TIP, No Needle, Disposable (VWR, BD309646)
- Pierce 16% Formaldehyde, Methanol Free (Thermofisher, 28906)
- SUPERase In RNase Inhibitor 20 U/uL (Thermo Fisher Scientific, AM2696)
- BSA 20 mg/ml (NEB, B9000S)
- 1M Tris-HCl (pH 7.5) (Thermo Fisher Scientific, 15567027)
- 5M NaCl (Thermo Fisher Scientific, AM9759)
- 1M MgCl2 (Thermo Fisher Scientific, AM9530G)
- TE Buffer (IDTE, 11-05-01-05)
- Dimethylformamide, 99.8% (Fisher Scientific, AC327175000)
- Dimethyl Sulfoxide (VWR, 97063-136)
- Nuclei Isolation Kit: Nuclei EZ Prep (Millipore Sigma, NUC101-1KT)
- Diethyl Pyrocarbonate (DEPC) (VWR, 97062-652)
- PBS, 1X (Genesee, 25-507)
- Triton X-100 for molecular biology (Sigma Aldrich, 93443-100ML)
- 10mM dNTP (Thermo Fisher Scientific, R0192)
- 192 indexed shortdT primers (100uM, 5′-/5Phos/ACGACGCTCTTCCGATCTNNNNNNNN[10bp barcode]TTTTTTTTTTTTTTTT-3′, where “N” is any base; IDT)
- 192 indexed randomN primers (100uM, 5'-/5Phos/ACGACGCTCTTCCGATCTNNNNNNNN[10bp barcode]NNNNNN-3', where "N" is any base; IDT)
- Maxima H Minus Reverse Transcriptase with Buffer (ThermoFisher, EP0753)
- T4 DNA Ligase (NEB, M0202L)
- EDTA 0.5M Solution (VWR, 97062-656)
- 384 indexed ligation primers (100uM, 5’-AATGATACGGCGACCACCGAGATCTACAC[10bp barcode]ACACTCTTTCCCTAC-3’; IDT)
- Adapter Primer (100uM, 5'-A*G*A*T*C*G*G*A*A*G*A*G*C*G*T*C*G*T*G*T*A*G*G*G*A*A*A*G*A*G*T*G*T*/3ddC/, where '*' represents phosphorothioate bonds between nucleotides and '/3ddC/' represents a dideoxycytidine modification; IDT)
- Elution buffer (Qiagen, 19086)
- NEBNext® Ultra II Non-Directional RNA Second Strand Synthesis Module (NEB, E6111L)
- Tn5 enzyme (in-house produced following Hennig et al. 2018)
- DNA binding buffer (Zymo Research, D4004-1-L)
- AMPure XP beads (Beckman Coulter, A63882)
- SDS, 20% Solution, RNase Free (ThermoFisher AM9820)
- Tween 20 (Millipore Sigma, P9416-100ML)
- Ethanol (Sigma Aldrich, 459844-4L)
- Universal P5 primer (10 μM, 5′-AATGATACGGCGACCACCGAGATCTACAC-3′; IDT)
- 384 indexed P7 primers (10 μM, 5′-CAAGCAGAAGACGGCATACGAGAT[i7]GTCTCGTGGGCTCGG-3′; IDT)
- NEBNext High-Fidelity 2X PCR Master Mix (NEB, M0541L)
- Qubit dsDNA HS kit (Invitrogen, Q32854)
- Qubit tubes (Invitrogen, Q32856)
- E-Gel EX Agarose Gel, 2% (ThermoFisher, G402002)
- E-Gel 50bp DNA Ladder (ThermoFisher, 10488099)
- Falcon Tubes, 15 ml (VWR Scientific, 21008-936)
- Falcon Tubes, 50 ml (VWR Scientific, 21008-940)
- Green pack LTS 200ul filter tips (GP-L200F) (Rainin Instrument, 17002428)
- Pipette Tips RT LTS 20uL FL 960A/10 (Rainin, 30389226)
- Pipette Tips RT LTS 200uL F 960/10 (Rainin, 30389239)
- Pipette Tips RT LTS 200uL FLW 960A/10 (Rainin, 30389241)
- 4-Chip Disposable Hemocytometers, Neubauer Improved, Bulldog Bio (VWR, 102966-632)
- DNA LoBind Tube 1.5 ml, PCR clean (Eppendorf North America, 22431021)
- 1.0mL Self-Standing Cryovial (GeneSeeSci, catalog number: 24-200P)
- LoBind clear, 96-well PCR Plate (Eppendorf North America, 30129512)
- 0.2mL 8-Strip Tubes with Individual Caps (PCR Tubes) (Genesee, 27-125U)
- Reagent reservoirs (Fisher Scientific, 07-200-127)
- Falcon® 5mL Round Bottom w/ Cell Strainer (Fisher Scientific, 352235)
- eXTReme FoilSeal Film (Genesee, 12-156)
- eXTReme Clear Sealing Film (Genesee, 12-157)

**Buffer Preparation**

1. **500mL Nuclei Buffer (Stored in 4 °C)**

10mM Tris-HCl, pH 7.5; 10mM NaCl; 3mM MgCl2 in nuclease free water:

| **Reagent** | **Stock concentration** | **Final concentration** | **Volume (ml)** |
| --- | --- | --- | --- |
| Tris-HCl (pH 7.5) | 1 M | 10 mM | 5 |
| NaCl | 5 M | 10 mM | 1 |
| MgCl2 | 1 M | 3 mM | 1.5 |
| Nuclease-free water | NA | NA | 492.5 |
| Final volume |  |  | 500 |

Filter the buffer through a 0.22uM filter and store the buffer in 4 °C for up to 1 year.

2. **20 mL 10% (volume) Triton-X-100 in nuclease-free water (stored in 4 °C)**

Add 2 mL Triton X-100 to 18 mL nuclease-free water. Mix the solution by pipetting up and down 20 times. The mix can be stored in 4 °C for up to 1 year.

3. **EZ Lysis Buffer + 0.1% RNase Inhibitor (Made fresh each time, stored on ice, 2 mL per tissue sample)**

EZ lysis buffer with 0.1% (volume) SUPERase In RNase Inhibitor . For each sample, combine 2 mL EZ lysis buffer and 2 μL SUPERase In RNase Inhibitor.

4. **EZ Lysis Buffer + 1% DEPC (Made fresh each time, stored on ice, DEPC added just before lysis step, 1 mL per tissue sample)**

EZ Lysis buffer with 1% (volume) DEPC. For each sample, combine 990 μL EZ lysis buffer and 10 μL DEPC

5. **Nuclear Suspension Buffer (NSB) (Made fresh each time, stored on ice)**

Nuclei Buffer with 1% SUPERase In RNase Inhibitor and 1% BSA: For every 1 mL NSB needed, combine 980 μL Nuclei Buffer, 10 μL SUPERase In RNase Inhibitor, and 10 μL BSA.

6. **Nuclear Suspension Buffer + 10% DMSO (NSB + 10% DMSO) (Made fresh each time, 100 μL needed per sample aliquot, stored on ice)**

For every 1 mL needed, add 900 μL Nuclear Buffer and 100 μL DMSO.

7. **Nuclear Suspension Buffer + 0.1% Triton-X-100 (NSB + Triton) (Made fresh each time, 750 μL needed per sample, stored on ice)**

For every 1 mL needed, add 990 μL Nuclei Buffer and 10 μL 10% Triton-X-100.

8. **Nuclear Buffer + 1% BSA + 0.1% Triton-X-100 (NBB) (Made fresh each time, ~8mL needed, store on ice)**

Add 7.84 mL Nuclei Buffer, 80 μL BSA, and 80 μL 10% Triton-X-100.

9. **0.1% Formaldehyde in PBS (Made fresh each time, 1 mL needed per sample, store on ice)**

For every 1 mL solution needed, add 1 mL PBS and 6.25 μL 16% Formaldehyde (Using 1mL glass vial of 16% formaldehyde: open and use a fresh tube of formaldehyde each time)

10. **2x Tagmentation Buffer (Stored in -20 °C)**

Prepare 200 mL of Tagmentation Buffer (filtered):

- 1M Tris HCl (pH 7.5): 4mL
- 1M MgCl2: 2mL
- DMF: 40mL
- H2O: 154mL

Aliquot the solution into 15 mL or 1.5 mL tubes for storage at -20 °C.

11. **1% SDS (Stored at room temperature)**

Mix 1 mL 10% SDS and 9 mL H2O

12. **10% Tween-20 (Stored in 4 °C)**

Mix 1 mL Tween-20 and 9 mL H2O, let sit for 10 minutes before mixing again. Repeat until the solution is homogeneous.

**Ligation Primer Loading (1 hour)**

13. Resuspend and dissolve the Ligation Adaptor Primer Oligo to 100 μM concentration in TE Buffer
14. In each well of an empty 96-well plate, add 5 μL of 100 μM dissolved Ligation Adaptor Primer and 5 μL 100 μM Barcoded Ligation Primers.
15. Anneal the adaptor and ligation primers together by running the following thermocycler program:

- 95 °C for 2 minutes
- Cool to 20 °C at a rate of -1 °C per minute
- Hold at 4 °C

The final annealed concentration will be 50 μM.

16. Dilute the primers to 3.125 μM by adding 150 μL of EB buffer. The resulting product is in stable, double-stranded form and can be stored at 4 °C or frozen. In 4 °C, the annealed primers should be stable for roughly three months and are suitable for short-term testing experiments.

**Tn5 Loading (1 hour)**

17. Protocol Derived from (Hennig et al. 2018), the purified Tn5 protein was also generated following this publication. The Tn5 loading procedure is listed below:

- First mix 150 µL of 100 µM Tn5-ME-B oligo (5’-GTCTCGTGGGCTCGGAGATGTGTATAAGAGACAG-3’, in TE buffer) with 150 µL of 100 µM Tn5-ME-rev oligo (-/5’Phos/CTGTCTCTTATACACATCT-3’, in TE buffer) reaching a final concentration of 50 µM
- Split the mixture into aliquots and perform the following thermocycler conditions: 95 °C for 5 minutes, slowly cooled to 65 °C (0.1C/sec or 2%), 65 °C for 5 minutes, slowly cooled to 4 °C (0.1C/sec or 2%).
- Further dilute the mixture to 35 µM by mixing 10 µL of the oligo mixture with 4.28 µL of TE buffer.
- Combine 1 µL of the Tn5 enzyme at 4 mg/mL with 19 µL of Tn5 Dilution Buffer (25 mM Tris pH 7.5, 800 mM NaCl, 0.1 mM EDTA, 1 mM DTT and 50% glycerol) and 2 µL of the 35 µM Tn5-ME-B/Tn5-MErev oligo mixture.
- Place this solution on a thermomixer at 23 °C for 30 minutes and dilute with 22 µL of glycerol and store at -20 °C for future usage.

Alternatively, use unloaded Tn5 from Diagenode (Catalog. C01070010-10).

**Nuclei Extraction (~2.5 hours for 6 samples)**

18. Cool centrifuge to 4 °C - make sure to use a bucket centrifuge for all centrifuging steps unless otherwise stated, as normal centrifuges may have difficulty making a neat pellet at the bottom of the tube, which is necessary to maximize nuclear recovery.

In a 6 cm dish on ice, cut each tissue section (0.1 g - 0.5 g) into small pieces (< 1 mm^3^) using a razor blade and 1 mL PBS with 10 μL DEPC added. Transfer the tissue and solution into a 1.5 mL tube and spin for 5 minutes at 200g at 4 °C.

NOTES:

- Make sure to add DEPC just before performing lysis, as DEPC has a short half-life in aqueous solutions
- Perform this step in a fume hood, as chopping tissue in a DEPC solution may be toxic
- For larger tissue samples, splitting into multiple 1.5 mL tubes is advised to make pipetting the samples easier
- Ideally, you don't want the tissue sections to thaw until the sections are being cut in the DEPC-PBS solution. To prevent thawing, have a separate container filled with dry ice to place the sections that are currently not being minced with the razor blade
- Generally, we work with a maximum of six tissue sections at one time - you can theoretically process more at the same time, but it may be difficult to manage

19. Dump Supernatant and add 1 mL ice-cold EZ lysis buffer + 1% DEPC to the tissue for nuclei extraction. Pipet the tissue up and down with a 1 mL pipet tip 10 times (cut the top of 1 mL pipet tip if needed for easier pipetting). Incubate on ice for 5 minutes.

NOTES:

- Make sure to add DEPC just before performing lysis, as DEPC has a short half-life in aqueous solutions and will degrade if not added immediately before lysis
- From this point on, use 1 mL pipet tips or wide bore tips when working with nuclei to avoid stress on nuclei

20. Filter tissue with a 40 μm cell strainer into a 6 cm dish and grind tissue on the strainer using a 5 mL syringe plunger. Add 500 μL EZ Lysis Buffer + 0.1% RNase Inhibitor and continue grinding tissue on the strainer. Move solution into a 1.5 mL microcentrifuge tube.

NOTES:

- It is not necessary to push the whole tissue through the filter! Use your own discretion in deciding when to stop grinding the tissue through the filter, but make sure not to tear through the filter!

21. Pellet the nuclei by centrifuging for 5 minutes, 500g at 4 °C. Dump supernatant. Resuspend each tube in 500 μL EZ Lysis Buffer + 0.1% RNase Inhibitor by pipetting up and down three times.
22. Pellet the nuclei by centrifuging for 5 minutes, 500g at 4 °C. Dump supernatant.
23. **Fixation:** Take each tube and add 1 mL of ice-cold 0.1% Formaldehyde suspended in PBS. Start a 10-minute timer immediately after formaldehyde is added. Mix up and down to resuspend the pellet.

For multiple samples, add 1 mL directly to the top of tubes without changing tips and without touching the tubes; start timer once the formaldehyde is added to all tubes. Once done, go back and pipet up and down the solution in each sample to resuspend the pellet, making sure to switch tips for each sample.

NOTES:

- Perform this step in a fume hood as formaldehyde is toxic

24. Pellet the nuclei immediately afterward by centrifuging for 3 minutes, 500g at 4 °C. Dump supernatant in a chemical waste container. Resuspend each tube in 500 μL EZ Lysis Buffer + 0.1% RNase Inhibitor by pipetting up and down three times.
25. Pellet the nuclei by centrifuging for 5 minutes, 500g at 4 °C. Dump supernatant. Resuspend each tube in 500 μL EZ Lysis Buffer + 0.1% RNase Inhibitor by pipetting up and down three times.
26. PERFORM THIS STEP IF YOU WISH TO STORE NUCLEI FOR LATER USE - OTHERWISE, SKIP TO THE SECOND PART OF THE NEXT STEP:

Pellet the nuclei by centrifuging for 5 minutes, 500g at 4 °C. Resuspend each tube in 100-500 μL NSB + 10% DMSO and split into 100 μL aliquots. Slow freeze in a -80 °C freezer and keep for storage. Optimally, use specialized slow-freezing chambers with 1.0 mL Self-Standing Cryovials (FreezeCell Cell Freezing Container, GeneSeeSci, catalog number: 27-802) (1.0 mL Self-Standing Cryovial, GeneSeeSci, catalog number: 24-200P) **(STOP POINT)**.

**Nuclei Wash (~15-30 minutes for 6-30 samples)**

27. 1) PERFORM BELOW IF YOU ARE WORKING WITH PREVIOUSLY FROZEN, STORED NUCLEI:

Thaw cells for 30 seconds in a 37 °C water bath. Add 400 μL NSB + Triton to each sample to resuspend pellets, and then sonicate for 12 seconds at low power. After the sonication, filter nuclei through a 20 μm filter. Wash the filter with an additional 250 μL NSB + Triton and then pellet the nuclei for 5 minutes, 500g at 4 °C.

2) PERFORM BELOW IF YOU ARE DIRECTLY CONTINUING FROM NUCLEI EXTRACTION:

Add 500 μL NSB + Triton to each sample to resuspend pellets, and then sonicate for 12 seconds at low power. After, filter nuclei through a 20 μm filter. Wash the filter with an additional 250 μL NSB + Triton and then pellet the nuclei for 5 minutes, 500g at 4 °C.

28. Resuspend the pellet in 100 μL of NSB.

**Nuclei Counting**

29. Count the nuclei concentration for each sample.

Optimally, use a buffer with DAPI and a fluorescent microscope to distinguish between actual nuclei and debris.

Dissolve 10 mg DAPI in 2 ml of deionized water (dH2O) with a final concentration of 5 mg/ml. Split the DAPI solution into multiple tubes (100 ul per tube).

Take out one tube (100 ul, 5 mg/ml DAPI), add 1.9 ml deionized water (dH2O). Split the diluted DAPI solution into multiple tubes (100 ul per tube, 0.25 mg/ml DAPI).

Store the DAPI solution in the -20 °C freezer.

Make the DAPI counting solution: in 500 μL of Nuclei Buffer, add 0.5 μL - 1μL of 0.25mg/mL DAPI solution.

Take 1 μL of the sample and combine it with 9 uL of the counting solution. Mix the solution and take 6 μL to dispense into a hemocytometer for counting the concentration of nuclei.

**Reverse Transcription (~1 - 2.5 hours depending on number of samples)**

30. For each well of 2 x 96 well plates, add a maximum of 20,000 nuclei in 4 μL of NSB; also add 0.5 μL of 10 mM dNTP.

NOTES:

- Nuclei generally distributed into PCR strips and then distributed into wells - make sure not to pipet up and down to avoid introducing extra stress to the nuclei
- It is recommended to use wide bore multichannel tips for mixing the nuclei

31. Add 1 μL of 50 μM short-dT primer and 1 μL of 50 μM randomN primer into each well using multi-channel pipettes or 96-well liquid handler. Incubate plates at 55 °C for 5 minutes. Immediately place plates on ice afterward.
32. Prepare the reverse transcription reaction mix by combining:

- 5X Maxima Buffer: 420μL
- Maxima Reverse Transcriptase: 105μL
- SUPERase In RNase Inhibitor: 105μL
- Nuclease Free H2O: 105μL

Add 3.5μL to each well for each of the plates; pipet up and down only once

33. Start the reverse transcription with the following thermocycler program:

- 4 °C for 2 minutes
- 10 °C for 2 minutes
- 20 °C for 2 minutes
- 30 °C for 2 minutes
- 40 °C for 2 minutes
- 50 °C for 2 minutes
- 55 °C for 15 minutes

**Pool/Centrifuge/Resuspend/Redistribute (15 minutes)**

34. Add 10 μL NBB into each well, pool solution, and move solution into a 15 mL tube. Centrifuge the tube for 3 minutes, 1000g at 4 °C.
35. Use a pipet to aspirate supernatant. Resuspend nuclei in 1 mL NBB and then transfer the nuclei into a 1.5 mL microcentrifuge tube. Centrifuge the tube for 3 minutes, 1000g at 4 °C to pellet the nuclei.

**Ligation (1 hour)**

36. Dump the supernatant. Resuspend the cells in 950 μL NBB. Distribute the nuclei into four PCR plates, with 2.5 μL of the solution going into each well.
37. To each well, add 1 μL of the appropriate DNA ligation primer/adaptor complex (3.125 μM).
38. Create a mixture of:

- 210 μL 10X T4 Ligation Buffer
- 21 μL SUPERase In RNase Inhibitor
- 210 μL T4 DNA Ligase
- 189 μL Nuclease Free Water

Add 1.5 μL of the mixture to each of the PCR plate wells.

39. Incubate plates for 30 minutes at room temperature with gentle shaking (300 rpm with Thermomixer, 50 rpm on Fisherbrand Nutating Mixer).
40. Add 1 μL EDTA (18 mM diluted in nuclease free water) into each well and pool all solution into a 15 mL tube.

**Pool/Centrifuge/Resuspend/Redistribute/Quantify (30 minutes)**

41. Centrifuge the tube for 3 minutes, 1000g at 4 °C. Pipet out the supernatant.
42. Resuspend the nuclei in 1 mL NBB. Transfer the cells into a microcentrifuge tube. Centrifuge the tube for 3 minutes, 1000g at 4 °C. Dump the supernatant.
43. Resuspend the nuclei in 500 μL NBB. Filter the nuclei using a 40 μM filter and then wash the filter with an additional 250 μL NBB. Centrifuge the tube for 3 minutes, 1000g at 4 °C. Dump the supernatant.
44. Resuspend the nuclei in 500 μL NBB for nuclei counting - it is recommended to use a fluorescent microscope with a solution with DAPI to distinguish nuclei from debris.
45. Distribute the nuclei into a 96 well plate with 10,000 nuclei per well, suspended in 4 μL total volume (final concentration = 2,500 nuclei/μL).

NOTES:

- You can directly freeze and store cells at this point, but it is recommended to proceed directly to second-strand synthesis as dsDNA should be more stable in storage compared to ssDNA
- If you choose to freeze the plate, it is okay to place it directly in -80 °C freezer without flash-freezing
- You can also store nuclei directly into PCR strips if you don't need to profile a whole plate of cells

**Second-Strand Synthesis (1 hour 15 minutes)**

46. Thaw Second-Strand Synthesis buffer in room temperature.
47. Prepare Second-Strand Synthesis mix: for each well, add ⅔ μL Second-Strand Synthesis buffer + ⅓ μL Second-Strand Synthesis Enzyme Mix.
48. Perform Second-Strand Synthesis in Thermocycler: incubate samples at 16 °C for one hour. **(STOP POINT).**

**0.8x Ampure Beads Purification (~1 hour for one plate)**

49. Take one plate of prepared cells after Second-Strand Synthesis and add 5 μL DNA binding buffer to each well, mix, and let the resulting solution sit for 5 minutes at room temperature.

NOTES:

- You can also perform this protocol with PCR strips if you do not need to profile a whole plate

50. Add 8 μL ampure beads to each well, mix well via pipetting, and let the resulting solution sit for 5 minutes at room temperature.
51. Place the solution on a magnetic rack and let the solution sit for 5 minutes.
52. Remove the resulting supernatant and add 50 μL of 80% ethanol (do not mix up and down). Remove the ethanol.
53. Wash one more time with 50 μL of 80% ethanol (do not mix up and down). Remove the ethanol, centrifuge the pellet down, place the plate on the magnetic rack, and remove the remaining residual ethanol.
54. Take the plate off of the magnetic rack and elute the beads in 7.6 μL of elution buffer. Incubate the solution for three minutes at room temperature.
55. Place the plate back on the magnetic rack and let the plate sit for three minutes at room temperature. Aspirate 6.6 μL of solution without touching the magnetic beads and transfer the solution into a new plate.

**Tagmentation (10 minutes)**

56. Prepare a mixture of 1:100 Tagmentase:Tagmentation Buffer mix. Add 6.6 μL of the mix to each well and pipet up and down to mix.
57. Incubate plate in the thermocycler at 55 °C for 5 minutes. Place on ice immediately following the reaction.

**SDS Treatment (45 minutes)**

58. For each well, add a mixture of:

- 0.4 μL 1% SDS
- 0.4 μL BSA
- 2 μL 10 μM Universal P5 Primer

59. Incubate the plate at 55 °C for 15 minutes. Place the plate on ice immediately following the reaction.
60. Add 2 μL 10% Tween-20 to each well.
61. Add 2 μL Indexed P7 primer to each well. Centrifuge the plate after this step.

**PCR (45 minutes)**

62. Add 20 μL NEBNext Master Mix into each well and pipet up and down. Place samples into a thermocycler and run the following reaction:

- 72 °C for 5 minutes
- 98 °C for 30 seconds
- 12-15 cycles of 98 °C for 10 seconds, 66 °C for 30 seconds, 72 °C for 30 seconds
- 72 °C for 5 minutes

NOTES:

- It may be helpful to run a qPCR to determine the optimal number of cycles for amplification
- You can store the resulting PCR products in -20 °C **(STOP POINT)**.

**Library Purification (1 hour)**

63. Pool all the wells together. Take 200 μL of the PCR product and perform a 0.8x ampure beads purification: start with adding 160 μL beads to the 200 μL of solution. Mix the solution via vortexing and let the resulting solution sit at room temperature for 5 minutes.
64. Place the solution on a magnetic rack and let the solution sit for 5 minutes until the beads are clearly separately from the solution.
65. Aspirate and remove the solution, making sure not to touch the beads. Add 1 mL of 80% ethanol to rinse beads and then remove the ethanol.
66. Add 1 mL of 80% ethanol for a second wash and then remove the ethanol.
67. Mix the bead with 105 μL of elution buffer followed by vortexing. Let the solution sit at room temperature for 3 minutes.
68. Place the solution on the magnetic rack after a brief centrifuge. Let the solution sit at room temperature for 3 minutes.
69. Transfer 100 μL of the solution into a new tube.
70. Quantify the library concentration and visualize the library via electrophoresis (performed using a Qubit and a 2% Agarose E-Gel). An example library is shown below:

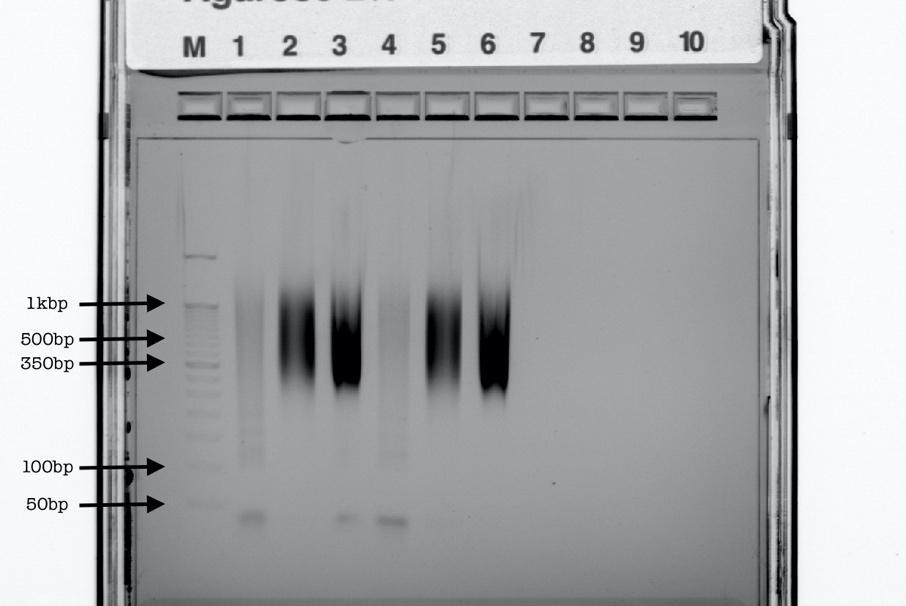

- Column M: 50 bp ladder
- Column 1: PCR product for the first 96-well plate, no purifications
- Column 2: The library after one round of 0.8x beads purification

71. Sequence the library on the Novaseq Platform (Read1:100 bp, Read2: 100 bp, Index 1: 10 bp, Index 2: 10 bp).

**References**

Hennig, Bianca P., Lars Velten, Ines Racke, Chelsea Szu Tu, Matthias Thoms, Vladimir Rybin, Hüseyin Besir, Kim Remans, and Lars M. Steinmetz. 2018. “Large-Scale Low-Cost NGS Library Preparation Using a Robust Tn5 Purification and Tagmentation Protocol.” *G3*  8 (1): 79–89.
